# Network toxicology focused investigation on the impacts of inorganic arsenic and cadmium on human and ecosystem health

**DOI:** 10.1101/2024.11.18.624172

**Authors:** Nikhil Chivukula, Shreyes Rajan Madgaonkar, Kundhanathan Ramesh, Swetha Mangot, Panneerselvam Karthikeyan, Shambanagouda Rudragouda Marigoudar, Krishna Venkatarama Sharma, Areejit Samal

## Abstract

Heavy metals like arsenic and cadmium are persistent environmental pollutants that pose serious health risks to humans and ecosystems due to their toxicity, bioaccumulation potential, and frequent presence in consumer products. Network toxicology offers a holistic *in silico* framework to elucidate the complex biological mechanisms of toxicity, thereby supporting New Approach Methodologies (NAMs) for toxicity assessment. In this study, network toxicological tools were utilized to investigate arsenic- and cadmium-induced toxicities. Toxicity endpoints associated with inorganic arsenic and cadmium compounds were curated from six exposome-relevant databases and mapped to key events (KEs) across adverse outcome pathways (AOPs) cataloged in AOP-Wiki. This led to construction of stressor-AOP networks, revealing 51 AOPs associated with arsenic and 78 with cadmium, and facilitated mechanistic case studies of pathways relevant to human and ecological health. Toxicity concentrations and bioconcentration factors from the ECOTOX database were then used to construct stressor-species networks, that helped identify species that are vulnerable and potentially bioaccumalate these chemicals. Further, the construction of species sensitivity distributions (SSDs) and toxicity-normalized SSDs (SSDn), provided a comparative framework for prioritizing these compounds in risk assessments. Further, integrating SSD data with stressor-species networks identified species groups particularly sensitive to arsenic and cadmium exposure, enhancing these networks’ utility for ecological risk assessment. The networks and related data generated in this study are freely available for further research at https://cb.imsc.res.in/heavymetaltox/. Overall, this study offers a comprehensive perspective on the toxicological impact of inorganic arsenic and cadmium compounds, supporting a One Health approach to their regulatory and mitigation strategies.

## 1. Introduction

Heavy metals are persistent and highly toxic pollutants in the environment due to their non-biodegradable nature and capacity to bioaccumulate in living organisms [1,2]. These pollutants are particularly concerning as they are released into the environment through various anthropogenic activities, where they can biomagnify through the food chain, eventually causing severe health and ecological impacts [1–3]. Among the heavy metals, arsenic, cadmium, and their compounds have been classified as Group 1 carcinogens by the International Agency for Research on Cancer (IARC) [4], indicating their well-established carcinogenicity in humans. The United States Agency for Toxic Substances and Disease Registry (ATSDR) has categorized these two metals as priority substances based on their frequency of occurrence, associated toxicity, and potential for human exposure [5]. In particular, inorganic forms of arsenic and cadmium are hazardous due to their higher rates of absorption, toxic valence states, and enhanced ability to cause cellular and organ-level damage [1,2,6,7]. Moreover, these compounds are readily discharged into water bodies through activities such as smelting, mining, and fertilizer use, where they accumulate in sediments and infiltrate food webs, posing threats to aquatic life and marine ecosystem health [8–12]. Therefore, a comprehensive understanding of their risks is essential to investigate their impacts on human and ecosystem health.

Network toxicology is an emerging interdisciplinary field that integrates toxicology, network biology, computational science, and environmental sciences to study the impact of toxic substances on biological systems [13]. In the context of New Approach Methodologies (NAMs), it offers a novel approach integrating diverse biological data within the network framework, offering a holistic understanding of stressor-induced toxicity and elucidating underlying toxicity mechanisms [13,14]. Adverse Outcome Pathway (AOP) framework is one such network toxicology framework to understand mechanisms underlying stressor-induced toxicities [15–17]. An AOP encapsulates the toxicity mechanism as a linear sequence of causally linked biological events, starting from a molecular initiating event and culminating in an adverse outcome of regulatory significance [15–17]. AOP networks, constructed by linking multiple AOPs through shared biological events, provide a comprehensive understanding of complex interactions and cumulative risks from various stressors, enhancing predictive toxicology, risk assessment, and regulatory decision-making [18–20]. Although AOP networks have been used to explore arsenic-induced toxicities such as male reproductive disorders [21], and cadmium-induced toxicities such as preeclampsia in humans and early life-stage mortality in aquatic organisms [22], these efforts have predominantly relied on mammalian-centric datasets. Recently, Sahoo *et al.* [23] leveraged the ECOTOX database developed by the United States Environmental Protection Agency (US EPA), to explore ecologically relevant toxicity mechanisms underlying petroleum hydrocarbons-induced toxicity through the AOP framework. Therefore, there is a need for a more comprehensive approach that integrates mammalian and ecologically relevant datasets to expand our understanding of the toxicities induced by arsenic and cadmium across both human and ecological contexts.

The stressor-species network is a network toxicology framework for examining the complex interactions between multiple environmental stressors and various species within ecosystems [23–26]. These networks help identify the impacts of various stressors on species, elucidating potential risks to ecosystems and aiding in the prioritization of regulatory measures [23–26]. Previously, Sahoo *et al.* [23] leveraged the diverse toxicity data available in the ECOTOX database to construct stressor-species networks to investigate petroleum hydrocarbons-induced toxicities in various ecologically relevant species. The networks built using toxicity data revealed the range of species affected by petroleum hydrocarbon-associated toxicity, while those constructed using bioconcentration factor (BCF) data highlighted the potential for these chemicals to accumulate within different species [23]. Furthermore, aquatic toxicity data within ECOTOX was leveraged to develop species sensitivity distributions (SSDs), enabling the derivation of hazard concentrations for the ecological risk assessment of these chemicals [23]. Thus, the integration of arsenic- and cadmium-associated ecotoxicological data from the ECOTOX database with stressor-species networks can provide valuable insights into their impacts on biodiversity and inform more effective risk management strategies.

In this study, the inorganic arsenic and cadmium compounds were identified from six exposome-relevant resources, and subsequently their corresponding toxicological endpoints were extracted. This endpoint data was systematically integrated within the AOP framework to achieve large-scale coverage of inorganic arsenic- and cadmium-induced toxicities affecting both human and ecosystem health. Furthermore, AOP networks, based on the associated AOPs, were constructed to explore diverse toxicity mechanisms. Thereafter, key toxicity pathways from these networks were identified, and their relevance to inorganic arsenic- and cadmium-induced toxicities was supported through existing literature. In addition, acute and chronic toxicity data, and bioconcentration factor data from the ECOTOX database was used to construct stressor-species networks, to enable the identification of the ecological effects of inorganic arsenic and cadmium compounds. Finally, the aquatic toxicity information in the ECOTOX database was used to construct species sensitivity distributions (SSDs), and the potential harmful concentrations of these chemicals in aquatic environments were estimated. In sum, this is among the first few studies to utilize diverse network toxicology approaches to explore the impacts of inorganic arsenic and cadmium on both human and ecosystem health.

## 2. Methods

### 2.1. Compilation and curation of inorganic compounds of arsenic and cadmium

The aim of this study is to gain a comprehensive understanding of the toxicological effects of inorganic arsenic and inorganic cadmium on both human and ecosystem health through the Adverse Outcome Pathway (AOP) framework. To achieve this, six exposome-relevant databases, namely ToxCast [27], Comparative Toxicogenomics Database (CTD) [28,29], DEDuCT [30–32], NeurotoxKb [33,34], AOP-Wiki [35], and ECOTOX [36,37], were used to identify arsenic and cadmium compounds with toxicologically-relevant biological endpoints in humans and ecosystems. Next, chemical information was extracted and the compounds were mapped to standardized PubChem [38] and Chemical Abstracts Service (CAS) [39] identifiers, to ensure a non-redundant chemical list. Subsequently, two-dimensional (2D) structural data was retrieved from PubChem and Open Babel [40] was used to generate key structural representations, including SMILES, InChI, and InChIKey. ClassyFire [41,42] was used to classify these chemicals as either inorganic or organic. The compounds that could not be classified by ClassyFire, were manually verified and classified based on their structural definitions [41]. Through this extensive systematic process, 62 inorganic arsenic compounds and 18 inorganic cadmium compounds (Table S1) were identified, and were used for subsequent analyses in this study.

### 2.2. Compilation and curation of ‘high confidence’ AOPs within AOP-Wiki

The AOP framework organizes the biological events (key events (KEs)) underlying a stressor-induced perturbation into a directional network, beginning with a molecular interaction (molecular initiating event (MIE)) and culminating in an adverse outcome (AO) relevant for regulation [15]. In this directed network, KEs function as vertices, with the MIE as the source vertex and the AO as the terminal vertex, while the connections between them, known as Key Event Relationships (KERs), serve as the edges, collectively forming AOPs [15]. These AOPs are developed globally, targeting diverse adverse outcomes, and are deposited into AOP-Wiki [35], the largest global collaborative platform for AOP development and associated scientific evidence. As AOP information is continuously updated by contributors based on new evidence, AOP-Wiki is considered a living document [43]. To ensure we obtained the most up-to-date information, a method developed in previous works [22,44] was followed to extract and filter data from AOP-Wiki, and identify non-empty, complete, and high quality AOPs, which are designated as ‘high confidence AOPs’ (Supplementary Information). Briefly, the flat file dated April 1, 2024, available for download on AOP-Wiki was accessed, relevant information was extracted using an in-house Python script, and 342 high confidence AOPs (Figure S1; Table S2) were identified through a systematic manual and computational effort (Supplementary Information). These 342 high confidence AOPs comprised 1163 unique KEs (Table S3) and 1827 unique KERs (Table S4).

### 2.3. Construction of stressor-AOP network for arsenic and cadmium

In this study, a stressor-AOP network was constructed by integrating multiple stressor-AOP associations to provide a broader perspective on the impacts of inorganic arsenic and cadmium on biological pathways and adverse outcomes [44,45]. Building on previous work [22,23,44], data from ToxCast [27], CTD [28,29], DEDuCT [30–32], NeurotoxKb [33,34], and ECOTOX [36,37], was used to obtain biological endpoints associated with inorganic arsenic and cadmium compounds in both humans and ecologically relevant species (Supplementary Information). Next, these endpoints were manually mapped to KEs within AOP-Wiki, and resultant chemical-KE mappings were grouped under arsenic and cadmium, respectively (Table S5; Supplementary Information). Finally, leveraging these arsenic-KE and cadmium-KE associations a bipartite stressor-AOP network was constructed, linking arsenic and cadmium to 310 and 318 high-confidence AOPs respectively, with varying levels of relevance and coverage scores (Table S6; Supplementary Information). The resultant networks were visualized using Cytoscape [46].

### 2.4. Construction of stressor-species networks by utilizing ECOTOX data

The stressor-species network provides a framework to understand how multiple environmental stressors interact with various species, offering insights into their effects on ecosystems and biodiversity [23–25]. ECOTOX, one of the largest openly accessible ecotoxicological resources, compiles manually curated toxicological data for over 12,000 chemicals tested on more than 13,000 ecologically relevant species [36]. In this study, stressor-species networks were constructed for the inorganic compounds of arsenic and cadmium by leveraging ECOTOX data on chemical concentrations associated with acute toxicity (LC_50_ or EC_50_), chronic toxicity (NOEC) and bioconcentration factors (BCF) to gain a comprehensive understanding of their toxicities and accumulation potential in different species. Based on Sahoo *et al.* [23], the acute toxicity concentrations were normalized to ppm equivalents (mg/L or mg/kg) by relying on conversion factors (Table S7). Then, for each stressor-species pair, studies with durations between 24 and 96 hours were considered, and for pairs with multiple concentration values, their geometric mean was computed. Thereafter, a network was constructed linking 15 inorganic arsenic and 8 inorganic cadmium compounds to 167 and 644 species, respectively (Table S8). Similarly, for chronic toxicity values, the concentrations were normalized to ppm equivalents, geometric mean was computed in case of more than one concentration for a stressor-species pair, and a network was constructed linking 16 arsenic and 4 cadmium compounds to 130 and 264 species respectively (Table S8). Similarly, BCF values were normalized, and the highest BCF value per stressor-species pair was selected, and a network was constructed linking 5 arsenic and 4 cadmium compounds to 32 and 147 species, respectively (Table S9). The resultant networks were visualized using Cytoscape [46].

### 2.5. Construction of Species Sensitivity Distributions for arsenic and cadmium by utilizing ECOTOX data

Species Sensitivity Distribution (SSD) is a widely used statistical method in ecological risk assessment to estimate species’ sensitivities to environmental stressors, and derive hazard concentration affecting 5% of the species (HC05), which informs regulatory guidelines for biodiversity protection [47–50]. Based on Sahoo *et al.* [23], SSDs were constructed using acute toxicity data (LC_50_ and EC_50_) and chronic toxicity data (NOEC) from ECOTOX for inorganic arsenic and cadmium compounds in aquatic environments (Supplementary Information). Briefly, the acute and chronic toxicity data for each chemical were obtained from the stressor-species network, the data was filtered to identify species with aquatic habitats, and only chemicals with data across at least five ECOTOX species groups were included to ensure robust SSDs (Table S10-S11). We employed the US EPA’s SSD Toolbox [51] and R-based ssdtools package [52] to construct SSDs and calculate corresponding HC05 values (Supplementary Information). Further, to identify sensitive species, the stressor-species links, within the stressor-species network constructed based on toxicity values, were annotated by comparing them with corresponding HC05 values.

Furthermore, toxicity-normalized SSDs (SSDn) were constructed to assess overall toxicity of inorganic arsenic and cadmium compounds [53,54]. The SSDn approach refines ecological risk assessment by normalizing chemical toxicity data based on a sensitive reference species, thereby enhancing the comparison of chemicals that share toxicological similarities [53,54]. Lambert *et al.* [54] suggested the following criterion to choose an appropriate reference species (referred to as ‘nSpecies’) for the construction of SSDn:

- All chemicals of interest must have toxicity data for the nSpecies
- When there are multiple potential nSpecies, the most sensitive species should be used for normalization
- When possible, normalize toxicity values to the nSpecies with the most similar sensitivity to all compounds (i.e., smallest range of percentiles in single-chemical SSDs)

In the case of arsenic, *Daphnia magna* was the only species with acute toxicity data across the four inorganic arsenic compounds (Table S10). Therefore, *Daphnia magna* was chosen as the nSpecies for acute toxicity data-based SSDn computation for arsenic. It was observed that 6 species namely, *Cyprinus carpio, Danio rerio, Daphnia magna, Oncorhynchus mykiss, Rhinella arenarum,* and *Terapon jarbua* had chronic toxicity data for all three inorganic arsenic compounds (Table S11). Among these, the chronic toxicity data of *Terapon jarbua* satisfied the other two criterion outlined by Lambert *et al.* [54], and therefore *Terapon jarbua* was chosen as nSpecies for chronic toxicity data-based SSDn computation for arsenic.

In the case of cadmium, it was observed that 10 species namely, *Artemia salina*, *Asellus aquaticus*, *Brachionus calyciflorus*, *Danio rerio*, *Daphnia magna*, *Daphnia pulex*, *Gambusia affinis*, *Hyalella azteca*, *Oncorhynchus mykiss*, and *Pimephales promelas* had acute toxicity data for all the four inorganic cadmium compounds (Table S10). Among these, the toxicity data of *Hyalella azteca* satisfied the other two criterion outlined by Lambert *et al.* [54], and therefore *Hyalella azteca* was chosen as nSpecies for cadmium. It was observed that 2 species namely, *Cyprinus carpio,* and *Oncorhynchus mykiss* had chronic toxicity data for all four inorganic cadmium compounds (Table S11). Among these, the chronic toxicity data of *Oncorhynchus mykiss* satisfied the other two criterion outlined by Lambert *et al.* [54], and therefore *Oncorhynchus mykiss* was chosen as nSpecies for chronic toxicity data-based SSDn computation for cadmium.

The toxicity data was normalized with respect to that of nSpecies for the identified inorganic compounds of arsenic and cadmium, and were designated as nSpecies equivalents. Then, for each species, the geometric mean of the nSpecies equivalents across different chemicals were calculated to obtain the corresponding species mean toxicity values (SMTVs) (Table S12-S13). This computed SMTV data was used to construct arsenic-specific SSDn and cadmium-specific SSDn by following the procedure used in SSD construction, and their corresponding toxicity-normalized hazard concentration (HC05n) values were computed. Then, the chemical specific HC05 values were estimated from the corresponding HC05n using the following formula suggested by Lambert *et al.* [54]:

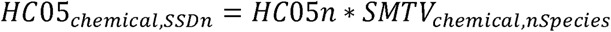

Here, SMTV_chemical,nSpecies_ refers to the SMTV for the nSpecies identified for a chemical, and HC05_chemical,SSDn_ represents the derived HC05 value for that chemical from the corresponding HC05n value. Additionally, the fold change was calculated between HC05_chemical,SSDn_ and the corresponding HC05 value obtained from single-chemical SSDs to assess the accuracy of the values computed from SSDn approach. Here, the following formula was used to compute the fold change:

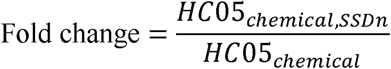

where, HC05_chemical_ represents the HC05 value calculated from the single-chemical SSD for the corresponding chemical. As per Lambert *et al.* [54], values falling within a 5-fold range are considered ‘Good’.

## 3. Results and Discussion

### 3.1. Overview of the curated list of inorganic arsenic and cadmium compounds

Arsenic and cadmium are highly toxic heavy metals that are widely present in the environment [1,2]. The inorganic forms of these metals are particularly hazardous due to their higher absorption rates, multiple valence states, and their potential to cause damage at both cellular and organ levels [1,2,6,7]. In this study, network toxicology based approaches were utilized to assess the adverse effects of these heavy metals on human and ecosystem health. 62 inorganic arsenic and 18 inorganic cadmium compounds were identified from six exposome-relevant databases (Methods; Table S1), of which many compounds are prohibited from usage in the European Union [55], with some listed as high concern chemicals under REACH regulations [56] (Table S1). Notably, some of these compounds have been detected in human biospecimens, including the placenta, urinary bladder, brain, skin, bone, kidney, and pancreas [57,58] (Table S1). Furthermore, the US EPA’s Chemicals and Products Database (CPDat) [59] records the presence of some of these compounds in consumer products such as batteries, electrical appliances, and construction materials (Table S1). Finally, for these compounds, stressor-AOP network was constructed to identify the underlying toxicity mechanisms, stressor-species networks were constructed to explore their effects on ecosystems, and Species Sensitivity Distributions (SSDs) were computed based on ECOTOX data to support their ecological risk assessments (Figure 1). Notably, the analysis and results reported in this study have been made available for further research through the associated website: https://cb.imsc.res.in/heavymetaltox/.

**Figure 1:**
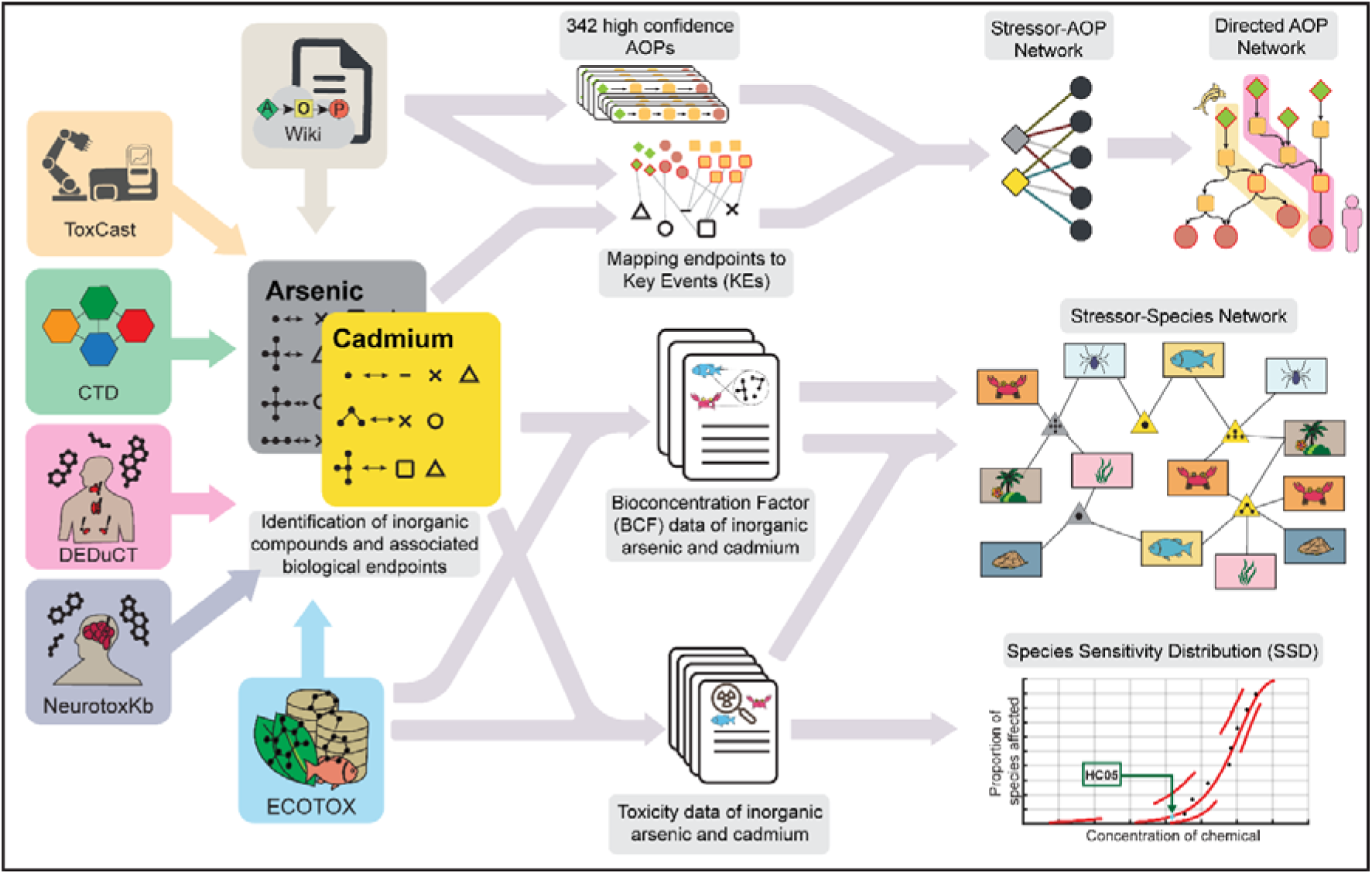
Summary of the workflow followed to identify inorganic arsenic and cadmium compounds from six exposome-relevant resources, followed by the exploration of their impacts on human and ecosystem health through stressor-Adverse Outcome Pathway networks, stressor-species networks and Species Sensitivity Distributions (SSDs) based on their toxicity values.

### 3.2. Stressor-AOP network for inorganic arsenic and cadmium

A stressor-AOP network linking inorganic arsenic and cadmium to their associated high-confidence AOPs was constructed through a systematic data integration approach (Methods; Supplementary Information; Table S6). 51 AOPs linked to arsenic (denoted as arsenic-AOPs), and 78 AOPs to cadmium (denoted as cadmium-AOPs), with Level 5 relevance and coverage score (CS) ≥ 0.4 (Table S6), were identified. From these, undirected AOP networks were constructed for arsenic and cadmium, revealing 48 arsenic-AOPs (Figure 2; Table S14) and 72 cadmium-AOPs (Figure 3; Table S15) as comprising the largest connected components, respectively. Notably, the higher number of cadmium-AOPs in this study, compared to Sahoo *et al.* [22], is a consequence of the expanded coverage of ecotoxicologically-relevant AOPs identified through the ECOTOX database. Then, directed AOP networks corresponding to the largest connected components were constructed for the arsenic-AOPs (comprising 152 unique KEs and 243 unique KERs) (Figure S2-S4; Table S16) and cadmium-AOPs (comprising 263 unique KEs and 422 unique KERs) (Figure S5-S7; Table S17) to gain deeper insights into their respective toxicity mechanisms. Further, to explore various network features, different node-centric network measures were computed for the directed arsenic-AOP and cadmium-AOP networks.

**Figure 2:**
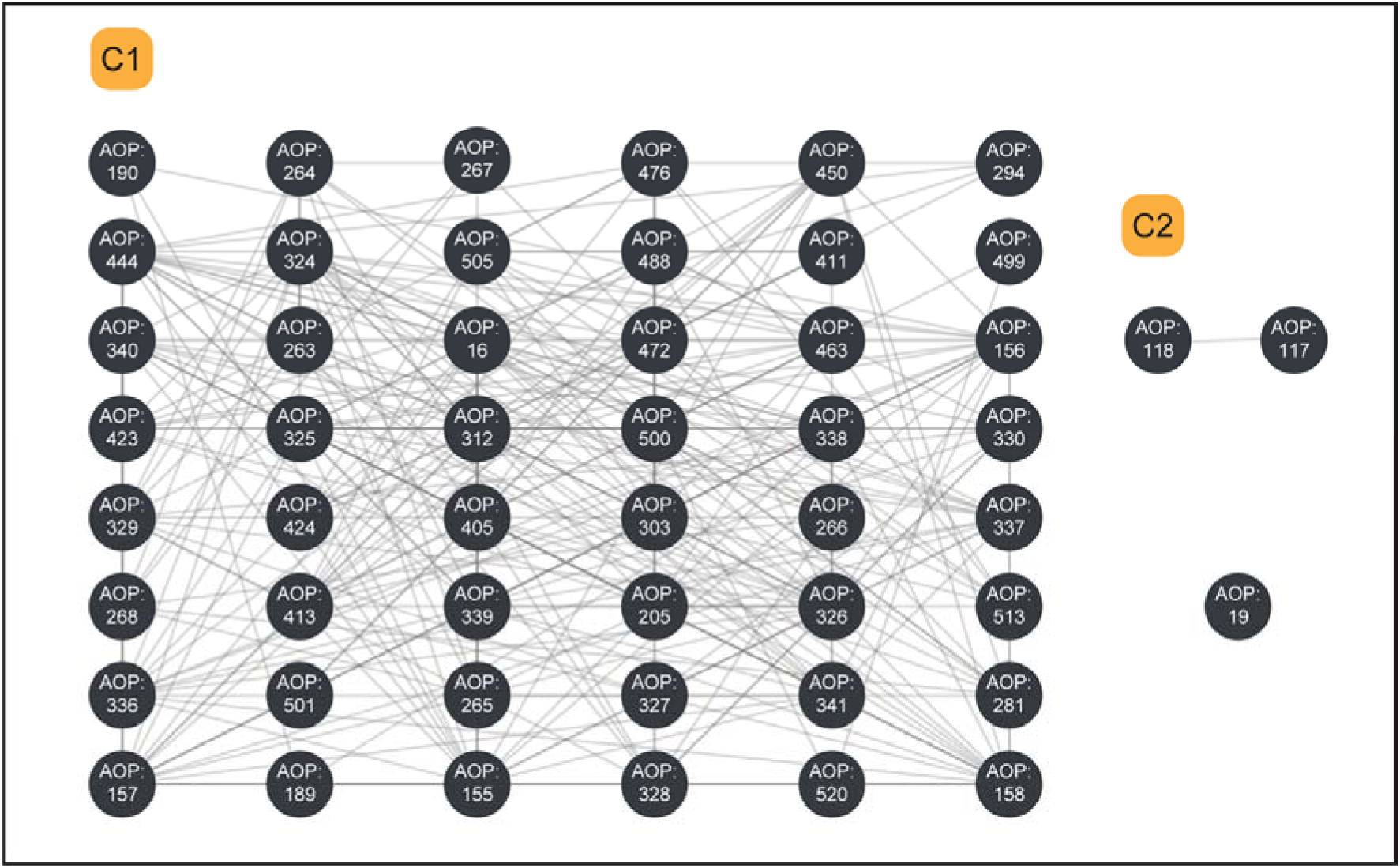
Undirected network of arsenic–AOPs. Each node corresponds to an arsenic–AOP, and an edge between two nodes denotes that the two AOPs share at least one KE. This undirected network has 2 connected components (with two or more nodes) which are labeled as C1 and C2, and 1 isolated node.

**Figure 3:**
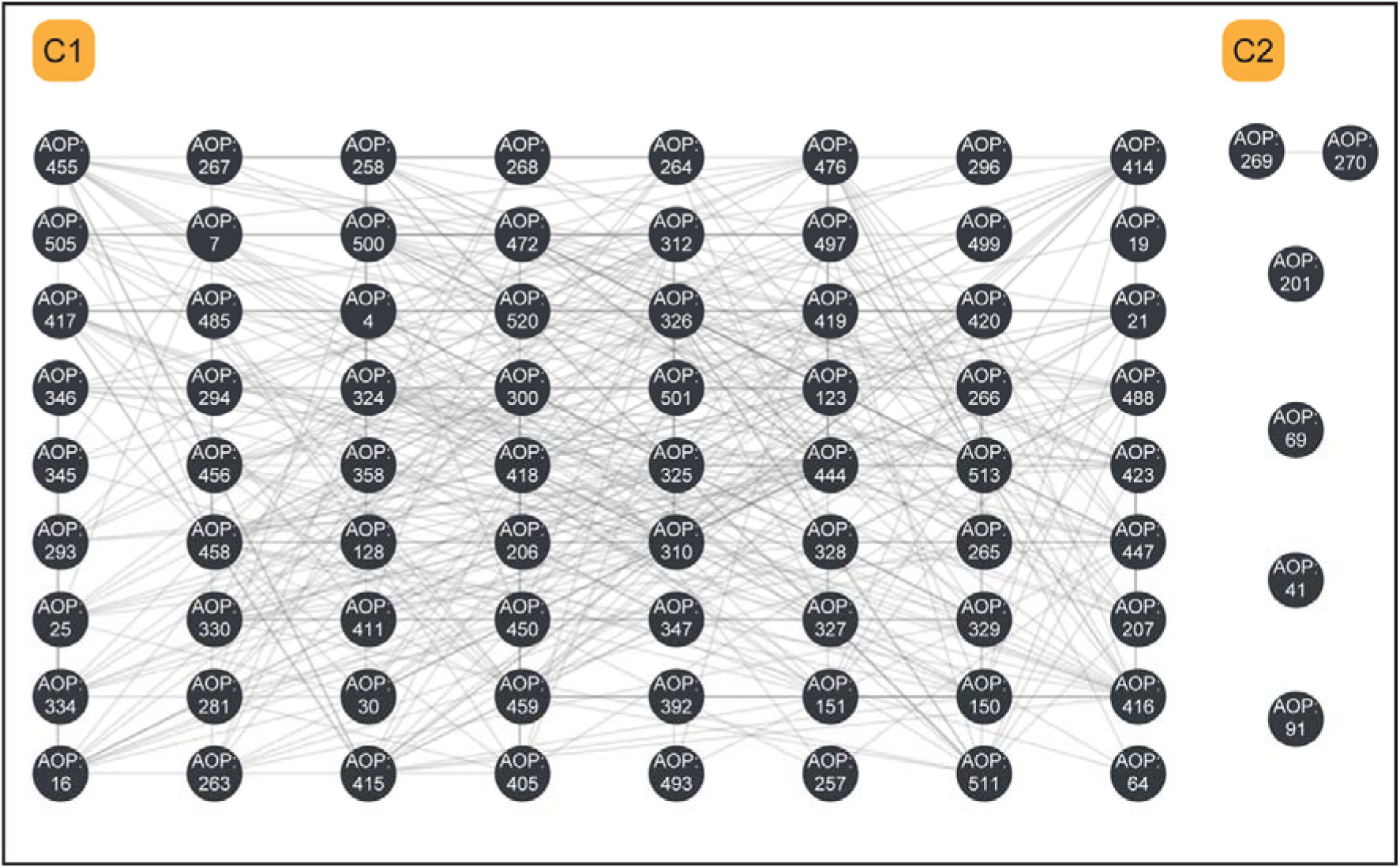
Undirected network of cadmium–AOPs. Each node corresponds to a cadmium– AOP, and an edge between two nodes denotes that the two AOPs share at least one KE. This undirected network has 2 connected components (with two or more nodes) which are labeled as C1 and C2, and 4 isolated nodes.

In the directed arsenic-AOP network, it was observed that the MIE ‘N/A, Mitochondrial dysfunction 1’ (KE:177) has the highest out-degree of 9, while the AOs ‘Increased Mortality’ (KE:351) and ‘Decrease, Population growth rate’ (KE:360) have the highest in-degree of 9 (Table S16). The MIE ‘Increased, Reactive oxygen species’ (KE:1115) has the highest betweenness centrality value, signifying that several toxicity pathways are passing through it in this network (Figure S3) [19]. The KE ‘Tissue resident cell activation’ (KE:1492) has the highest eccentricity, denoting that it is the most remotely placed KE in this network (Figure S4) [60].

In the directed cadmium-AOP network, it was observed that the MIE ‘N/A, Mitochondrial dysfunction 1’ (KE:177) has the highest out-degree of 13, while the KE ‘increased Vascular endothelial dysfunction’ (KE:1928) and AO ‘Decrease, Population growth rate’ (KE:360) have the highest in-degree of 9 (Table S17). The MIE ‘N/A, Mitochondrial dysfunction 1’ (KE:177) has the highest betweenness centrality value, signifying that several toxicity pathways are passing through it in this network (Figure S6) [19]. The MIE ‘DNA adduct formation’ (KE:10058) and KE ‘Releasing, Apoptosis-Inducing Factor (AIF)’ (KE:2031) has the highest eccentricity, denoting that they are the most remotely placed KEs in this network (Figure S7) [60].

It was observed that 83 of 152 KEs in the directed arsenic-AOP network were associated to inorganic arsenic-induced toxicity through this systematic data-centric approach (Table S16). Similarly, 156 of 263 KEs were associated with inorgnic cadmium-induced toxicity within the directed cadmium-AOP network (Table S17). To further support Key Events (KEs) within directed arsenic-AOP and cadmium-AOP networks that are not linked to the stressors through our systematic approach, natural language processing (NLP) based tools like Abstract Sifter [61] and AOP-HelpFinder [62,63] was used, and associated published evidence (Tables S15-S16) was idenitified. Finally, toxicity pathways were selected in both humans and aquatic organisms to explore mechanisms underlying stressor-induced toxicities (Figure 4).

**Figure 4:**
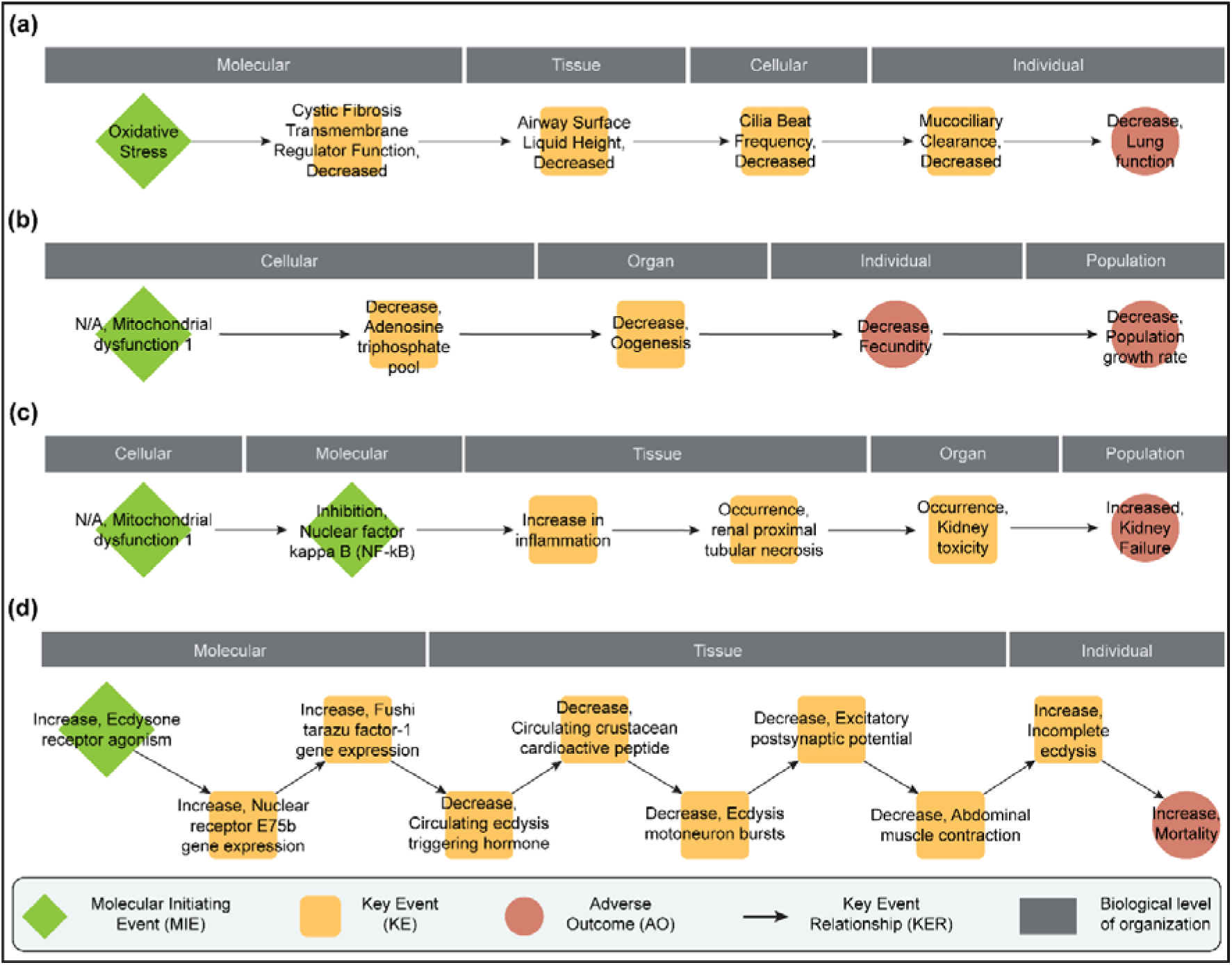
Toxicity pathways explored in case studies of arsenic- and cadmium-induced toxicities. **(a)** Arsenic-induced toxicity pathway leading to decreased lung function in humans. **(b)** Arsenic-induced toxicity pathway leading to decline in population growth rate in aquatic organisms. **(c)** Cadmium-induced toxicity pathway leading to kidney failure in humans. **(d)** Cadmium-induced toxicity pathway leading to increased mortality in aquatic organisms.

#### 3.2.1. Toxicity pathway linking arsenic exposure to decreased lung function in humans

Arsenic exposure has been shown to induce oxidative stress in both human *in vitro* studies and *in vivo* rat models [64,65]. Bomberger *et al.* [66] demonstrated that arsenic promotes degradation of the cystic fibrosis transmembrane conductance regulator (CFTR) function, leading to reduced airway surface liquid volume, decreased cilia beat frequency, and impaired mucociliary clearance in human airway epithelial cells. This impairment in clearance can cause pathogen accumulation, ultimately reducing lung function [67]. Additionally, a population-wide study found an association between arsenic exposure and impaired lung function [68]. Thus, we were able to explore a potential toxicity pathway underlying arsenic-induced decrease in lung function by leveraging various published evidence (Figure 4a).

#### 3.2.2. Toxicity pathway linking arsenic exposure to decreased population growth in aquatic organisms

Arsenic exposure has been shown to disrupt mitochondrial membrane potential in PLHC-1 fish cells, with significant damage observed at various concentrations and exposure durations [69]. Additionally, arsenic exposure has been shown to impair ATP synthesis, resulting in a reduced ATP pool in Mediterranean polychaete species [70]. Arsenic exposure has also been found to impair oocyte development in freshwater fish from India, with an increased number of atretic follicles observed, ultimately leading to reduced fecundity [71]. Moreover, Zhang *et al.* [72] highlighted the detrimental effects of arsenic on the survival of various aquatic organisms through a comprehensive literature review. Thus, we were able to explore a potential toxicity pathway underlying arsenic-induced decline in population growth among aquatic organisms by leveraging various published evidence (Figure 4b).

#### 3.2.3. Toxicity pathway linking cadmium exposure to kidney failure in humans

Cadmium has been shown to induce mitochondrial dysfunction in HEK293 cells by increasing membrane permeability, reducing membrane potential, and causing mitochondrial swelling [73]. Cadmium exposure disrupted kidney mitochondria, altered Nrf2 signaling pathway and triggered inflammatory responses in chickens [74]. In rats, cadmium exposure lead to renal tubular failure and subsequent kidney toxicity [75]. Furthermore, several epidemiological studies have demonstrated a strong correlation between cadmium exposure and renal damage in humans [76,77]. Thus, we were able to explore a potential toxicity pathway underlying cadmium-induced kidney failure in humans by leveraging various published evidence (Figure 4c).

#### 3.2.4. Toxicity pathway linking cadmium exposure to increased mortality in aquatic organisms

Cadmium exposure in aquatic organisms has been shown to induce agonism of the ecdysone receptor, resulting in the upregulation of stress-related genes and alterations in ecdysteroid levels [78,79]. In *Drosophila*, the E75b gene, an ecdysteroid-responsive gene, is upregulated upon ecdysone activation [80]. Additionally, the FTZ-F1 nuclear receptor is essential for mediating developmental responses to ecdysone in *Drosophila*, which correlates with increased expression of the FTZ-F1 gene [81,82]. Furthermore, in insects, elevated ecdysteroid levels have been linked to the downregulation of ecdysis triggering hormone (ETH) release [83]. In crabs, a reduction in crustacean cardioactive peptide (CCAP) levels was observed following the knockout of ETH, suggesting that CCAP levels decrease in the absence of ETH [84]. As a neuropeptide and master regulator of ecdysis [85,86], CCAP’s downregulation can result in impaired neuronal activity [86,87], including diminished muscle contractions [88,89], ultimately leading to incomplete ecdysis and increased mortality in crustaceans [85,86]. Subsequently, Luo *et al.* [90] reported that cadmium exposure inhibited ecdysis in freshwater crabs in a dose-dependent manner, indicating that cadmium could potentially be lethal to crustaceans. Thus, we were able to explore a potential toxicity pathway underlying cadmium-induced moratlity among aquatic organisms by leveraging various published evidence (Figure 4d).

### 3.3. Stressor-species networks associating impacts of inorganic arsenic and cadmium compounds with ecologically relevant species

A stressor-species network reveals the intricate interactions between environmental stressors and their effects on ecosystems [23,25,26]. In this study, data within the ECOTOX knowledgebase [36] was used to construct three types of stressor-species networks: one based on acute toxicity data (LC_50_ and EC_50_), one based on chronic toxicity data (NOEC) and another using bioconcentration factor (BCF) data; to explore the impacts of inorganic arsenic and cadmium on ecosystems, and identify vulnerable species (Methods; Supplementary Information). It was observed that 167 species (across 9 ECOTOX species groups) are linked to 15 inorganic arsenic compounds and 644 species (across 12 ECOTOX species groups) are linked to 8 inorganic cadmium compounds (Figure 5a; Table S8) in the acute toxicity network. Species within the ‘Fish’ ECOTOX group were most frequently tested for inorganic arsenic-induced acute toxicity, while ‘Crustaceans’ were most frequently tested for inorganic cadmium-induced acute toxicity (Table S8). Notably, Sodium arsenite (CAS: 7784-46-5) and Cadmium chloride (CAS: 10108-64-2) are documented as being toxic to the highest number of species (Table S8). Additionally, *Daphnia magna* has been tested with the highest number of both arsenic and cadmium compounds (Table S8).

**Figure 5:**
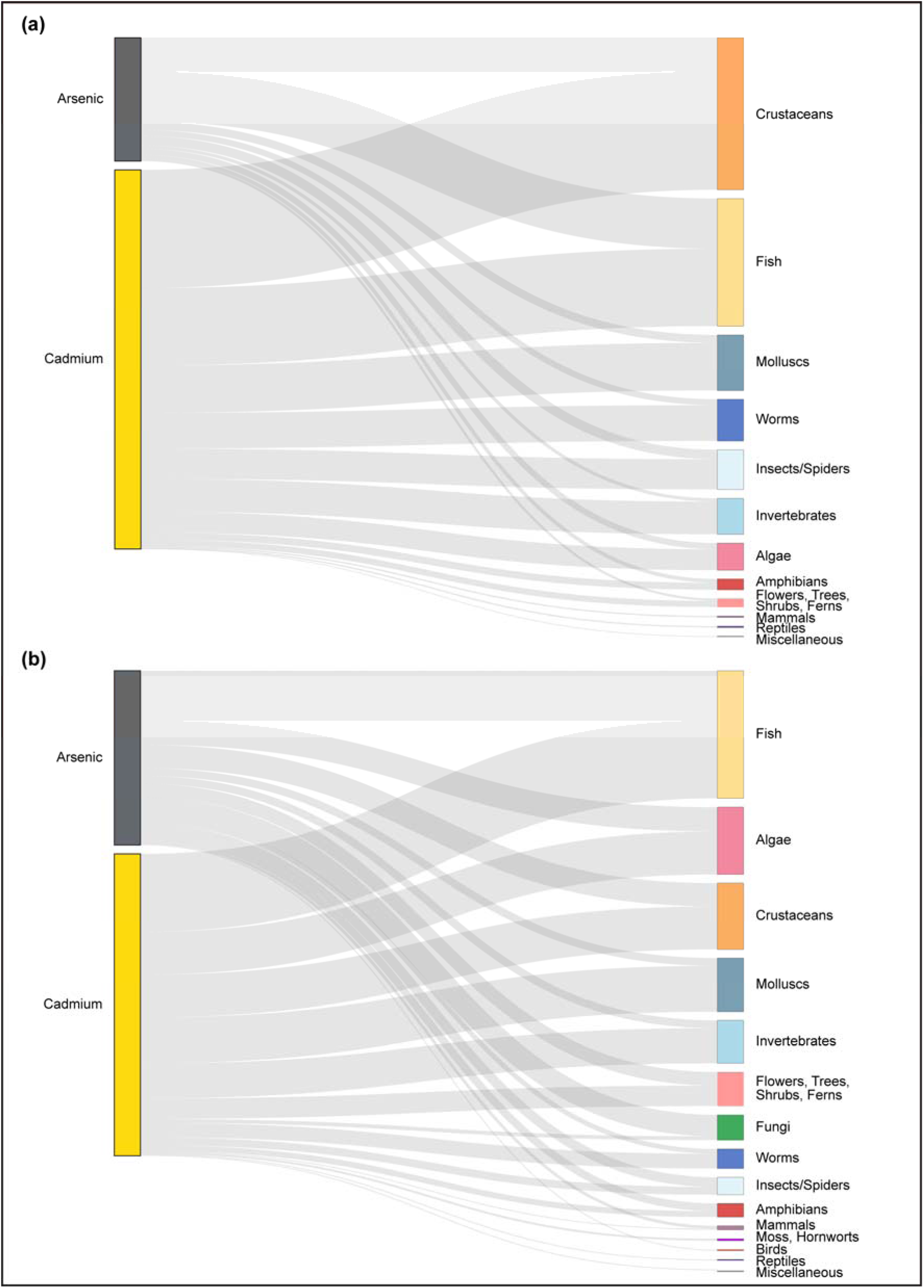
Sankey plot depicting associations between inorganic arsenic and cadmium compounds, and ECOTOX species groups through their: **(a)** acute toxicity concentration data (LC_50_ and EC_50_); and **(b)** chronic toxicity concentration data (NOEC). The plot provides associations between inorganic arsenic and cadmium with 15 ECOTOX species groups.

In the stressor-species network based on chronic toxicity data, it was observed that 130 species (across 12 ECOTOX species groups) are linked to 16 inorganic arsenic compounds and 264 species (across 14 ECOTOX species groups) are linked to 4 inorganic cadmium compounds (Figure 5b; Table S8). Species within the ‘Fish’ ECOTOX group were most frequently tested for both inorganic arsenic-induced and cadmium-induced chronic toxicity (Table S8). Notably, Sodium arsenite (CAS: 7784-46-5) and Cadmium chloride (CAS: 10108-64-2) are documented as being toxic to the highest number of species (Table S8). Additionally, *Danio rerio* has been tested with the highest number of arsenic compounds, while *Cyprinus carpio* and *Oncorhynchus mykiss* have been tested with the highest number of cadmium compounds (Table S8).

Similarly, in the BCF-based network, it was observed that 32 species (across 10 ECOTOX species groups) are linked to 5 inorganic arsenic compounds and 147 species (across 11 ECOTOX species groups) to 4 inorganic cadmium compounds (Figure 6; Table S9). Species within the ‘Flowers, Trees, Shrubs, Ferns’ ECOTOX group were documented to absorb most arsenic compounds, while ‘Algae’ were documented to absorb most cadmium compounds (Table S9). Notably, Disodium arsenate (CAS:7778-43-0) and Cadmium chloride (CAS:10108-64-2) were documented as having bioconcentration potential in largest number of species (Table S9). Additionally, *Daphnia magna* has been documented to absorb the highest number of arsenic compounds, while *Lemna minor* has been documented to absorb the highest number of cadmium compounds (Table S9).

**Figure 6:**
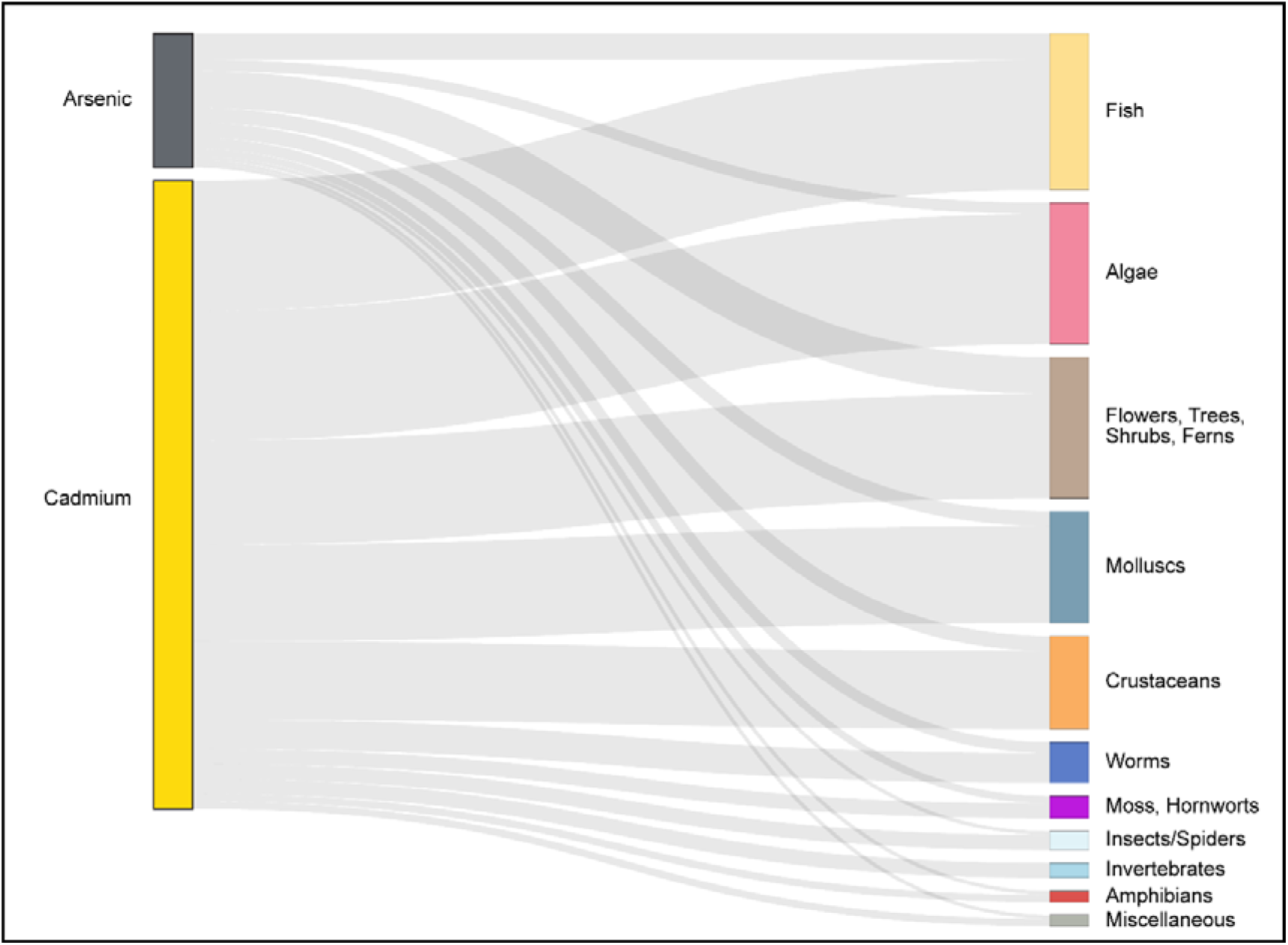
Sankey plot depicting associations between inorganic arsenic and cadmium compounds, and ECOTOX species groups through their bioconcentration factors (BCFs). The plot provides associations between inorganic arsenic and cadmium with 11 ECOTOX species groups.

### 3.4. Species Sensitivity Distributions (SSDs) for inorganic arsenic and cadmium compounds

Species Sensitivity Distributions (SSDs) are statistical tools used in ecological risk assessments to evaluate the sensitivity of various species to toxic environmental substances [23,91–93]. SSDs are utilized to derive hazard concentration values like HC05, which represent concentrations expected to be harmful to 5% of species within an ecosystem [23,91–93]. Additionally, the toxicity-normalized SSD (SSDn) is a novel approach that normalizes chemical toxicity data based on a reference species, thereby improving the comparison of chemicals with similar toxicological profiles [53,54]. In this study, both SSD and SSDn were computed for inorganic arsenic and cadmium compounds based on acute and chronic toxicity data compiled in the ECOTOX database (Methods; Supplementary Information). Further, the derived HC05 values were compared with the toxicity concentrations in the stressor-species network, and a species was identified as sensitive to a chemical if the corresponding toxicity value was below the HC05 value. These sensitive species associations were highlighted within the stressor-species network to provide a comprehensive view of all the sensitive species associated with inorganic arsenic and cadmium compounds. In comparison with the stressor-species networks constructed by Sahoo *et al*. [23], this approach offers a novel perspective, as it integrates hazard concentrations into network visualization, thereby enhancing our understanding of the toxicity impacts across different species. Such stressor-species visualizations are accessible on the associated website: https://cb.imsc.res.in/heavymetaltox/. Figure 7 shows an example of the stressor-species network constructed based on chronic toxcity values associated with inorganic arsenic compounds.

**Figure 7:**
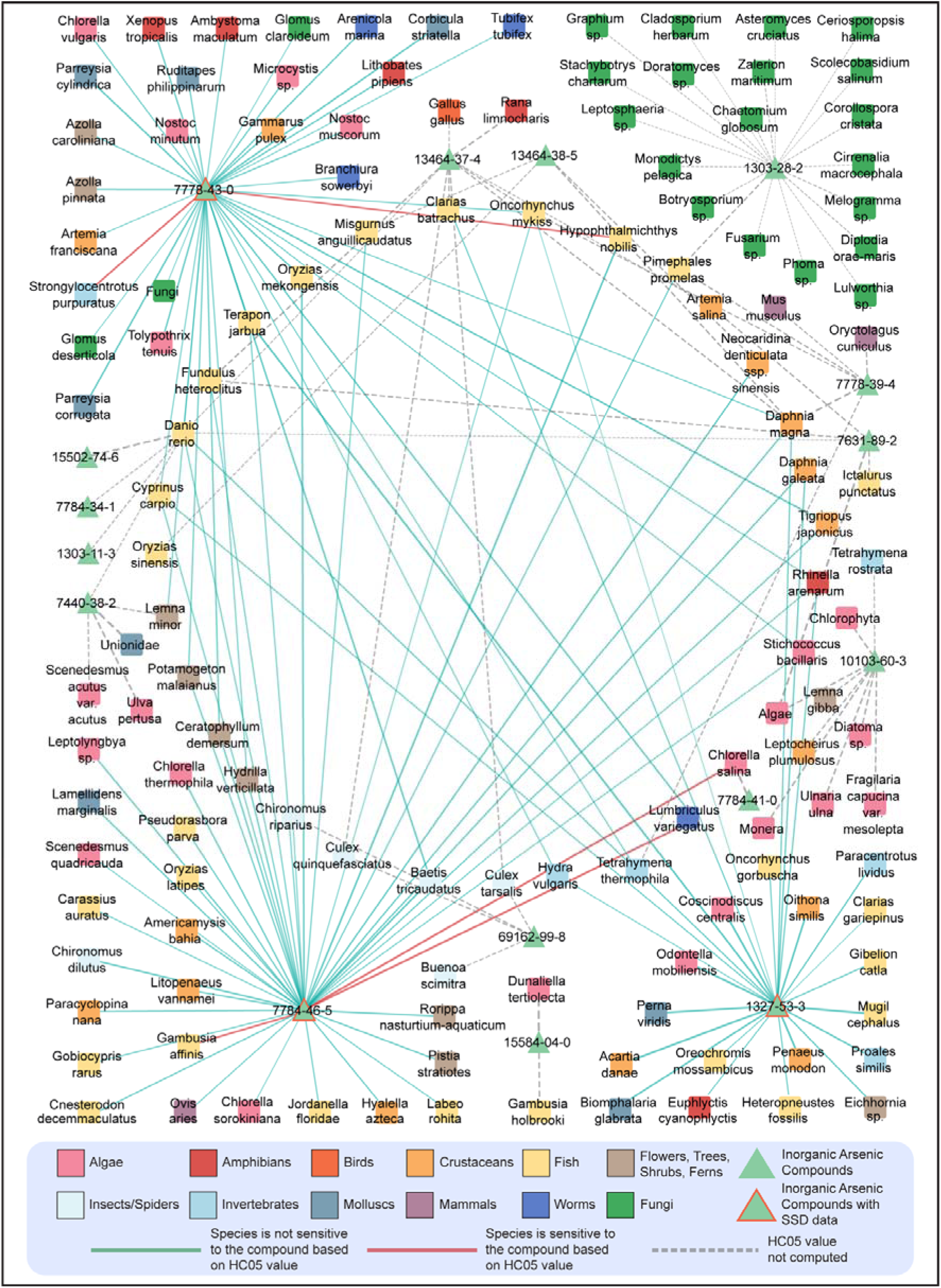
Stressor-species network constructed for inorganic arsenic compounds using the chronic toxicity concentrations as edge weights. The network comprises 16 inorganic arsenic compounds, 130 species belonging to 12 ECOTOX species groups, and 179 stressor-species links. The 4 inorganic arsenic compounds for which SSDs were computed are highlighted with a ‘red’ border. A stressor-species link is shown in ‘red’ if the species is sensitive to the corresponding stressor, otherwise it is shown in ‘blue’. A ‘gray dotted’ link corresponds to a stressor-species pair, where the HC05 values was not derived for the stressor.

#### 3.4.1. SSDs based on acute toxicity endpoints

The acute toxicity endpoints for aquatic species from the ECOTOX database was used to construct SSDs, and the HC05 values were derived for 4 inorganic arsenic compounds, namely Arsenic oxide (CAS:1327-53-3), Disodium arsenate (CAS:7778-43-0), Sodium arsenite (CAS:7784-46-5), and Trisodium arsenate (CAS:13464-38-5), and 4 inorganic cadmium compounds, namely Cadmium (CAS:7440-43-9), Cadmium chloride (CAS:10108-64-2), Cadmium sulfate (CAS:10124-36-4), and Cadmium nitrate (CAS:10325-94-7), using both the US EPA SSD Toolbox and the R-based ssdtools package (Methods; Table S18). It was observed that both tools consistently identified the same best-fit models, and derived similar HC05 values for all chemicals (Table S19). Figures S8-S15 present the SSD plots for each chemical generated using the corresponding best-fit models from both tools.

The HC05 values indicate the toxic effects of chemicals across various species, with lower HC05 values signifying a higher toxic potential [94]. Based on the model-averaged HC05 values, it was observed that cadmium compounds are in general more toxic than the arsenic compounds, aligning with the findings of Karthikeyan *et al.* [92] regarding sea water quality criterion for heavy metals. Furthermore, SSDs can help identify species that are particularly sensitive to specific chemicals [92]. It was observed that species in the ECOTOX species groups ‘Crustaceans’ and ‘Algae’ were commonly sensitive to both inorganic arsenic and cadmium compounds. Based on the annotated stressor-species network (Table S8), it was observed that *Scenedesmus acutus* var. *acutus* was sensitive to more than one inorganic arsenic compound, while *Eisenia fetida*, *Oncorhynchus* tshawytscha, and *Salmo trutta* were sensitive to more than one inorganic cadmium compound. Moreover, *Oncorhynchus kisutch,* which is sensitive to Cadmium chloride (CAS:10108-64-2), and *Periophthalmus waltoni,* which is sensitive to Cadmium (CAS:7440-43-9), were also documented to bioaccumulate these chemicals (Tables S8-S9).

Additionally, the HC05 values were derived using the SSDn approach for each individual chemical (Methods; Figures S16-S17; Table S19). It was observed that the SSDn method provided consistent estimates of HC05 values for all these chemicals (Methods; Table S19), highlighting its utility as a reliable tool for assessing the acute toxicity of heavy metal pollutants.

#### 3.4.2. SSDs based on chronic toxicity endpoints

Similarly, SSDs were constructed by utilizing the chronic toxicity endpoints for aquatic species from the ECOTOX database for 3 inorganic arsenic compounds, namely Arsenic oxide (CAS:1327-53-3), Disodium arsenate (CAS:7778-43-0), and Sodium arsenite (CAS:7784-46-5) and 4 inorganic cadmium compounds, namely Cadmium (CAS:7440-43-9), Cadmium chloride (CAS:10108-64-2), Cadmium sulfate (CAS:10124-36-4), and Cadmium nitrate (CAS:10325-94-7), and it was observed that the results were consistent across both the tools (Methods; Table S20). Figures S18-S24 present the SSD plots for each chemical generated using the corresponding best-fit models from both tools. Thereafter, the individual HC05 values were derived, and it was observed that cadmium compounds are more toxic than the arsenic compounds (Table S20). Furthermore, it was observed that species in the ECOTOX species groups ‘Crustaceans’ and ‘Fish’ were commonly sensitive to both inorganic arsenic and cadmium compounds. Based on the annotated stressor-species network (Table S8), it was observed that *Salmo trutta* was sensitive to more than one inorganic cadmium compound, while there were no species that were sensitive to more than one inorganic arsenic compound. Moreover, *Hyalella azteca* which is sensitive to Cadmium chloride (CAS:10108-64-2), was also documented to bioaccumulate this chemical (Tables S8-S9). Additionally, the HC05 values were derived using the SSDn approach (Methods; Figures S25-S26; Table S21), and it was observed that this method provided consistent estimates of HC05, other than that for cadmium.

In conclusion, both the acute and chronic toxicity endpoints for aquatic organisms from the ECOTOX database were utilized to construct SSDs and derive HC05 values for the inorganic arsenic and cadmium compounds. Additionally, we employed the SSDn approach and found that it provided consistent HC05 estimates for these chemicals.

## 4. Conclusion

Arsenic and cadmium are highly toxic environmental pollutants with significant health and ecological risks. Network toxicology provides crucial insights into the complex biological responses associated with exposure to these pollutants, enhancing our understanding of their systemic impacts across species and ecosystems. In this study, diverse network-based approaches were applied to understand the toxicities of inorganic arsenic and cadmium compounds. First, a list of 62 inorganic arsenic and 18 inorganic cadmium compounds was systematically curated from six exposome-relevant databases, and their toxicological endpoints were retrieved. Next, a set of 342 high-confidence AOPs was compiled from AOP-Wiki, that comprised 1163 unique Key Events (KEs), and the toxicological endpoints were systematically mapped to these KEs. This enabled construction of a stressor-AOP network linking 51 and 78 high-confidence AOPs to inorganic arsenic and cadmium, respectively, each with Level 5 relevance and coverage score ≥0.4. Next, undirected and directed AOP networks were constructed, and four case studies were performed to explore mechanisms of toxicity in response to arsenic and cadmium exposure in humans and other species. Further, using ECOTOX, stressor-species networks were constructed based on acute toxicity data, linking 15 inorganic arsenic and 8 inorganic cadmium compounds to 167 and 644 species, respectively, and on chronic toxicity data, linking 16 inorganic arsenic and 4 inorganic cadmium compounds to 130 and 264 species, respectively. Through these networks, it was observed that species belonging to ‘Fish’ and ‘Crustaceans’ ECOTOX groups were most vulnerable to inorganic arsenic and cadmium compounds. Additionally, bioconcentration factor (BCF) data was used to link 5 inorganic arsenic and 4 inorganic cadmium compounds to 32 and 147 species, respectively, and it was observed that species belonging to ‘Fish’, ‘Algae’, and ‘Flowers, Trees, Shrubs, Ferns’ ECOTOX groups were commonly documented to bioaccumalte these compounds. Then, the HC05 values were derived by constructing Species Sensitivity Distributions (SSDs) and toxicity-normalized SSDs (SSDn) for inorganic arsenic and cadium compounds affecting aquatic species. This data, integrated into the stressor-species networks, enabled identification of sensitive species groups. The networks and related data generated in this study are freely available for further research at https://cb.imsc.res.in/heavymetaltox/. Overall, this study leverages diverse network toxicological approaches to comprehensively assess the toxicological impacts of arsenic and cadmium exposure across both human and ecological contexts.

However, the inferences from this study are constrained by the limitations of the underlying data, particularly the toxicity data obtained from ECOTOX. For SSDs, the lack of geographical specificity restricts the applicability of inferences to particular locations. Additionally, the higher fold change value observed when comparing HC05 values derived from SSDn and single-compound SSDs for cadmium may indicate that, despite selecting the most sensitive species available, the species may not be sensitive enough for assessing this particular chemical. Furthermore, the approaches used in this study are limited in their ability to assess the bioaccumulation of arsenic and cadmium compounds across trophic levels.

Nonetheless, the network-based approaches employed in this study have enhanced our understanding of inorganic arsenic- and cadmium-induced toxicities. The inclusion of ecotoxicological endpoints from the ECOTOX database, alongside mammalian toxcicity endpoints, has broadened the scope and applicability of associated AOPs in advancing our understanding of arsenic- and cadmium-induced toxicities. The stressor-AOP network provided a mechanistic perspective by highlighting pathways underlying toxic effects associated with inorganic arsenic and cadmium exposure. The stressor-species networks identified species vulnerable to exposure from these compounds, as well as those with potential for bioaccumulation. Additionally, the SSD and toxicity-normalized SSD (SSDn) approaches facilitated comparisons among these compounds, offering a framework for prioritizing them in further toxicity assessment studies. By integrating SSD data with the stressor-species networks, species groups particularly sensitive to exposure from these compounds were identified, thereby strengthening the utility of stressor-species networks in ecological risk assessment. In sum, this study provides a comprehensive understanding of the toxicity mechanisms and impacts of inorganic arsenic and cadmium compounds on human and ecosystem health, thereby supporting a One Health-focused approach towards enhanced regulatory and mitigation strategies for these compounds.

## Data availability

The data associated with this study is contained in the article, or in the supplementary information files, or in the associated website: https://cb.imsc.res.in/heavymetaltox/.

## Acknowledgement

Areejit Samal would also like to acknowledge funding from the Department of Atomic Energy (DAE), Government of India via Apex project to The Institute of Mathematical Sciences (IMSc) Chennai. This work was also undertaken as part of the project on ‘Marine Ecotoxicology and Ecological Risk Assessment’ (MEERA) Programme [MoES/OSMART/EFC/2021 dated 07.03.2022]. The authors are thankful to the Ministry of Earth Sciences (MoES), and the Director, National Centre for Coastal Research (NCCR), Government of India for the support and encouragement to carry out this work. The funders have no role in study design, data collection, data analysis, manuscript preparation or decision to publish.

## CRediT author contribution statement

**Nikhil Chivukula:** Conceptualization, Data Compilation, Data Curation, Formal Analysis, Methodology, Software, Visualization, Writing; **Shreyes Rajan Madgaonkar**: Conceptualization, Data Compilation, Data Curation, Formal Analysis, Methodology, Software, Visualization, Writing; **Kundhanathan Ramesh:** Data Compilation, Data Curation, Formal Analysis, Visualization; **Swetha Mangot:** Data Compilation, Data Curation; **Panneerselvam Karthikeyan:** Formal Analysis, Methodology; **Shambanagouda Rudragouda Marigoudar:** Conceptualization, Formal Analysis, Methodology, Writing; **Krishna Venkatarama Sharma:** Conceptualization, Formal Analysis, Methodology, Writing; **Areejit Samal:** Conceptualization, Supervision, Formal Analysis, Methodology, Writing.

## Supplementary Tables

**Table S1**: This table contains the list of 62 inorganic arsenic compounds and 18 inorganic cadmium compounds identified from ToxCast, Comparative Toxicogenomics Database (CTD), DEDuCT, NeurotoxKb, and ECOTOX. For each chemical, the table provides the primary chemical identifier, CAS chemical identifier, PubChem chemical identifier, chemical name, and the heavy metal present in the chemical. Further, for each chemical, the table provides chemical Kingdom, chemical Superclass, chemical Class predicted by ClassyFire, its chemical structure in SMILES, InChI and InChIKey format, its molecular weight in g/mol computed using RDKit, its product use category and functional use reported in CPDat, its presence as high production volume (HPV) chemical in US HPV or OECDHPV list, its presence in substances of very high concern (SVHC) list, its presence in REACH list of prohibited chemicals, and its presence in human biospecimens as documented in TeXAs (https://cb.imsc.res.in/texas/) and Exposome Explorer (http://exposome-explorer.iarc.fr) (separated by ’|’ symbol).

**Table S2**: This table contains the curated list of 342 high confidence adverse outcome pathways from AOP-Wiki. For each ecotoxicologically-relevant AOP, the table provides the corresponding information on AOP identifier, AOP title, Handbook Version that was followed while constrcution of the AOP, Organisation for Economic Co-operation and Development (OECD) status, and taxonomic applicability of the AOP (separated by ’|’ symbol).

**Table S3**: This table contains information on the 1163 Key Events (KEs) present in the curated list of 342 AOPs. For each KE, the table provides the corresponding information on KE identifier, KE title, level of biological organization, and associated AOP identifier(s) (separated by ’|’ symbol).

**Table S4**: This table contains information on Key Event Relationships (KERs) present in each of the curated list of 342 AOPs. For each AOP, the table provides the AOP identifier, corresponding KER identifier, upstream KE identifier, downstream KE identifier, MIE(s) among upstream and downstream KEs (separated by ’|’ symbol), AO(s) among upstream and downstream KEs (separated by ’|’ symbol), adjacency of KER, weight of evidence of KER, and quantitative understanding of KER. Note that the KER identifiers starting with 10000 were manually assigned by the authors as the KER was mentioned in the AOP page but was not assigned an identifier in AOP-Wiki.

**Table S5**: This table contains information on 579 Key Events (KEs) from 342 high confidence AOPs that are associated with inorganic arsenic and cadmium. For each KE, the table provides corresponding information on KE identifier, CAS chemical identifier (separated by ’|’) for associated chemical(s), associated heavy metal(s), source(s) from which the associations are inferred (separated by ’|’ symbol), associated ToxCast assay endpoint(s) (separated by ’|’ symbol), associated CTD disease(s) (separated by ’|’ symbol), associated CTD phenotype(s) (separated by ’|’ symbol), associated DEDuCT endpoint(s) (separated by ’|’ symbol), associated NeurotoxKb endpoint(s) (separated by ’|’ symbol), AOP Identifier(s) from which KE mapping was inferred (separated by ’|’ symbol), and ECOTOX measurements and trends [mentioned as (measurement,trend)] (separated by ’|’ symbol).

**Table S6**: This table provides edge list for the complete stressor-AOP network for arsenic and cadmium. For each edge in the network, the table provides the stressor, AOP identifier, computed coverage score and level of relevance.

**Table S7**: This table provides conversion factors used to convert the toxicity concentration values within ECOTOX to ppm equivalent values. Here, MW stands for the molecular weight of the corresponing chemical.

**Table S8:** This table provides edge list for the stressor-species network, for the inorganic compounds of arsenic and cadmium, constructed using concentration value for acute toxicity endpoints (LC50 and EC50) and chronic toxicity endpoints (NOEC). For each edge in the network, the table provides toxicity endpoint type, the CAS chemical identifier, chemical name as in ECOTOX, the heavy metal present in the chemical, Latin name, ECOTOX species group and habitat (separated by ‘|’ symbol) for the species, considered toxicity concentration value in ppm equivalent unit, logarithm of concentration value in ppm equivalent unit (rounded up to 6 decimals), and whether the species is sensitive to the chemical based on the corresponding derived HC05 value.

**Table S9**: This table provides edge list for the stressor-species network for inorganic compounds of arsenic and cadmium, constructed using bioconcentration factors. For each edge in the network, the table provides the CAS chemical identifier, chemical name, the heavy metal present in the chemical, Latin name and ECOTOX species group for the species, bioconcentration factor in L/kg, and logarithm of bioconcentration factor value (rounded up to 6 decimals).

**Table S10**: This table contains acute toxicity concentration data used to construct species sensitivity distributions (SSDs) for four inorganic arsenic compounds and four inorganic cadmium compounds. Further, the table provides information on the CAS chemical identifier, chemical name, the heavy metal present in the chemical, Latin name and ECOTOX species group for the species, value and unit of the toxicity concentration of the chemical.

**Table S11**: This table contains chronic toxicity concentration data used to construct species sensitivity distributions (SSDs) for three inorganic arsenic compounds and four inorganic cadmium compounds. Further, the table provides information on the CAS chemical identifier, chemical name, the heavy metal present in the chemical, Latin name and ECOTOX species group for the species, value and unit of the toxicity concentration of the chemical.

**Table S12**: This table contains acute toxicity concentration data used to construct toxicity-normalized species sensitivity distributions (SSDn) for inorganic arsenic and inorganic cadmium. Further, the table provides information on the heavy metal present in the chemical, Latin name and ECOTOX species group for the species, value and unit of the normalized toxicity concentration of the chemical.

**Table S13**: This table contains chronic toxicity concentration data used to construct toxicity-normalized species sensitivity distributions (SSDn) for inorganic arsenic and inorganic cadmium. Further, the table provides information on the heavy metal present in the chemical, Latin name and ECOTOX species group for the species, value and unit of the normalized toxicity concentration of the chemical.

**Table S14**: This table contains information on the evidence and the connected component for each of the 51 arsenic-AOPs. For each AOP, the table provides the corresponding AOP identifier, computed fraction of KERs with ’High’ evidence (i.e., F(High)), computed fraction of KERs with ’Moderate’ evidence (i.e., F(Moderate)), computed fraction of KERs with ’Low’ evidence (i.e., F(Low)), computed fraction of KERs with ’Not Specified’ evidence (i.e., F(Not Specified)), computed cumulative weight of evidence (WoE), and the connected component identifier in the undirected AOP network associated with arsenic.

**Table S15**: This table contains information on the evidence and the connected component for each of the 78 cadmium-AOPs. For each AOP, the table provides the corresponding AOP identifier, computed fraction of KERs with ’High’ evidence (i.e., F(High)), computed fraction of KERs with ’Moderate’ evidence (i.e., F(Moderate)), computed fraction of KERs with ’Low’ evidence (i.e., F(Low)), computed fraction of KERs with ’Not Specified’ evidence (i.e., F(Not Specified)), computed cumulative weight of evidence (WoE), and the connected component identifier in the undirected AOP network associated with cadmium.

**Table S16**: This table contains information on the computed network statistics and literature evidence of the association with Arsenic for each of the 152 KEs present in the largest connected component (C1) of the arsenic-AOP network. For each KE, the table provides the corresponding KE identifier, KE title, level of biological organization, KE type, betweenness centrality value, eccentricity value, in-degree value, out-degree value, convergence information, organism in which arsenic exposure is studied, study type, chemical that was tested, dosage information, abbreviated description of association with arsenic exposure, and the corresponding reference.

**Table S17**: This table contains information on the computed network statistics and literature evidence of the association with Cadmium for each of the 263 KEs present in the largest connected component (C1) of the cadmium-AOP network. For each KE, the table provides the corresponding KE identifier, KE title, level of biological organization, KE type, betweenness centrality value, eccentricity value, in-degree value, out-degree value, convergence information, organism in which arsenic exposure is studied, study type, chemical that was tested, dosage information, abbreviated description of association with cadmium exposure, and the corresponding reference.

**Table S18**: This table contains information on the derived HC05 values from the acute toxicity data of four inorganic arsenic compounds and four inorganic cadmium compounds. For each of these chemicals, the table provides the CAS chemical identifier, chemical name, the heavy metal present in the chemical, number of associated species in the constructed SSD, number of ECOTOX species groups in the constructed SSD, HC05 value, HC05 lower confidence limit (LCL), HC05 upper confidence limit (UCL) for the corresponding best-fit model, model average HC05 value and corresponding standard error (SE) of the model average HC05 value derived from US EPA SSD Toolbox and ssdtools package. The values are given in equivalent ppm units and rounded up to 4 decimal places.

**Table S19**: This table contains information on the derived HC05n values and the corresponding estimated HC05 values from the acute toxicity data associated with four inorganic arsenic compounds and four inorganic cadmium compounds. For each of these chemicals, the table provides the CAS chemical identifier, chemical name, the heavy metal present in the chemical, the representative species used in SSDn computation (nSpecies), species mean toxicity value (SMTV) corresponding to the nSpecies, best-fit model, derived HC05n value, HC05n lower confidence limit (LCL), HC05n upper confidence limit (UCL) for the corresponding best-fit model, model average HC05n value and corresponding standard error (SE) of the model average HC05n value derived from US EPA SSD Toolbox and ssdtools package. The table also provides information on the estimated HC05 value computed from HC05n value, the HC05 value as computed in corresponding single-chemical SSD, and fold change of the estimated HC05 to single-chemical HC05. The HC05n values are mentioned in corresponding nSpecies equivalents. The SMTV and other HC05 values are given in equivalent ppm units and rounded up to 4 decimal places.

**Table S20**: This table contains information on the derived HC05 values from the chronic toxicity data of three inorganic arsenic compounds and four inorganic cadmium compounds. For each of these chemicals, the table provides the CAS chemical identifier, chemical name, the heavy metal present in the chemical, number of associated species in the constructed SSD, number of ECOTOX species groups in the constructed SSD, HC05 value, HC05 lower confidence limit (LCL), HC05 upper confidence limit (UCL) for the corresponding best-fit model, model average HC05 value and corresponding standard error (SE) of the model average HC05 value derived from US EPA SSD Toolbox and ssdtools package.

**Table S21**: This table contains information on the derived HC05n values and the corresponding estimated HC05 values from the chronic toxicity data associated with three inorganic arsenic compounds and four inorganic cadmium compounds. For each of these chemicals, the table provides the CAS chemical identifier, chemical name, the heavy metal present in the chemical, the representative species used in SSDn computation (nSpecies), species mean toxicity value (SMTV) corresponding to the nSpecies, best-fit model, derived HC05n value, HC05n lower confidence limit (LCL), HC05n upper confidence limit (UCL) for the corresponding best-fit model, model average HC05n value and corresponding standard error (SE) of the model average HC05n value derived from US EPA SSD Toolbox and ssdtools package. The table also provides information on the estimated HC05 value computed from HC05n value, the HC05 value as computed in corresponding single-chemical SSD, and fold change of the estimated HC05 to single-chemical HC05. The HC05n values are mentioned in corresponding nSpecies equivalents.

**Table S22**: This table contains information on 17749 chemical-gene-phenotype-disease (CGPD) tetramers constructed for inorganic compounds of arsenic and cadmium from Comparative Toxicogenomics Database (CTD). For each tetramer, the table provides the corresponding information on the CAS chemical identifier, chemical name, the heavy metal present in the chemical, NCBI gene identifier, NCBI gene name, phenotype identifier, phenotype name, MESH disease identifier, and MESH disease name.

## Supplementary Figures

**Figure S1**: Workflow to filter high confidence adverse outcome pathways (AOPs) from AOP-Wiki by employing computation and manual curation in conjunction.

**Figure S2**: Directed network corresponding to the connected component (C1) in the directed arsenic-AOP network comprising 152 KEs and 243 KERs. Among the 152 KEs, 31 are categorized as MIEs (denoted as diamond), 28 are categorized as AOs (denoted as circle), and the remaining 93 are categorized as KEs (denoted as rounded square). The 83 Kes (including MIEs and AOs) associated with inorganic arsenic are marked in ‘red’. In this figure, the 152 KEs are arranged vertically according to their level of biological organization.

**Figure S3**: Directed network corresponding to the largest connected component (C1) in the arsenic-AOP network, where the KEs (including MIEs and AOs) are colored based on their betweenness centrality values. The 83 KEs (including MIEs and AOs) associated with inorganic arsenic are marked in ‘red’. In this figure, the 152 KEs are arranged vertically according to their level of biological organization.

**Figure S4**: Directed network corresponding to the largest connected component (C1) in the arsenic-AOP network, where the KEs (including MIEs and AOs) are colored based on their eccentricity values. The 83 KEs (including MIEs and AOs) associated with inorganic arsenic are marked in ‘red’. In this figure, the 152 KEs are arranged vertically according to their level of biological organization.

**Figure S5**: Directed network corresponding to the connected component (C1) in the directed cadmium-AOP network comprising 263 KEs and 422 KERs. Among the 263 KEs, 47 are categorized as MIEs (denoted as diamond), 41 are categorized as AOs (denoted as circle), and the remaining 175 are categorized as KEs (denoted as rounded square). The 156 KEs (including MIEs and AOs) associated with inorganic cadmium are marked in ‘red’. In this figure, the 263 KEs are arranged vertically according to their level of biological organization.

**Figure S6**: Directed network corresponding to the largest connected component (C1) in the cadmium-AOP network, where the KEs (including MIEs and AOs) are colored based on their betweenness centrality values. The 156 KEs (including MIEs and AOs) associated with inorganic cadmium are marked in ‘red’. In this figure, the 263 KEs are arranged vertically according to their level of biological organization.

**Figure S7**: Directed network corresponding to the largest connected component (C1) in the cadmium-AOP network, where the KEs (including MIEs and AOs) are colored based on their eccentricity values. The 156 KEs (including MIEs and AOs) associated with inorganic cadmium are marked in ‘red’. In this figure, the 263 KEs are arranged vertically according to their level of biological organization.

**Figure S8**: The plots of SSD for Arsenic oxide (CAS:1327-53-3) based on acute toxicity data and computed using the best-fit Weibull model. **(a)** As determined by US EPA SSD Toolbox where the HC05 value is denoted by cyan colored diamond. **(b)** As determined by ssdtools where the HC05 value is denoted by a dotted line.

**Figure S9**: The plots of SSD for Disodium arsenate (CAS:7778-43-0) based on acute toxicity data and computed using the best-fit Weibull model. **(a)** As determined by US EPA SSD Toolbox where the HC05 value is denoted by cyan colored diamond. **(b)** As determined by ssdtools where the HC05 value is denoted by a dotted line.

**Figure S10**: The plots of SSD for Sodium arsenite (CAS:7784-46-5) based on acute toxicity data and computed using the best-fit BurrIII model. **(a)** As determined by US EPA SSD Toolbox where the HC05 value is denoted by cyan colored diamond. **(b)** As determined by ssdtools where the HC05 value is denoted by a dotted line.

**Figure S11**: The plots of SSD for Trisodium arsenate (CAS:13464-38-5) based on acute toxicity data and computed using the best-fit Weibull model. **(a)** As determined by US EPA SSD Toolbox where the HC05 value is denoted by cyan colored diamond. **(b)** As determined by ssdtools where the HC05 value is denoted by a dotted line.

**Figure S12**: The plots of SSD for Cadmium (CAS:7440-43-9) based on acute toxicity data and computed using the best-fit Weibull model. **(a)** As determined by US EPA SSD Toolbox where the HC05 value is denoted by cyan colored diamond. **(b)** As determined by ssdtools where the HC05 value is denoted by a dotted line.

**Figure S13**: The plots of SSD for Cadmium chloride (CAS:10108-64-2) based on acute toxicity data and computed using the best-fit Log-Logistic model. **(a)** As determined by US EPA SSD Toolbox where the HC05 value is denoted by cyan colored diamond. **(b)** As determined by ssdtools where the HC05 value is denoted by a dotted line.

**Figure S14**: The plots of SSD for Cadmium sulfate (CAS:10124-36-4) based on acute toxicity data and computed using the best-fit Log-Normal model. **(a)** As determined by US EPA SSD Toolbox where the HC05 value is denoted by cyan colored diamond. **(b)** As determined by ssdtools where the HC05 value is denoted by a dotted line.

**Figure S15**: The plots of SSD for Cadmium nitrate (CAS:10325-94-7) based on acute toxicity data and computed using the best-fit Weibull model. **(a)** As determined by US EPA SSD Toolbox where the HC05 value is denoted by cyan colored diamond. **(b)** As determined by ssdtools where the HC05 value is denoted by a dotted line.

**Figure S16**: The plots of toxicity-normalized SSD (SSDn) for Arsenic based on the acute toxicity data and computed using the best-fit BurrIII model. **(a)** As determined by US EPA SSD Toolbox where the HC05 value is denoted by cyan colored diamond. **(b)** As determined by ssdtools where the HC05 value is denoted by a dotted line.

**Figure S17**: The plots of toxicity-normalized SSD (SSDn) for Cadmium based on the acute toxicity data and computed using the best-fit Log-Logistic model. **(a)** As determined by US EPA SSD Toolbox where the HC05 value is denoted by cyan colored diamond. **(b)** As determined by ssdtools where the HC05 value is denoted by a dotted line.

**Figure S18**: The plots of SSD for Arsenic oxide (CAS:1327-53-3) based on chronic toxicity data and computed using the best-fit Log-Normal model. **(a)** As determined by US EPA SSD Toolbox where the HC05 value is denoted by cyan colored diamond. **(b)** As determined by ssdtools where the HC05 value is denoted by a dotted line.

**Figure S19**: The plots of SSD for Disodium arsenate (CAS:7778-43-0) based on chronic toxicity data and computed using the best-fit Log-Normal model. **(a)** As determined by US EPA SSD Toolbox where the HC05 value is denoted by cyan colored diamond. **(b)** As determined by ssdtools where the HC05 value is denoted by a dotted line.

**Figure S20**: The plots of SSD for Sodium arsenite (CAS:7784-46-5) based on chronic toxicity data and computed using the best-fit BurrIII model. **(a)** As determined by US EPA SSD Toolbox where the HC05 value is denoted by cyan colored diamond. **(b)** As determined by ssdtools where the HC05 value is denoted by a dotted line.

**Figure S21**: The plots of SSD for Cadmium (CAS:7440-43-9) based on chronic toxicity data and computed using the best-fit Log-Logistic model. **(a)** As determined by US EPA SSD Toolbox where the HC05 value is denoted by cyan colored diamond. **(b)** As determined by ssdtools where the HC05 value is denoted by a dotted line.

**Figure S22**: The plots of SSD for Cadmium chloride (CAS:10108-64-2) based on chronic toxicity data and computed using the best-fit BurrIII model. **(a)** As determined by US EPA SSD Toolbox where the HC05 value is denoted by cyan colored diamond. **(b)** As determined by ssdtools where the HC05 value is denoted by a dotted line.

**Figure S23**: The plots of SSD for Cadmium sulfate (CAS:10124-36-4) based on chronic toxicity data and computed using the best-fit Log-Normal model. **(a)** As determined by US EPA SSD Toolbox where the HC05 value is denoted by cyan colored diamond. **(b)** As determined by ssdtools where the HC05 value is denoted by a dotted line.

**Figure S24**: The plots of SSD for Cadmium nitrate (CAS:10325-94-7) based on chronic toxicity data and computed using the best-fit Weibull model. **(a)** As determined by US EPA SSD Toolbox where the HC05 value is denoted by cyan colored diamond. **(b)** As determined by ssdtools where the HC05 value is denoted by a dotted line.

**Figure S25**: The plots of toxicity-normalized SSD (SSDn) for Arsenic based on the chronic toxicity data and computed using the best-fit Log-Logistic model. **(a)** As determined by US EPA SSD Toolbox where the HC05 value is denoted by cyan colored diamond. **(b)** As determined by ssdtools where the HC05 value is denoted by a dotted line.

**Figure S26**: The plots of toxicity-normalized SSD (SSDn) for Cadmium based on the chronic toxicity data and computed using the best-fit Log-Logistic model. **(a)** As determined by US EPA SSD Toolbox where the HC05 value is denoted by cyan colored diamond. **(b)** As determined by ssdtools where the HC05 value is denoted by a dotted line.

