## Supplementary Text and Supplementary Figures for "Network toxicology focused investigation on the impacts of inorganic arsenic and cadmium on human and ecosystem health"

#### for

### Supplementary Text

#### 1. Compilation of AOPs within AOP-Wiki

Adverse Outcome Pathway (AOP) is a toxicological knowledge framework that captures biological processes underlying stressor-induced toxicity. In this framework, toxicity-related biological events, referred to as Key Events (KEs), are organized sequentially, beginning with the interaction of the stressor with a biological target, known as the Molecular Initiating Event (MIE), and culminating in an Adverse Outcome (AO) [1–4]. The causal, directional relationships between these events are termed Key Event Relationships (KERs) [1–4]. AOPs are stressor-agnostic and context-specific, meaning each AOP is tailored to a particular biological scenario [1–4]. Developed globally, these AOPs are deposited in the AOP-Wiki [5], which is the largest, publicly accessible repository hosted by the Society for the Advancement of Adverse Outcome Pathways (SAAOP). AOP-Wiki hosts several AOPs, where each AOP is documented in the form of KEs (including MIE and AO) and KERs, all of which are supported by scientific evidence [6]. Therefore, in this study, we relied on AOP-Wiki to obtain the latest available AOPs.

We first downloaded the XML file (released April 1, 2024) from the 'Project Downloads' page in AOP-Wiki to obtain the latest information on AOPs. Then, we parsed and extracted information associated with AOPs like AOP identifier, AOP title, associated KEs (including MIEs and AOs) and KERs, linked stressors, handbook version followed for development of AOP, status according to OECD, and biological applicability information such as taxonomy, sex and life-stage of the organism, and their corresponding weight of evidence, using an in-house Python script. Additionally, for each KE, we extracted information like KE title, KE identifier, level of biological organization, action name, object name, object identifiers and process name, and information associated with KERs like upstream/downstream KEs, evidence for biological plausibility of KER, adjacency, and the extent of quantitative understanding of KER.

#### 2. Identification of 'high confidence' AOPs within AOP-Wiki

AOPs are considered living documents as they are collaboratively developed globally, and are continuously updated based on new scientific evidence [6]. Consequently, many AOPs may remain incomplete, or lack sufficient information for further analysis [6]. Therefore, based on our previous studies [7,8], we systematically checked each AOP to filter high quality and complete AOPs (Figure S1). First, we manually checked and removed AOPs that comprised KEs with title as 'unknown', or lacked any KEs or KERs. Next, we employed NetworkX library [9] in Python to check for disconnected AOPs, and manually updated and filtered out AOPs that contained disconnected components. Finally, we checked for the presence of MIE, AO, and a directed path between them, and filtered out AOPs that lacked any. Through this combined manual and computational efforts, we identified 342 complete, connected and high quality AOPs from AOP-Wiki (last accessed on July 9, 2024), which we designated as 'high confidence AOPs' (Table S2). These 342 high confidence AOPs comprised 1163 unique KEs (Table S3) and 1827 unique KERs (Table S4).

#### 3. Computation of cumulative weight of evidence (WoE) of AOPs

AOP-Wiki provides a weight of evidence (WoE) information for each KER in an AOP based on its biological plausibility (Table S4). The WoE is a qualitative score represented in terms of 'High', 'Moderate', 'Low' and 'Not Specified'. Following Ravichandran *et al.* [10], we computed the fraction of KERs within an AOP with 'High' WoE [represented as F(High)], 'Moderate' WoE [represented as F(Moderate)], 'Low' WoE [represented as F(Low)] and 'Not Specified' WoE [represented as F(Not Specified)], and subsequently assigned a cumulative WoE to the AOPs based on the following criteria [10]:

i. If F(High) ≥ 0.5, the cumulative WoE of AOP is 'High'

ii. If F(High) < 0.5, but (F(High) + F(Moderate)) ≥ 0.5, the cumulative WoE of AOP is 'Moderate'

iii. If (F(High) + F(Moderate)) < 0.5, but (F(High) + F(Moderate) + F(Low)) ≥ 0.5, the cumulative WoE of AOP is 'Low'

iv. If none of the above criteria are satisfied, the cumulative WoE of AOP is 'Not Specified'

Tables S13-S14 provide the cumulative WoE of the AOPs associated with inorganic arsenic and cadmium, respectively.

#### 4. Identification of KEs associated with inorganic arsenic and cadmium

In this study, we aimed to comprehensively investigate the toxicities induced by inorganic arsenic and cadmium on human and ecosystem health using the AOP framework. Based on our previous studies [7,8,11], we utilized toxicogenomics and biological endpoint data related to inorganic arsenic and cadmium compounds from ToxCast [12], Comparative Toxicogenomics Database (CTD) [13,14], DEDuCT [15–17], NeurotoxKb [18,19], AOP-Wiki [5], and ECOTOX [20]. We systematically integrated data from these resources to identify KEs within AOP-Wiki that are linked to inorganic arsenic- and cadmium-induced toxicities.

##### 4.1. Identification of KEs using ToxCast

ToxCast is a program by the United States Environmental Protection Agency (US EPA) designed to enhance chemical toxicity predictions through *in vitro* high-throughput screening of various environmental chemicals [12]. These *in vitro* approaches provide crucial information on the associated biological processes and gene alterations triggered by stressor interactions, which can aid in identification of potential MIEs related to active chemicals

[7,8,11,21–23]. In this study, we utilized the latest version of ToxCast data, ToxCast invitrodb 4.1 [24], available on the US EPA repository [25], to identify KEs associated with inorganic arsenic and cadmium.

First, we extracted the chemicals and their corresponding assay information from the 'mc5-6_winning_model_fits-flags_invitrodb_v4_1_SEPT2023.csv' file, and identified active assay endpoints for each chemical (defined as 'hitc' ≥ 0.9) [26]. Next, we determined whether these active chemicals exhibited an 'activatory' or 'inhibitory' response based on the 'top' value of the corresponding winning model from the 'mc4_all_model_fits_invitrodb_v4_1_SEPT2023.csv' file [26].

To ensure that these active endpoints are not due to non-specific activation of reported genes (referred to as 'cytotoxicity-associated burst' phenomenon) [27], we applied the following Z-score statistic proposed by Judson *et al.* [27]:

$$Z(chemical, assay) = \frac{-logAC_{50}(chemical,assay) - median[-logAC_{50}(chemical,cytotox)]}{\text{global cytotoxicity MAD}}$$

Here, 'log$AC_{50}$(chemical,assay)' is the logarithm of the $AC_{50}$ value of the chemical in the assay, 'log$AC_{50}$(chemical,cytotox)' is the logarithm of the $AC_{50}$ value of the chemical in the corresponding cytotoxicity assay, and the 'global cytotoxicity MAD' is the median of the MAD (median average deviations) of the log$AC_{50}$(chemical,cytotox) distributions across all chemicals [27]. Based on this definition, Z-scores ranging between +3 and -3 are considered indicative of cytotoxicity-associated bursts [27].

In this study, we retrieved the global cytotoxicity MAD and log$AC_{50}$(chemical,cytotox) (given by the column titled 'cytotox_median_log') from the 'cytotox_invitrodb_v4_1_SEPT2023.xlsx' file to identify the cytotoxicity-associated bursts for inorganic arsenic and cadmium compounds [26]. We then discarded assays with Z-scores between +3 and -3, resulting in 281 assay endpoints linked to one inorganic arsenic compound and four inorganic cadmium compounds, which we subsequently mapped to KEs within AOP-Wiki.

We first extracted the genes corresponding to these 281 assay endpoints from the 'assay_annotations_invitrodb_v4_1_SEPT2023.xlsx' file, along with assay endpoint-gene mappings from the 'assay_gene_mappings_invitrodb_v4_1_SEPT2023' file. Next, using the KE-gene annotations provided by Saarimäki *et al.* [28], we identified gene sets linked to KEs at the molecular or cellular levels of biological organization. We then mapped these KEs to ToxCast assay endpoints based on gene overlaps and manually filtered the results using the assay endpoint descriptions. Through this comprehensive toxicogenomics-based manual curation, we identified 52 KEs associated with 40 assay endpoints across one inorganic arsenic compound (CAS:1327-53-3) and two inorganic cadmium compounds (CAS:10108-64-2, CAS:10325-94-7) (Table S5).

**4.2. Identification of KEs using CTD**

The Comparative Toxicogenomics Database (CTD) is a comprehensive public resource that links environmental chemicals (C), genes (G), phenotypes (P), and diseases (D) to advance understanding of their effects on health [13,14]. This resource facilitates the construction of CGPD-tetramers, which help identify confident associations between chemicals, phenotypes, and diseases, enabling their mapping to KEs within AOP-Wiki [29,30]. Based on our previous work [7,8,11], we retrieved high confidence CGPD-tetramers associated with inorganic arsenic and cadmium compounds, and leveraged them to identify KEs within AOP-Wiki.

We accessed the CTD June 2024 release and constructed the CGPD-tetramers where we considered: (i) chemical-gene and chemical-phenotype associations with literature evidence; (ii) chemical-disease and gene-disease associations with 'marker/mechanism' or ‘marker/mechanism|therapeutic’ evidence; (iii) gene-phenotype associations with GO annotations based on only the experimental results [31]. This process resulted in a list of 17749 CGPD-tetramers comprising three inorganic arsenic compounds and three inorganic cadmium compounds, 1162 genes, 483 phenotypes and 213 diseases (Table S22). Next, we generated the immediate neighboring GO terms for the CGPD-tetramer phenotype GO terms using the GOSim package [32] in the R programming language. We then overlapped these GO terms with the process identifiers of KEs in AOP-Wiki, manually examined them, and identified 197 KEs linked to 105 phenotypes across three inorganic arsenic compounds (CAS:1327-53-3, CAS:56320-22-0, CAS:7784-34-1) and three inorganic cadmium compounds (CAS:10108-64-2, CAS:7440-43-9, CAS:10124-36-4) (Table S5). Additionally, we manually examined the disease terms and identified 92 KEs associated with 106 diseases across three inorganic arsenic compounds (CAS:1327-53-3, CAS:56320-22-0, CAS:7784-34-1) and two inorganic cadmium compounds (CAS:7440-43-9, CAS:10108-64-2) (Table S5).

**4.3. Identification of KEs using DEDuCT and NeurotoxKb**

DEDuCT is one of the largest databases compiling curated information on endocrine disrupting chemicals (EDCs) and their corresponding endocrine-mediated endpoints from published literature [15–17]. In this study, we extracted the endocrine-mediated endpoints corresponding to inorganic arsenic and cadmium compounds within DEDuCT and considered them to identify relevant KEs within AOP-Wiki. We manually inspected both the endocrine-mediated endpoints and KE titles in AOP-Wiki, and identified 76 KEs associated with 46 endocrine-mediated endpoints across two inorganic arsenic compounds (CAS:7440-38-2, CAS:1327-53-3) and three inorganic cadmium compounds (CAS:7440-43-9, CAS:10108-64-2, CAS:10325-94-7) (Table S5).

NeurotoxKb is a manually curated resource focusing on mammalian neurotoxicity endpoints associated with environmental chemicals, compiled from published literature

[18,19]. In this study, we extracted the neurotoxic endpoints corresponding to inorganic arsenic and cadmium compounds within NeurotoxKb, and considered them to identify relevant KEs within AOP-Wiki. We manually inspected both the neurotoxic endpoints and KE titles in AOP-Wiki, and identified 20 KEs associated with 13 neurotoxic endpoints across one inorganic arsenic compound (CAS:7440-38-2) and two inorganic cadmium compounds (CAS:7440-43-9, CAS:10108-64-2) (Table S5).

##### 4.4. Identification of KEs using AOP-Wiki

AOP-Wiki catalogs information on prototypical stressors for each AOP, based on well-documented associations with these stressors [6]. In this study, we first extracted information on prototypical stressors associated with AOPs from the flat download file using an in-house Python script. We then identified that 4 AOPs (AOP:263, AOP:413, AOP:499, AOP:500) were associated with two inorganic arsenic compounds (CAS:7440-38-2, CAS:15502-74-6), and 4 AOPs (AOP:257, AOP:296, AOP:499, AOP:500) with two inorganic cadmium compounds (CAS:7440-43-9, CAS:10108-64-2). Therefore, we considered the KEs within these AOPs as associated with inorganic arsenic and cadmium, respectively.

##### 4.5. Identification of KEs using ECOTOX

ECOTOX is a comprehensive US EPA resource that consolidates toxicological information for over 12000 chemicals across more than 13000 terrestrial and aquatic species from published literature, offering valuable insights into ecological impacts of these chemicals [20]. Therefore, we here leveraged ECOTOX to identify ecotoxicologically-relevant KEs within AOP-Wiki associated with inorganic arsenic and cadmium compounds.

We first downloaded the ECOTOX dataset files (released June 2024) by selecting the 'Download ASCII Data' option on the ECOTOX website [33]. Then, using an in-house

Python script, we parsed 'tests.txt', 'results.txt', 'chemicals.txt' and 'species.txt' files to extract data associated with inorganic arsenic and cadmium compounds. ECOTOX curates the biological effects associated with the chemicals as 'Effect', the corresponding measure as 'Measurement', and the trend of the effect as 'Trend'. We removed the data points where the Measurement value was either 'Not Reported' or empty, as they lack sufficient information on adverse effects, and proceeded to map the remaining data to KEs within AOP-Wiki. We manually inspected the Effect, Measurement and Trend information from ECOTOX, and KE titles within AOP-Wiki, and identified 273 KEs to be associated with 499 unique ECOTOX endpoints across 25 inorganic arsenic compounds (CAS:7784-46-5, CAS:7778-43-0, CAS:1327-53-3, CAS:13464-37-4, CAS:1303-11-3, CAS:7440-38-2, CAS:7631-89-2, CAS:7778-39-4, CAS:'10103-60-3, CAS:13466-06-3, CAS:7784-41-0, CAS:7778-44-1, CAS:15584-04-0, CAS:15502-74-6, CAS:69162-99-8, CAS:13464-38-5, CAS:22541-54-4, CAS:17428-41-0, CAS:1303-33-9, CAS:1303-28-2, CAS:7784-34-1, CAS:7778-41-8, CAS:17029-22-0, CAS:15120-17-9, CAS:12777-38-7) and 8 inorganic cadmium compounds (CAS:10108-64-2, CAS:7440-43-9, CAS:7790-80-9, CAS:10325-94-7, CAS:10124-36-4, CAS:10022-68-1, CAS:1306-19-0, CAS:1306-23-6) (Table S5).

Overall, we identified 579 KEs that are associated with either inorganic arsenic or cadmium compounds through the integration of heterogenous toxicogenomics and biological endpoints data from six exposome-relevant resources, namely, ToxCast, CTD, DEDuCT, NeurotoxKb, AOP-Wiki and ECOTOX (Table S5).

**5. Construction of stressor-AOP network**

Stressor-AOP network provides a broader perspective on impacts of stressors across diverse biological processes by linking the stressors to AOPs [8,22]. To better understand the perturbances caused by inorganic arsenic and cadmium, we constructed a stressor-AOP network which is a bipartite graph that linked arsenic and cadmium to different AOPs within

AOP-Wiki. In order to obtain high confidence associations between the chemicals and AOPs, we relied only on the curated list of 342 high confidence AOPs (Table S2).

We first grouped KEs associated with various inorganic arsenic compounds to identify common KEs related to inorganic arsenic. Similarly, we grouped KEs for different inorganic cadmium compounds to identify those associated with inorganic cadmium. Next, we linked arsenic and cadmium to any AOP if they shared at least one common KE. Thereafter, we characterized these links based on two criteria namely, the coverage score and the level of relevance. The coverage score of a stressor-AOP link is defined as the ratio of number of KEs within that AOP associated with the stressor to the total number of KEs within that AOP [34]. This score is a real valued number between 0 and 1, and we denote this score as the edge weight of linkage between a stressor and an AOP in our stressor-AOP network. The level of relevance is a qualitative score used to identify the relevance of stressor-AOP association within the stressor-AOP network and we denote this score as an attribute of the edge in our stressor-AOP network [8]. Level of relevance is a five-level criterion defined as follows:

- *Level 1*: The stressor is associated with at least one KE within an AOP, where the KE is neither MIE nor AO within that AOP
- *Level 2*: The stressor is associated with at least one AO within an AOP, but not associated with any MIE within that AOP
- *Level 3*: The stressor is associated with at least one MIE within an AOP, but not associated with any AO within that AOP
- *Level 4*: The stressor is associated with at least one MIE and one AO within an AOP
- *Level 5*: The stressor is associated with at least one MIE and one AO within an AOP and there exists a directed path between the associated MIE and AO

Table S6 contains all the data on the stressor-AOP network constructed for inorganic arsenic and cadmium, including the coverage score and level of relevance for each of the stressor-AOP links.

#### 6. Construction of Species Sensitivity Distribution (SSD) for inorganic compounds of arsenic and cadmium

Species Sensitivity Distribution (SSD) is a widely used statistical tool that aids in ecotoxicological risk assessments [35–37]. SSD utilizes statistical distributions to provide a threshold concentration for the stressor (HC05) which is hazardous to 5% of species and not harmful to the other 95% of species in a particular environment [38–40]. We relied on the steps outlined in Sahoo *et al.* [11], to curate relevant data from ECOTOX, and construct SSDs for inorganic arsenic and cadmium compounds. First, we filtered the data to obtain species with habitat as 'Water', and standardized the concentrations to ppm equivalents for the inorganic compounds of arsenic and cadmium. To obtain relevant acute toxicity data, we considered only $LC_{50}$ or $EC_{50}$ values obtained from studies ranging between 24 and 96 hours [41], and calculated the geometric mean if multiple values were available per stressor-species pair [42,43]. To obtain relevant chronic toxicity data, we considered only NOEC values, and calculated the geometric mean if multiple values were available per stressor-species pair. Finally, to ensure robust SSDs, we included only chemicals with data across at least five ECOTOX species groups [44]. Through this extensive process, we identified acute toxicity data for four inorganic arsenic compounds (CAS:1327-53-3, CAS:7778-43-0, CAS:7784-46-5, CAS:13464-38-5), and four inorganic cadmium compounds (CAS:7440-43-9, CAS:10108-64-2, CAS:10124-36-4, CAS:10325-94-7) (Table S10). Furthermore, we identified that chronic toxicity data is available for three inorganic arsenic compounds (CAS:1327-53-3, CAS:7778-43-0, CAS:7784-46-5) and four inorganic cadmium compounds (CAS:7440-43-9, CAS:10108-64-2, CAS:10124-36-4, CAS:10325-94-7) (Table S11). We utilized this data to construct SSDs based on acute and chronic toxicities associated with inorganic arsenic and cadmium.

To construct the SSDS for inorganic compounds of arsenic and cadmium, we relied on two tools namely, SSD Toolbox developed by the US EPA [45] and R-based ssdtools package developed by the Ministry of Environment and Climate Change Strategy of British Columbia [43]. In both these tools, we fit five statistical distributions, namely, Log-Normal, Log-Logistic, Log-Gumbel, Weibull and Burr Type III to the curated toxicity data through maximum-likelihood method, and obtained the HC05 value based on the best-fit model determined by the minimum corrected Akaike Information Criterion (AICc) value [43,46,47]. Furthermore, we set the number of bootstrap resampling iterations as 10000 to quantify uncertainty in fitted parameters and estimate confidence intervals [43,46]. We observed that the Burr Type III model did not adequately fit the chronic toxicity data for the inorganic cadmium compound CAS:10325-94-7 in both tools. Therefore, only the remaining four statistical distributions were considered to compute the SSD for this chemical.

Recently, model averaging, which assigns weights to individual distributions to derive a weighted average, has been proposed as a more reliable method for computing SSDs, as it has been observed to produce more stable HC05 values compared to single-model estimates [43,48]. Accordingly, we additionally computed model-averaged HC05 values for each inorganic arsenic and cadmium compound using both tools.

[2] D.L. Villeneuve, D. Crump, N. Garcia-Reyero, M. Hecker, T.H. Hutchinson, C.A. LaLone, B. Landesmann, T. Lettieri, S. Munn, M. Nepelska, M.A. Ottinger, L. Vergauwen, M. Whelan, Adverse Outcome Pathway (AOP) Development I: Strategies

and Principles, Toxicological Sciences 142 (2014) 312–320. https://doi.org/10.1093/toxsci/kfu199.
[3] D.L. Villeneuve, D. Crump, N. Garcia-Reyero, M. Hecker, T.H. Hutchinson, C.A. LaLone, B. Landesmann, T. Lettieri, S. Munn, M. Nepelska, M.A. Ottinger, L. Vergauwen, M. Whelan, Adverse Outcome Pathway Development II: Best Practices, Toxicological Sciences 142 (2014) 321–330. https://doi.org/10.1093/toxsci/kfu200.
[4] OECD, Revised Guidance Document on Developing and Assessing Adverse Outcome Pathways, Series on Testing and Assessment No. 184, OECD Publishing, Paris (2017). https://one.oecd.org/document/env/jm/mono(2013)6/en/pdf (accessed March 24, 2024).
[5] AOP-Wiki: Adverse Outcome Pathway Wiki, (2024). https://aopwiki.org/ (accessed June 25, 2024).
[6] AOP-Wiki Developers' Handbook, (2024). https://aopwiki.org/handbooks/5 (accessed June 25, 2024).
[7] A.K. Sahoo, N. Chivukula, K. Ramesh, J. Singha, S.R. Marigoudar, K.V. Sharma, A. Samal, An integrative data-centric approach to derivation and characterization of an adverse outcome pathway network for cadmium-induced toxicity, Science of The Total Environment 920 (2024) 170968. https://doi.org/10.1016/j.scitotenv.2024.170968.
[8] A.K. Sahoo, N. Chivukula, S.R. Madgaonkar, K. Ramesh, S.R. Marigoudar, K.V. Sharma, A. Samal, Leveraging integrative toxicogenomic approach towards development of stressor-centric adverse outcome pathway networks for plastic additives, Archives of Toxicology 98 (2024) 3299–3321. https://doi.org/10.1007/s00204-024-03825-z.
[9] A.A. Hagberg, D.A. Schult, P.J. Swart, Exploring Network Structure, Dynamics, and Function using NetworkX, in: G. Varoquaux, T. Vaught, J. Millman (Eds.), Proceedings of the 7th Python in Science Conference (SciPy 2008), 2008: pp. 11–15.
[10] J. Ravichandran, B.S. Karthikeyan, A. Samal, Investigation of a derived adverse outcome pathway (AOP) network for endocrine-mediated perturbations, Science of The Total Environment 826 (2022) 154112. https://doi.org/10.1016/j.scitotenv.2022.154112.
[11] A.K. Sahoo, S.R. Madgaonkar, N. Chivukula, P. Karthikeyan, K. Ramesh, S.R. Marigoudar, K.V. Sharma, A. Samal, Network-based investigation of petroleum hydrocarbons-induced ecotoxicological effects and their risk assessment, bioRxiv (2024) 2024.07.18.604159. https://doi.org/10.1101/2024.07.18.604159.
[12] D.J. Dix, K.A. Houck, M.T. Martin, A.M. Richard, R.W. Setzer, R.J. Kavlock, The ToxCast Program for Prioritizing Toxicity Testing of Environmental Chemicals, Toxicological Sciences 95 (2007) 5–12. https://doi.org/10.1093/toxsci/kfl103.
[13] A.P. Davis, T.C. Wiegers, R.J. Johnson, D. Sciaky, J. Wiegers, C.J. Mattingly, Comparative Toxicogenomics Database (CTD): update 2023, Nucleic Acids Research 51 (2023) D1257–D1262. https://doi.org/10.1093/nar/gkac833.
[14] CTD: Comparative Toxicogenomics Database, (2024). https://ctdbase.org/ (accessed June 27, 2024).
[15] B.S. Karthikeyan, J. Ravichandran, K. Mohanraj, R.P. Vivek-Ananth, A. Samal, A curated knowledgebase on endocrine disrupting chemicals and their biological systems-level perturbations, Science of The Total Environment 692 (2019) 281–296. https://doi.org/10.1016/j.scitotenv.2019.07.225.
[16] B.S. Karthikeyan, J. Ravichandran, S.R. Aparna, A. Samal, DEDuCT 2.0: An updated knowledgebase and an exploration of the current regulations and guidelines from the perspective of endocrine disrupting chemicals, Chemosphere 267 (2021) 128898. https://doi.org/10.1016/j.chemosphere.2020.128898.
[17] DEDuCT: Database of Endocrine Disrupting Chemicals and Toxicity Profiles, (2020). https://cb.imsc.res.in/deduct/ (accessed June 25, 2024).

[32] H. Fröhlich, N. Speer, A. Poustka, T. Beißbarth, GOSim – an R-package for computation of information theoretic GO similarities between terms and gene products, BMC Bioinformatics 8 (2007) 166. https://doi.org/10.1186/1471-2105-8-166.
[33] US EPA ECOTOX Knowledgbase, (2024). https://cfpub.epa.gov/ecotox/ (accessed June 13, 2024).
[34] Z. Chai, C. Zhao, Y. Jin, Y. Wang, P. Zou, X. Ling, H. Yang, N. Zhou, Q. Chen, L. Sun, W. Chen, L. Ao, J. Cao, J. Liu, Generating adverse outcome pathway (AOP) of inorganic arsenic-induced adult male reproductive impairment via integration of phenotypic analysis in comparative toxicogenomics database (CTD) and AOP wiki, Toxicology and Applied Pharmacology 411 (2021) 115370. https://doi.org/10.1016/j.taap.2020.115370.
[35] S.A.L.M. Kooijman, A safety factor for LC50 values allowing for differences in sensitivity among species, Water Research 21 (1987) 269–276. https://doi.org/10.1016/0043-1354(87)90205-3.
[36] N.M. van Straalen, C.A.J. Denneman, Ecotoxicological evaluation of soil quality criteria, Ecotoxicology and Environmental Safety 18 (1989) 241–251. https://doi.org/10.1016/0147-6513(89)90018-3.
[37] M. Kamo, Species Sensitivity Distribution in Ecological Risk Assessment, in: M. Kamo (Ed.), Theories in Ecological Risk Assessment, Springer Nature Singapore, Singapore, 2023: pp. 103–134. https://doi.org/10.1007/978-981-99-0309-2_5.
[38] L. Posthuma, G.W. Suter II, T.P. Traas, eds., Species Sensitivity Distributions in Ecotoxicology, CRC Press, Boca Raton, 2001.
[39] R. Dowse, D. Tang, C.G. Palmer, B.J. Kefford, Risk assessment using the species sensitivity distribution method: Data quality versus data quantity, Environmental Toxicology and Chemistry 32 (2013) 1360–1369. https://doi.org/10.1002/etc.2190.
[40] US EPA, Guidelines for Ecological Risk Assessment, (1998). https://www.epa.gov/risk/guidelines-ecological-risk-assessment (accessed April 22, 2024).
[41] U.S. EPA, Methods for Measuring the Acute Toxicity of Effluents and Receiving Waters to Freshwater and Marine Organisms. Fifth Edition, U.S. Environmental Protection Agency, 2002. https://www.epa.gov/risk/guidelines-ecological-risk-assessment (accessed April 22, 2024).
[42] J.R. Wheeler, E.P.M. Grist, K.M.Y. Leung, D. Morritt, M. Crane, Species sensitivity distributions: data and model choice, Marine Pollution Bulletin 45 (2002) 192–202. https://doi.org/10.1016/S0025-326X(01)00327-7.
[43] C. Schwarz, A. Tillmanns, Improving statistical methods to derive species sensitivity distribution, Province of British Columbia, Victoria, 2019. https://a100.gov.bc.ca/pub/acat/public/viewReport.do?reportId=57400.
[44] S. Belanger, M. Barron, P. Craig, S. Dyer, M. Galay-Burgos, M. Hamer, S. Marshall, L. Posthuma, S. Raimondo, P. Whitehouse, Future needs and recommendations in the development of species sensitivity distributions: Estimating toxicity thresholds for aquatic ecological communities and assessing impacts of chemical exposures, Integrated Environmental Assessment and Management 13 (2017) 664–674. https://doi.org/10.1002/ieam.1841.
[45] US EPA Species Sensitivity Distribution (SSD) Toolbox, (2024). https://www.epa.gov/comptox-tools/species-sensitivity-distribution-ssd-toolbox (accessed May 10, 2024).
[46] M. Etterson, Technical Manual: SSD Toolbox Version 1.0, U.S. Environmental Protection Agency, 2020.

[47] J.E. Cavanaugh, A.A. Neath, The Akaike information criterion: Background, derivation, properties, application, interpretation, and refinements, WIREs Computational Statistics 11 (2019) e1460. https://doi.org/10.1002/wics.1460.
[48] D.R. Fox, R.A. van Dam, R. Fisher, G.E. Batley, A.R. Tillmanns, J. Thorley, C.J. Schwarz, D.J. Spry, K. McTavish, Recent Developments in Species Sensitivity Distribution Modeling, Environmental Toxicology and Chemistry 40 (2021) 293–308. https://doi.org/10.1002/etc.4925.

### Supplementary Figures

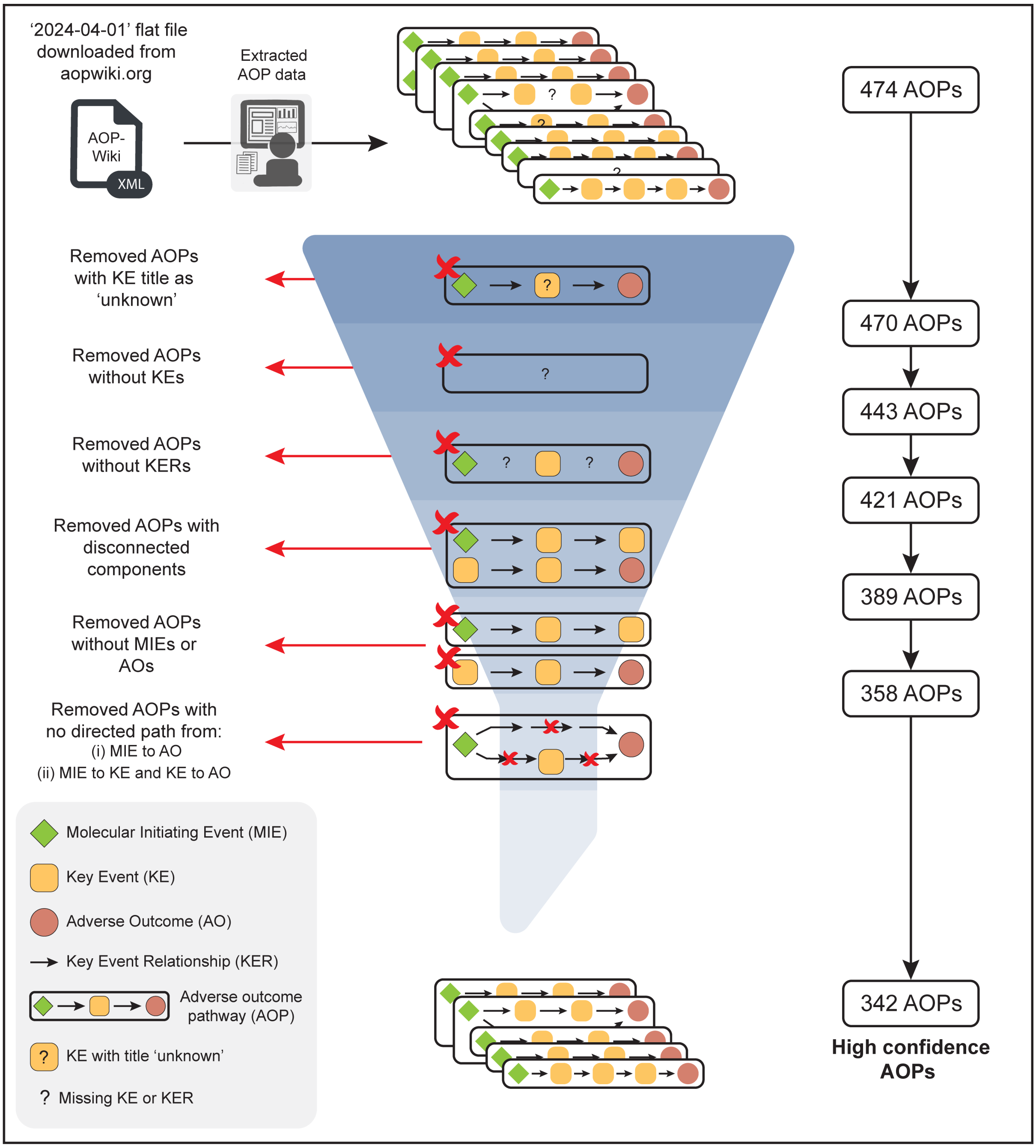


**Figure S1:** Workflow to filter high confidence adverse outcome pathways (AOPs) from AOP- Wiki by employing computation and manual curation in conjunction.

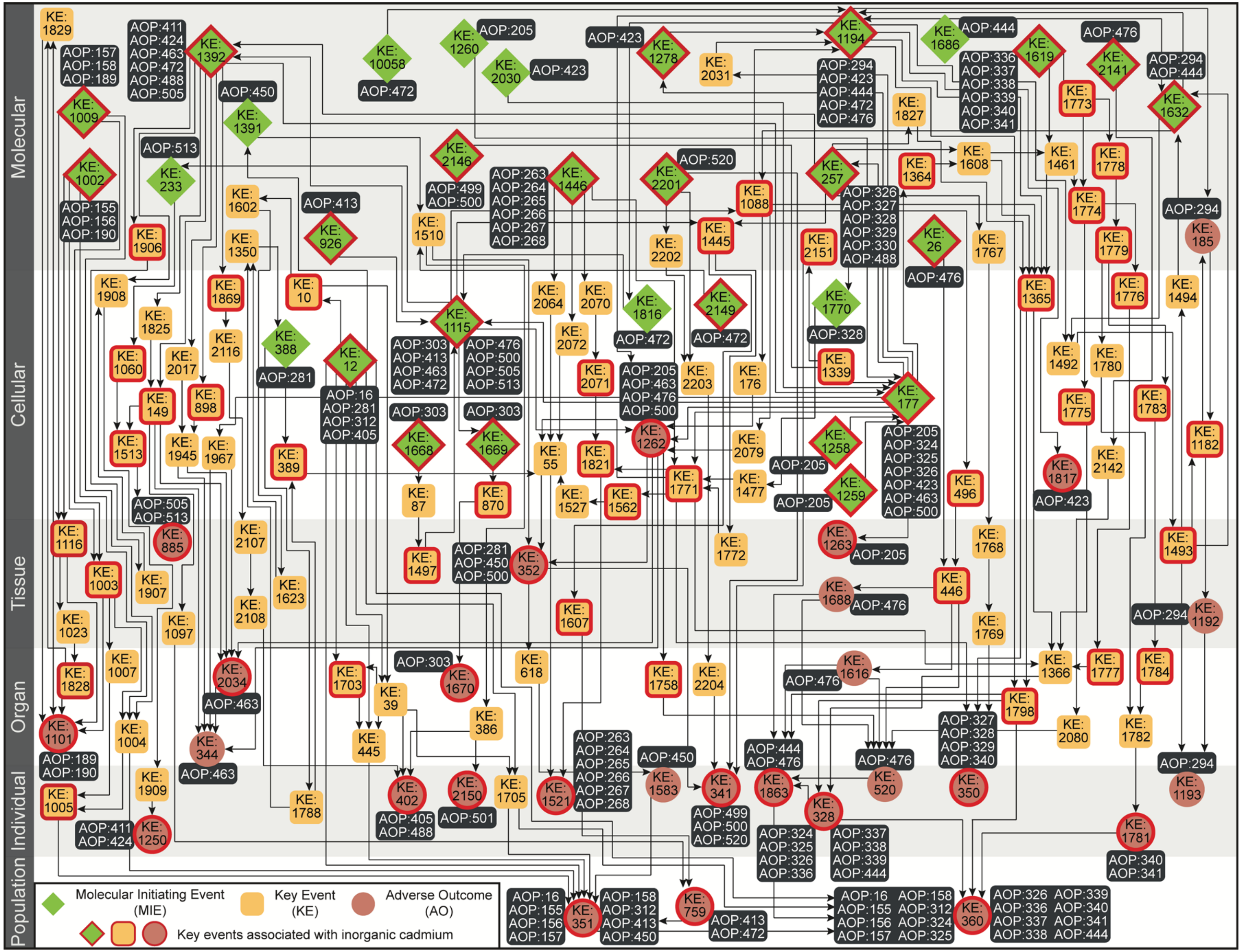
Molecular
Cellular
Tissue
Organ
Individual
Population
KE: 1829
AOP:157 AOP:158 AOP:189
AOP:411 AOP:424 AOP:463 AOP:472 AOP:488 AOP:505
KE: 1392
KE: 1009
KE: 1002
AOP:155 AOP:156 AOP:190
AOP:513
KE: 233
AOP:450
KE: 1391
KE: 10058
AOP:472
KE: 1260
AOP:205
KE: 2030
AOP:423
AOP:423
KE: 1278
KE: 2031
KE: 1194
AOP:294 AOP:423 AOP:444 AOP:472 AOP:476
KE: 1686
AOP:444
AOP:336 AOP:337 AOP:338 AOP:339 AOP:340 AOP:341
KE: 1619
AOP:476
KE: 2141
AOP:294 AOP:444
KE: 1632
KE: 1773
KE: 1827
KE: 1608
KE: 1461
KE: 1778
KE: 1364
KE: 2146
AOP:499 AOP:500
AOP:263 AOP:264 AOP:265 AOP:266 AOP:267 AOP:268
KE: 1446
KE: 2201
AOP:520
KE: 1088
KE: 257
AOP:326 AOP:327 AOP:328 AOP:329 AOP:330 AOP:488
KE: 1602
KE: 1350
AOP:413
KE: 926
KE: 1510
KE: 1906
KE: 2202
KE: 1445
KE: 2151
KE: 26
AOP:476
KE: 1767
KE: 1774
KE: 1779
AOP:294
KE: 185
KE: 1908
KE: 1825
KE: 1869
KE: 10
KE: 1115
KE: 2064
KE: 2070
KE: 1816
AOP:472
KE: 2149
AOP:472
KE: 1770
AOP:328
KE: 1365
KE: 1776
KE: 1494
KE: 1060
KE: 2017
KE: 2116
KE: 388
AOP:281
KE: 12
AOP:303 AOP:413 AOP:463 AOP:472
AOP:476 AOP:500 AOP:505 AOP:513
KE: 2072
KE: 2071
AOP:205 AOP:463 AOP:476 AOP:500
KE: 2203
KE: 176
KE: 1339
KE: 177
KE: 1492
KE: 1780
KE: 149
KE: 898
AOP:16 AOP:281 AOP:312 AOP:405
AOP:303
AOP:303
KE: 1775
KE: 1783
KE: 1513
KE: 1945
KE: 1967
KE: 389
KE: 1668
KE: 1669
KE: 55
KE: 1821
KE: 1262
KE: 2079
KE: 1258
AOP:205 AOP:324 AOP:325 AOP:326 AOP:423 AOP:463 AOP:500
KE: 1817
KE: 2142
KE: 1182
AOP:505 AOP:513
KE: 87
KE: 870
KE: 1527
KE: 1562
KE: 1771
KE: 1477
AOP:205
AOP:205
KE: 1259
KE: 496
AOP:423
KE: 1116
KE: 885
KE: 2107
KE: 1497
AOP:281 AOP:450 AOP:500
KE: 352
KE: 1772
KE: 1263
AOP:205
KE: 1768
KE: 1493
KE: 1003
KE: 1907
KE: 1623
KE: 1688
AOP:476
KE: 446
KE: 1023
KE: 1097
KE: 2108
KE: 1607
KE: 1769
AOP:294
KE: 1192
KE: 1828
KE: 1007
KE: 2034
AOP:463
KE: 1703
AOP:303
KE: 1670
KE: 39
KE: 618
KE: 1758
KE: 2204
AOP:476
KE: 1616
KE: 1366
KE: 1777
KE: 1784
KE: 1101
AOP:189 AOP:190
KE: 1004
KE: 344
AOP:463
KE: 445
KE: 386
AOP:263 AOP:264 AOP:265 AOP:266 AOP:267 AOP:268
AOP:450
AOP:444 AOP:476
KE: 1798
AOP:327 AOP:328 AOP:329 AOP:340
AOP:476
KE: 2080
KE: 1782
AOP:294
KE: 1005
KE: 1909
KE: 1788
KE: 402
AOP:405 AOP:488
KE: 2150
AOP:501
KE: 1705
KE: 1521
KE: 1583
KE: 341
AOP:499 AOP:500 AOP:520
KE: 1863
KE: 328
KE: 520
KE: 350
KE: 1193
AOP:411 AOP:424
KE: 1250
AOP:324 AOP:325 AOP:326 AOP:336
AOP:337 AOP:338 AOP:339 AOP:444
KE: 1781
AOP:340 AOP:341
Molecular Initiating Event (MIE)
Key Event (KE)
Adverse Outcome (AO)
Key events associated with inorganic cadmium
AOP:16 AOP:155 AOP:156 AOP:157
KE: 351
AOP:158 AOP:312 AOP:413 AOP:450
KE: 759
AOP:413 AOP:472
AOP:16 AOP:155 AOP:156 AOP:157
AOP:158 AOP:312 AOP:324 AOP:325
KE: 360
AOP:326 AOP:336 AOP:337 AOP:338
AOP:339 AOP:340 AOP:341 AOP:444

**Figure S2**: Directed network corresponding to the connected component (C1) in the directed arsenic-AOP network comprising 152 KEs and 243 KERs. Among the 152 KEs, 31 are categorized as MIEs (denoted as diamond), 28 are categorized as AOs (denoted as circle), and the remaining 93 are categorized as KEs (denoted as rounded square). The 83 KEs (including MIEs and AOs) associated with inorganic arsenic are marked in 'red'. In this figure, the 152 KEs are arranged vertically according to their level of biological organization.

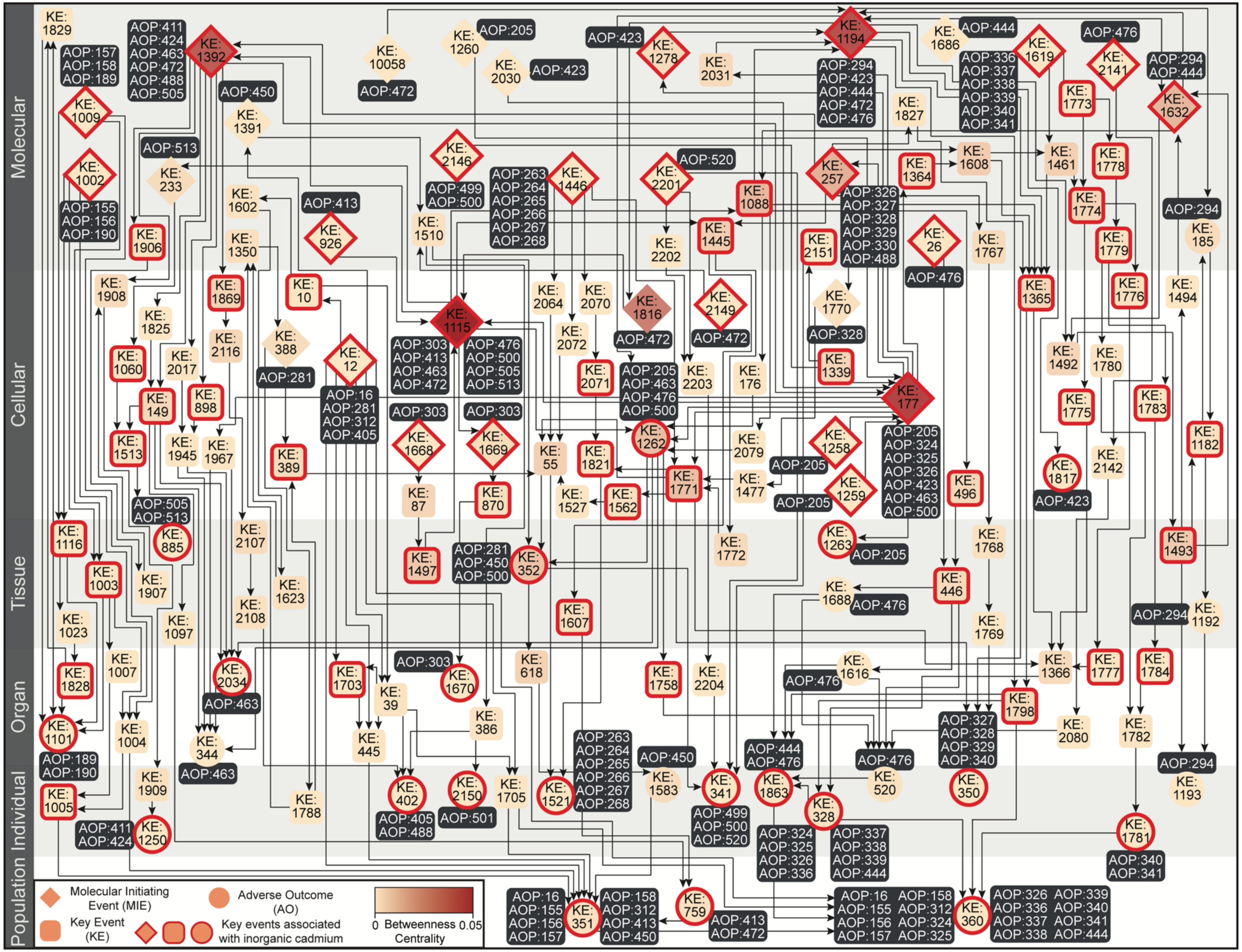

Molecular
Cellular
Tissue
Organ
Individual
Population
KE: 1829
AOP:157 AOP:158 AOP:189
AOP:411 AOP:424 AOP:463 AOP:472 AOP:488 AOP:505
KE: 1392
KE: 10058
AOP:472
KE: 1260
AOP:205
KE: 2030
AOP:423
AOP:423
KE: 1278
KE: 2031
KE: 1194
AOP:294 AOP:423 AOP:444 AOP:472 AOP:476
KE: 1686
AOP:444
AOP:336 AOP:337 AOP:338 AOP:339 AOP:340 AOP:341
KE: 1619
AOP:476
KE: 2141
AOP:294 AOP:444
KE: 1632
KE: 1773
KE: 1827
KE: 1009
AOP:450
KE: 1391
AOP:513
KE: 233
KE: 1002
AOP:155 AOP:156 AOP:190
KE: 2146
AOP:499 AOP:500
AOP:263 AOP:264 AOP:265 AOP:266 AOP:267 AOP:268
KE: 1446
KE: 2201
AOP:520
KE: 257
KE: 1364
KE: 1608
KE: 1461
KE: 1778
KE: 1088
AOP:326 AOP:327 AOP:328 AOP:329 AOP:330 AOP:488
KE: 1602
AOP:413
KE: 926
KE: 1510
KE: 1445
KE: 2202
KE: 2151
KE: 26
AOP:476
KE: 1767
KE: 1774
KE: 1779
AOP:294
KE: 185
KE: 1906
KE: 1350
KE: 1908
KE: 1869
KE: 10
KE: 1115
KE: 2064
KE: 2070
KE: 1816
AOP:472
KE: 2149
AOP:472
KE: 1770
AOP:328
KE: 1365
KE: 1776
KE: 1494
KE: 1825
KE: 2116
KE: 388
AOP:281
KE: 12
AOP:303 AOP:413 AOP:463 AOP:472
AOP:476 AOP:500 AOP:505 AOP:513
KE: 2072
KE: 1060
KE: 2017
KE: 149
KE: 898
AOP:16 AOP:281 AOP:312 AOP:405
AOP:303
AOP:303
KE: 2071
AOP:205 AOP:463 AOP:476 AOP:500
KE: 2203
KE: 176
KE: 1339
KE: 177
AOP:205 AOP:324 AOP:325 AOP:326 AOP:423 AOP:463 AOP:500
KE: 1492
KE: 1780
KE: 1775
KE: 1783
KE: 1513
KE: 1945
KE: 1967
KE: 389
KE: 1668
KE: 1669
KE: 55
KE: 1821
KE: 1262
KE: 2079
KE: 1258
AOP:205
KE: 1771
KE: 1477
KE: 1259
AOP:205
KE: 496
KE: 1817
AOP:423
KE: 2142
KE: 1182
AOP:505 AOP:513
KE: 87
KE: 870
KE: 1527
KE: 1562
KE: 1116
KE: 885
KE: 2107
KE: 1497
AOP:281 AOP:450 AOP:500
KE: 352
KE: 1772
KE: 1263
AOP:205
KE: 1768
KE: 1493
KE: 1003
KE: 1907
KE: 1623
KE: 446
KE: 1688
AOP:476
KE: 1023
KE: 1097
KE: 2108
KE: 1607
KE: 1769
AOP:294
KE: 1192
KE: 1828
KE: 1007
KE: 2034
AOP:463
KE: 1703
AOP:303
KE: 1670
KE: 618
KE: 1758
KE: 2204
AOP:476
KE: 1616
KE: 1366
KE: 1777
KE: 1784
KE: 1101
AOP:189 AOP:190
KE: 1004
KE: 344
AOP:463
KE: 39
KE: 445
KE: 386
AOP:263 AOP:264 AOP:265 AOP:266 AOP:267 AOP:268
AOP:450
AOP:444 AOP:476
AOP:476
KE: 1798
AOP:327 AOP:328 AOP:329 AOP:340
KE: 2080
KE: 1782
AOP:294
KE: 1005
KE: 1909
KE: 1788
KE: 402
AOP:405 AOP:488
KE: 2150
AOP:501
KE: 1705
KE: 1521
KE: 1583
KE: 341
AOP:499 AOP:500 AOP:520
KE: 1863
KE: 328
AOP:324 AOP:325 AOP:326 AOP:336
AOP:337 AOP:338 AOP:339 AOP:444
KE: 520
KE: 350
KE: 1193
AOP:411 AOP:424
KE: 1250
KE: 1781
AOP:340 AOP:341
Molecular Initiating Event (MIE)
Key Event (KE)
Adverse Outcome (AO)
Key events associated with inorganic cadmium
0 Betweenness 0.05 Centrality
AOP:16 AOP:155 AOP:156 AOP:157
KE: 351
AOP:158 AOP:312 AOP:413 AOP:450
KE: 759
AOP:413 AOP:472
AOP:16 AOP:155 AOP:156 AOP:157
AOP:158 AOP:312 AOP:324 AOP:325
KE: 360
AOP:326 AOP:336 AOP:337 AOP:338
AOP:339 AOP:340 AOP:341 AOP:444

**Figure S3**: Directed network corresponding to the largest connected component (C1) in the arsenic-AOP network, where the KEs (including MIEs and AOs) are colored based on their betweenness centrality values. The 83 KEs (including MIEs and AOs) associated with inorganic arsenic are marked in ‘red’. In this figure, the 152 KEs are arranged vertically according to their level of biological organization.

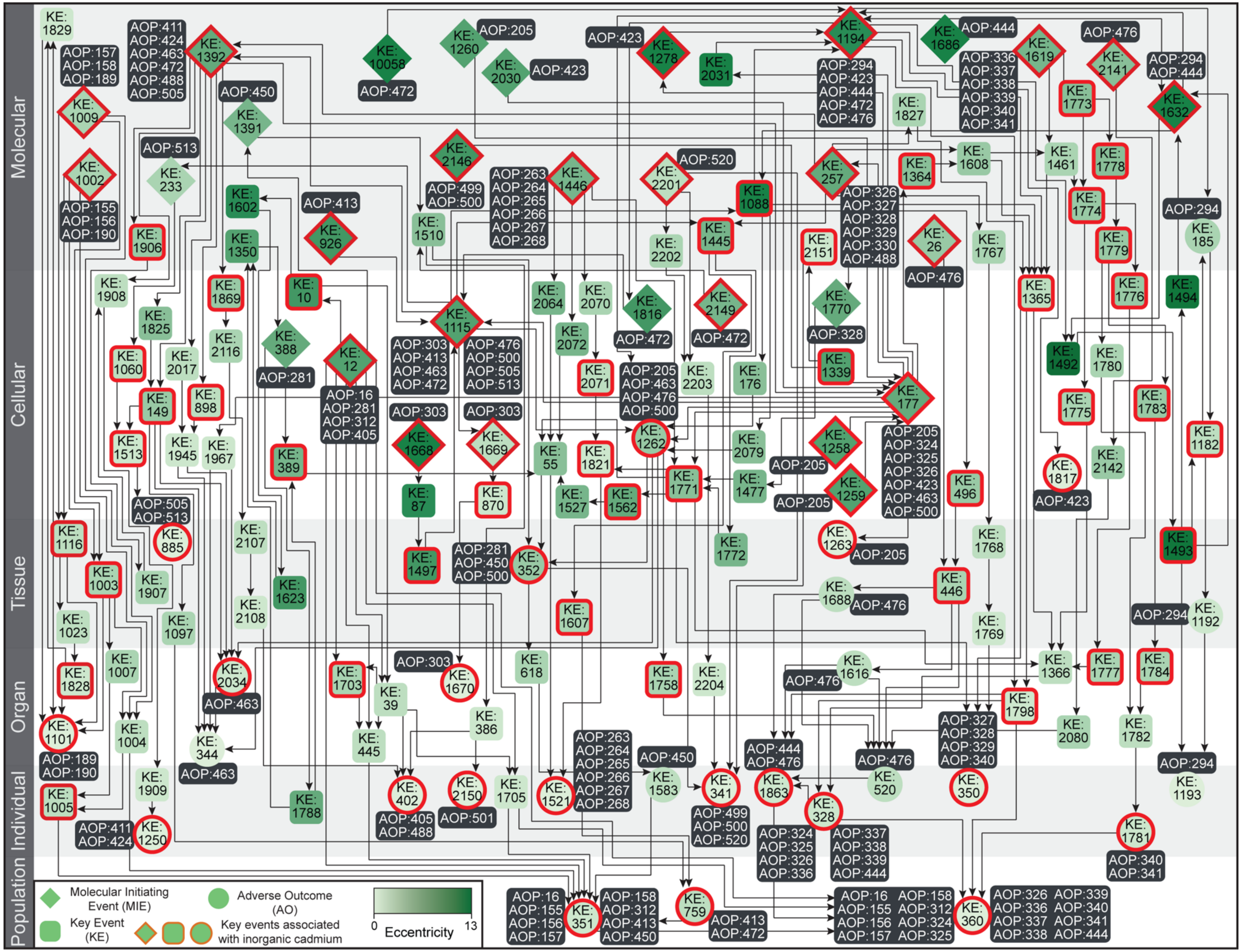

Molecular
Cellular
Tissue
Organ
Individual
Population
KE: 1829
AOP:157 AOP:158 AOP:189
AOP:411 AOP:424 AOP:463 AOP:472 AOP:488 AOP:505
KE: 1392
KE: 1009
KE: 1002
AOP:155 AOP:156 AOP:190
AOP:450
KE: 1391
KE: 10058
AOP:472
KE: 1260
AOP:205
KE: 2030
AOP:423
AOP:423
KE: 1278
KE: 2031
KE: 1194
AOP:294 AOP:423 AOP:444 AOP:472 AOP:476
KE: 1686
AOP:444
AOP:336 AOP:337 AOP:338 AOP:339 AOP:340 AOP:341
KE: 1619
AOP:476
KE: 2141
AOP:294 AOP:444
KE: 1632
KE: 1773
KE: 1827
AOP:513
KE: 233
KE: 1602
AOP:413
KE: 926
KE: 2146
AOP:499 AOP:500
AOP:263 AOP:264 AOP:265 AOP:266 AOP:267 AOP:268
KE: 1446
KE: 2201
AOP:520
KE: 1088
KE: 257
AOP:326 AOP:327 AOP:328 AOP:329 AOP:330 AOP:488
KE: 1364
KE: 1608
KE: 1461
KE: 1778
KE: 1774
KE: 1906
KE: 1350
KE: 1510
KE: 2202
KE: 1445
KE: 2151
KE: 26
AOP:476
KE: 1767
KE: 1779
AOP:294
KE: 185
KE: 1908
KE: 1869
KE: 10
KE: 1825
KE: 2116
KE: 388
AOP:281
KE: 12
AOP:16 AOP:281 AOP:312 AOP:405
KE: 1115
AOP:303 AOP:413 AOP:463 AOP:472
AOP:476 AOP:500 AOP:505 AOP:513
KE: 2064
KE: 2070
KE: 1816
AOP:472
KE: 2149
AOP:472
KE: 1770
AOP:328
KE: 1365
KE: 1776
KE: 1494
KE: 1060
KE: 2017
KE: 149
KE: 898
KE: 2072
KE: 2071
AOP:205 AOP:463 AOP:476 AOP:500
KE: 2203
KE: 176
KE: 1339
KE: 177
KE: 1492
KE: 1780
KE: 1775
KE: 1783
KE: 1513
KE: 1945
KE: 1967
KE: 389
AOP:303
KE: 1668
AOP:303
KE: 1669
KE: 55
KE: 1821
KE: 1262
KE: 2079
KE: 1258
AOP:205 AOP:324 AOP:325 AOP:326 AOP:423 AOP:463 AOP:500
KE: 1182
KE: 1817
KE: 2142
AOP:205
KE: 1771
KE: 1477
KE: 1259
AOP:205
KE: 496
AOP:423
KE: 87
KE: 870
KE: 1527
KE: 1562
AOP:505 AOP:513
KE: 885
KE: 1116
KE: 2107
KE: 1497
AOP:281 AOP:450 AOP:500
KE: 352
KE: 1772
KE: 1263
AOP:205
KE: 1768
KE: 1493
KE: 1003
KE: 1907
KE: 1623
KE: 2108
KE: 1023
KE: 1097
KE: 1607
KE: 1688
AOP:476
KE: 446
KE: 1769
AOP:294
KE: 1192
KE: 1828
KE: 1007
KE: 2034
AOP:463
KE: 1703
AOP:303
KE: 1670
KE: 618
KE: 1758
KE: 2204
AOP:476
KE: 1616
KE: 1366
KE: 1777
KE: 1784
KE: 1101
AOP:189 AOP:190
KE: 1004
KE: 344
AOP:463
KE: 39
KE: 445
KE: 386
AOP:263 AOP:264 AOP:265 AOP:266 AOP:267 AOP:268
AOP:450
AOP:444 AOP:476
AOP:476
KE: 1798
AOP:327 AOP:328 AOP:329 AOP:340
KE: 2080
KE: 1782
AOP:294
KE: 1005
KE: 1909
KE: 1788
KE: 402
AOP:405 AOP:488
KE: 2150
AOP:501
KE: 1705
KE: 1521
KE: 1583
KE: 341
AOP:499 AOP:500 AOP:520
KE: 1863
KE: 328
AOP:324 AOP:325 AOP:326 AOP:336
AOP:337 AOP:338 AOP:339 AOP:444
KE: 520
KE: 350
KE: 1193
AOP:411 AOP:424
KE: 1250
KE: 1781
AOP:340 AOP:341
AOP:16 AOP:155 AOP:156 AOP:157
KE: 351
AOP:158 AOP:312 AOP:413 AOP:450
KE: 759
AOP:413 AOP:472
AOP:16 AOP:155 AOP:156 AOP:157
AOP:158 AOP:312 AOP:324 AOP:325
KE: 360
AOP:326 AOP:336 AOP:337 AOP:338
AOP:339 AOP:340 AOP:341 AOP:444
Molecular Initiating Event (MIE)
Adverse Outcome (AO)
Key Event (KE)
Key events associated with inorganic cadmium
0
13
Eccentricity

**Figure S4**: Directed network corresponding to the largest connected component (C1) in the arsenic-AOP network, where the KEs (including MIEs and AOs) are colored based on their eccentricity values. The 83 KEs (including MIEs and AOs) associated with inorganic arsenic are marked in ‘red’. In this figure, the 152 KEs are arranged vertically according to their level of biological organization.

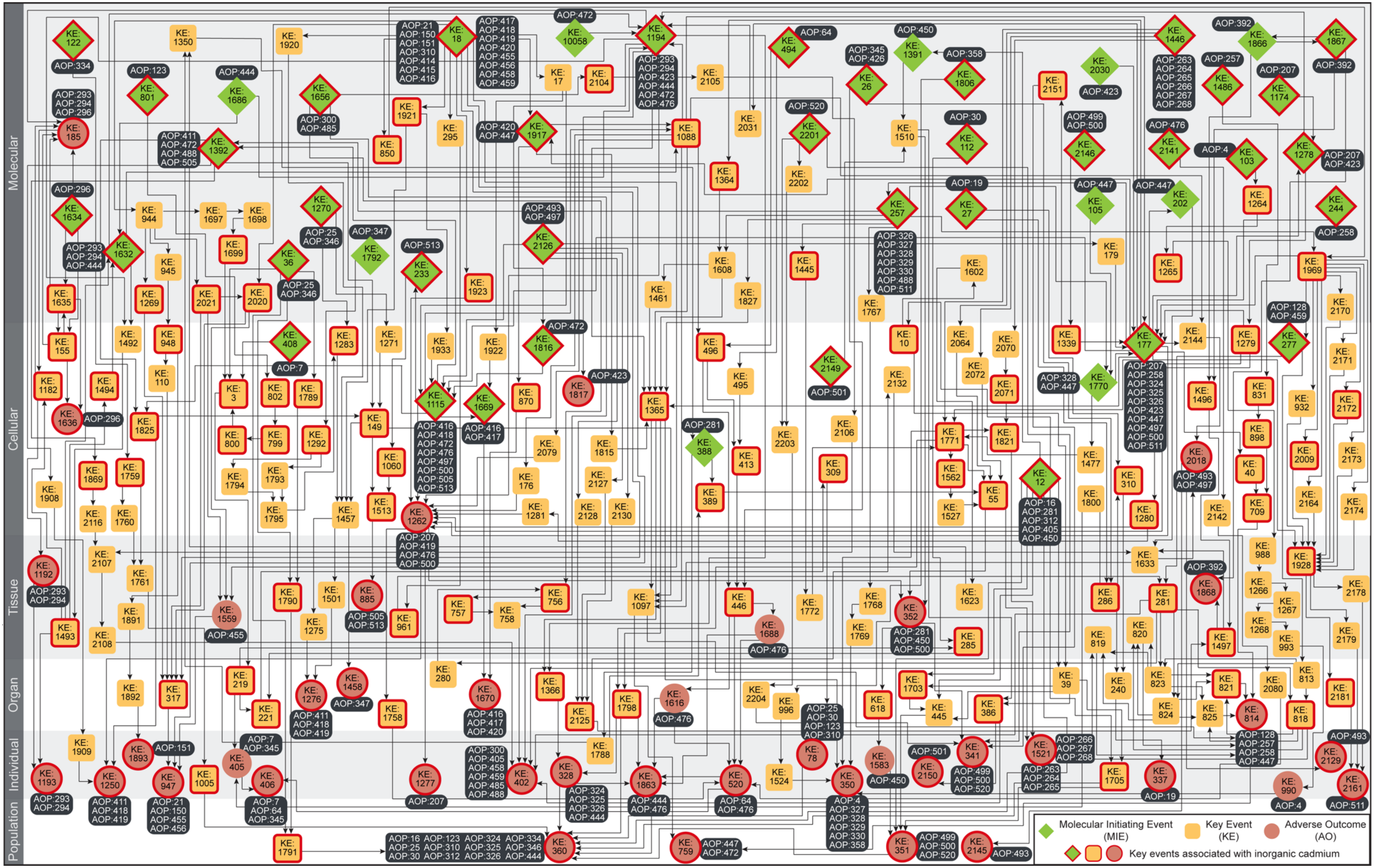
Molecular
Cellular
Tissue
Organ
Individual
Population
Molecular Initiating Event (MIE)
Key Event (KE)
Adverse Outcome (AO)
Key events associated with inorganic cadmium

**Figure S5**: Directed network corresponding to the connected component (C1) in the directed cadmium-AOP network comprising 263 KEs and 422 KERs. Among the 263 KEs, 47 are categorized as MIEs (denoted as diamond), 41 are categorized as AOs (denoted as circle), and the remaining 175 are categorized as KEs (denoted as rounded square). The 156 KEs (including MIEs and AOs) associated with inorganic cadmium are marked in 'red'. In this figure, the 263 KEs are arranged vertically according to their level of biological organization.

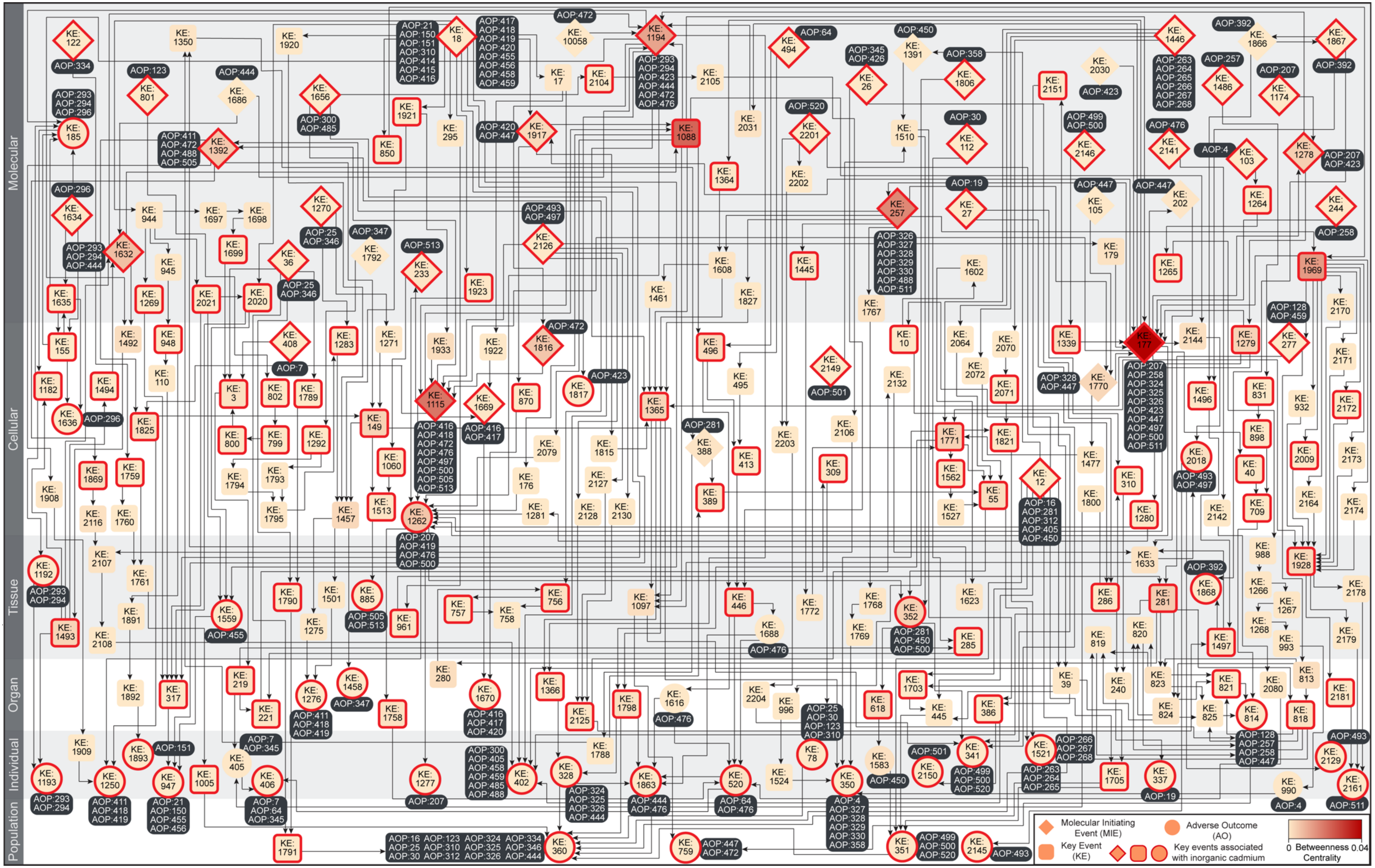
Molecular
Cellular
Tissue
Organ
Individual
Population
Molecular Initiating Event (MIE)
Adverse Outcome (AO)
Key Event (KE)
Key events associated with inorganic cadmium
0 Betweenness 0.04
Centrality

**Figure S6**: Directed network corresponding to the largest connected component (C1) in the cadmium-AOP network, where the KEs (including MIEs and AOs) are colored based on their betweenness centrality values. The 156 KEs (including MIEs and AOs) associated with inorganic cadmium are marked in 'red'. In this figure, the 263 KEs are arranged vertically according to their level of biological organization.

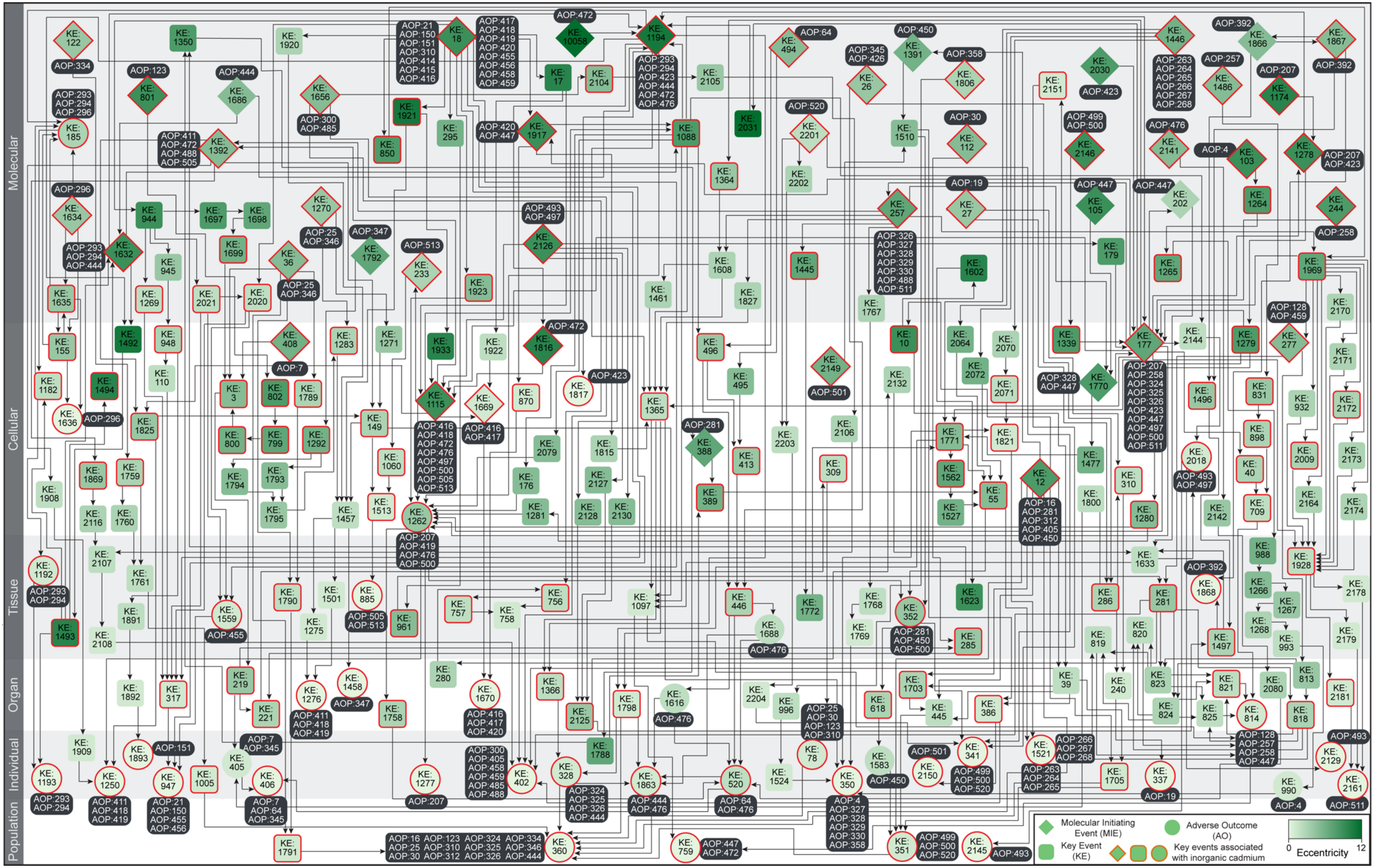
Molecular
Cellular
Tissue
Organ
Individual
Population
Molecular Initiating Event (MIE)
Adverse Outcome (AO)
Key Event (KE)
Key events associated with inorganic cadmium
Eccentricity
0
12

**Figure S7**: Directed network corresponding to the largest connected component (C1) in the cadmium-AOP network, where the KEs (including MIEs and AOs) are colored based on their eccentricity values. The 156 KEs (including MIEs and AOs) associated with inorganic cadmium are marked in ‘red’. In this figure, the 263 KEs are arranged vertically according to their level of biological organization.

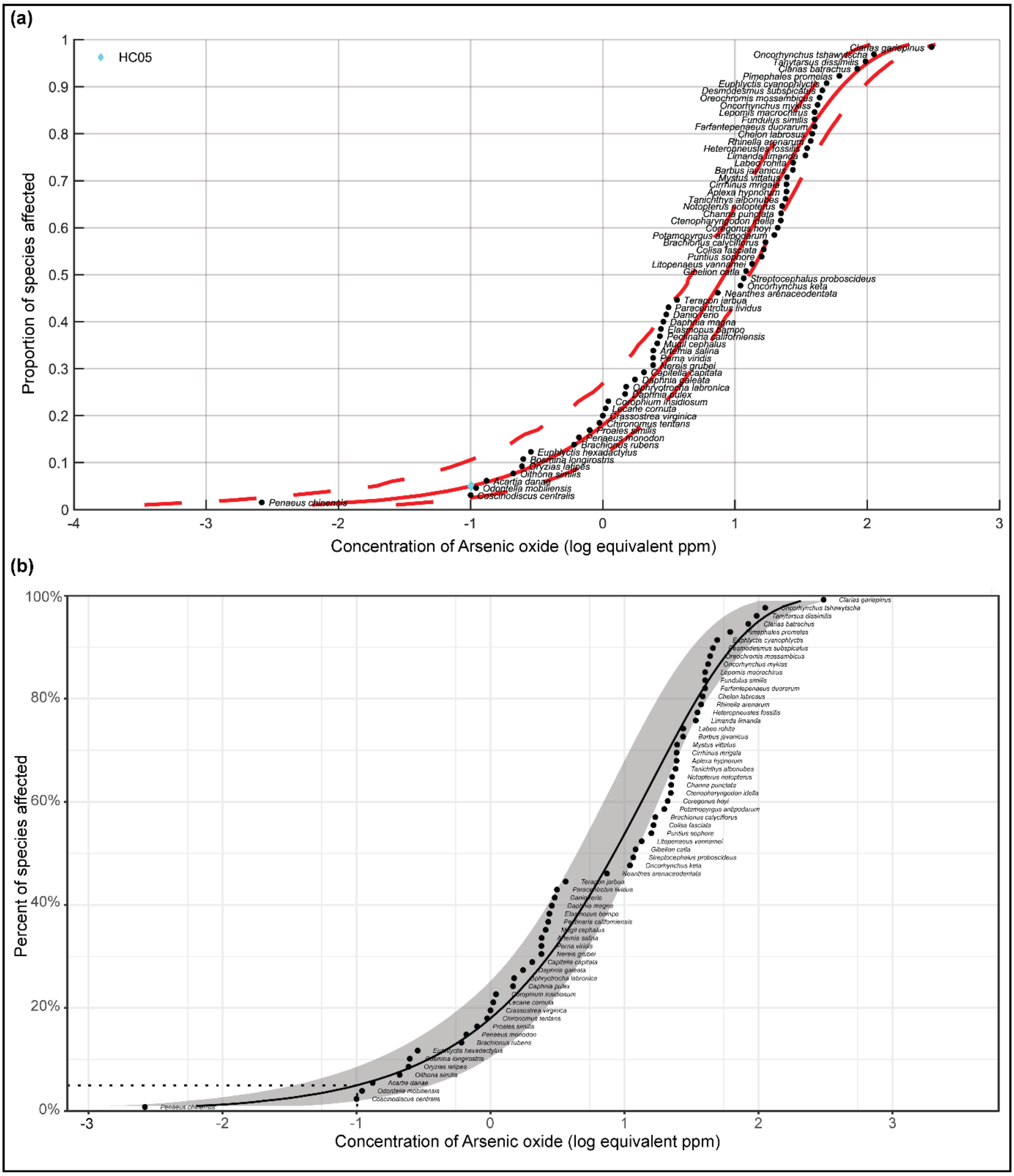


**Figure S8**: The plots of SSD for Arsenic oxide (CAS:1327-53-3) based on acute toxicity data and computed using the best-fit Weibull model. **(a)** As determined by US EPA SSD Toolbox where the HC05 value is denoted by cyan colored diamond. **(b)** As determined by ssdtools where the HC05 value is denoted by a dotted line.

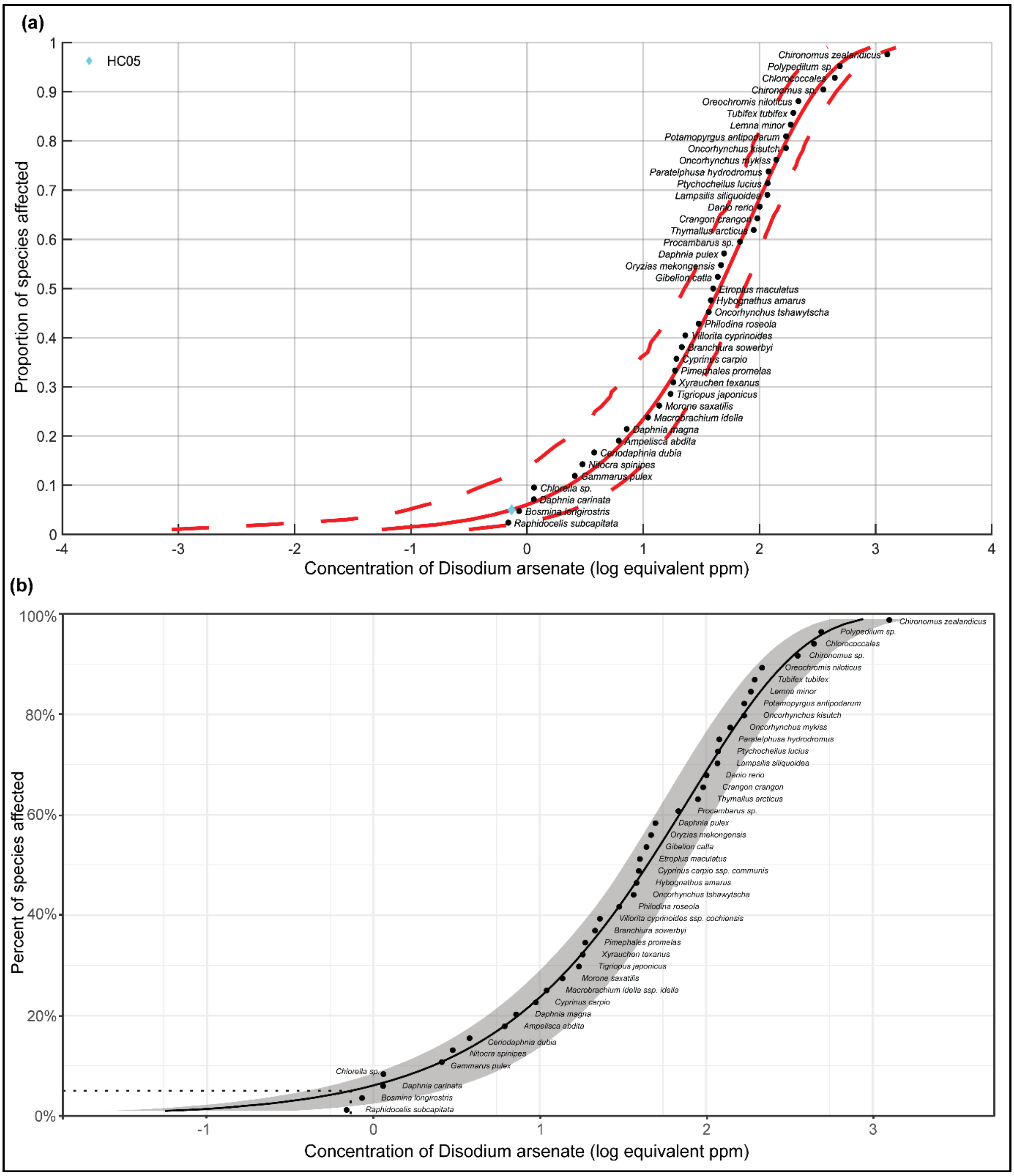


**Figure S9**: The plots of SSD for Disodium arsenate (CAS:7778-43-0) based on acute toxicity data and computed using the best-fit Weibull model. **(a)** As determined by US EPA SSD Toolbox where the HC05 value is denoted by cyan colored diamond. **(b)** As determined by ssdtools where the HC05 value is denoted by a dotted line.

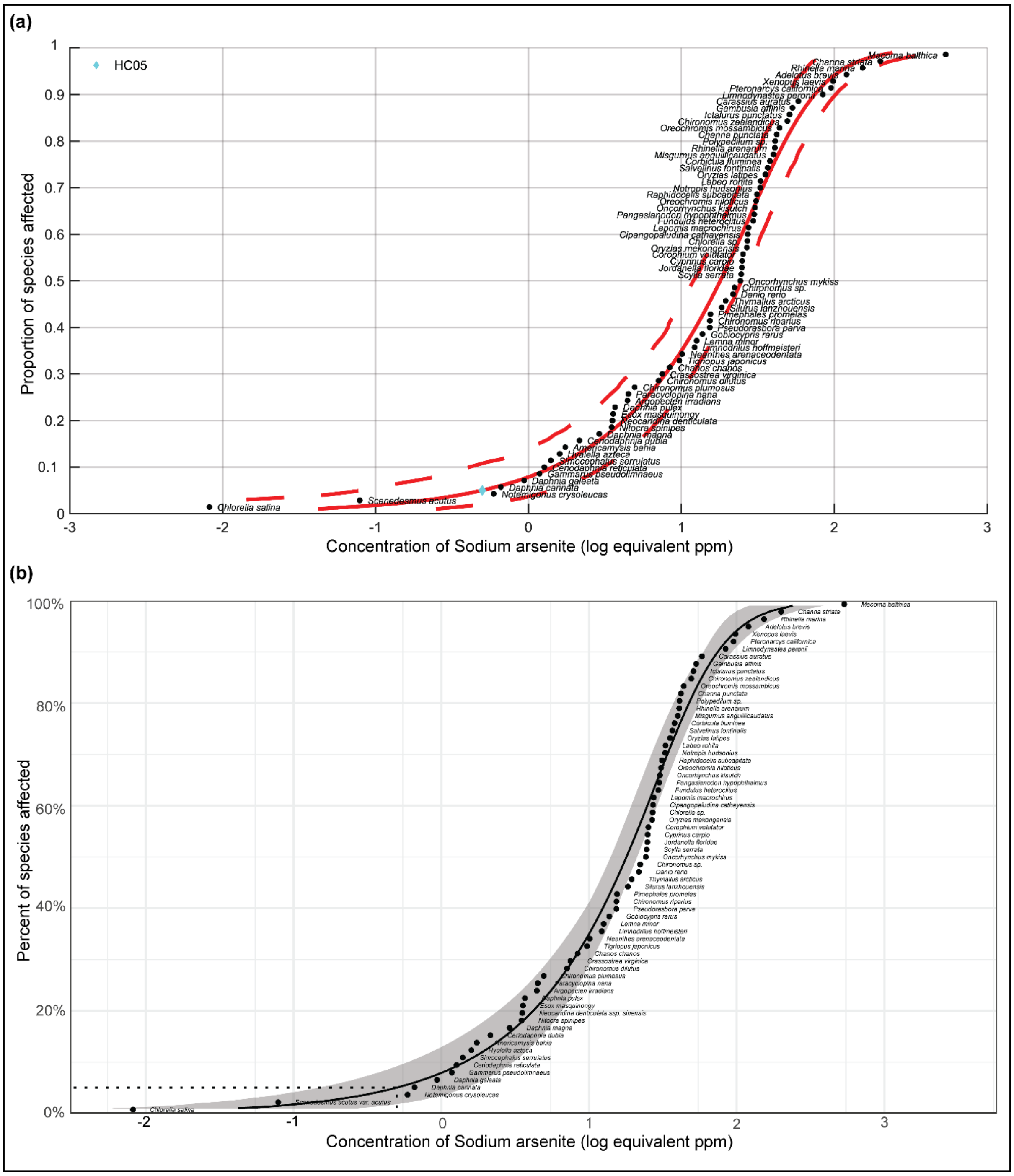


**Figure S10**: The plots of SSD for Sodium arsenite (CAS:7784-46-5) based on acute toxicity data and computed using the best-fit BurrIII model. **(a)** As determined by US EPA SSD Toolbox where the HC05 value is denoted by cyan colored diamond. **(b)** As determined by ssdtools where the HC05 value is denoted by a dotted line.

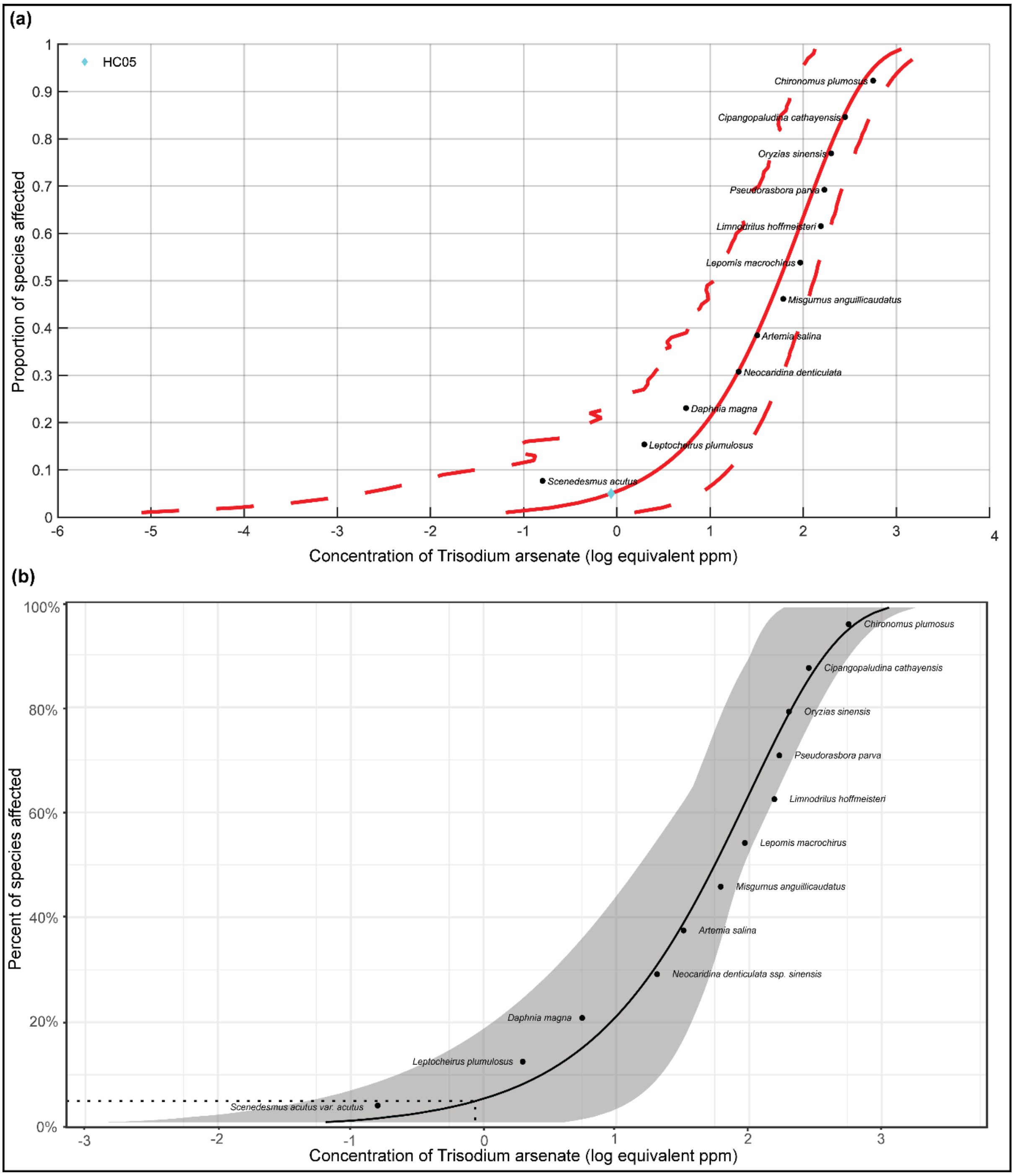


**Figure S11**: The plots of SSD for Trisodium arsenate (CAS:13464-38-5) based on acute toxicity data and computed using the best-fit Weibull model. **(a)** As determined by US EPA SSD Toolbox where the HC05 value is denoted by cyan colored diamond. **(b)** As determined by ssdtools where the HC05 value is denoted by a dotted line.

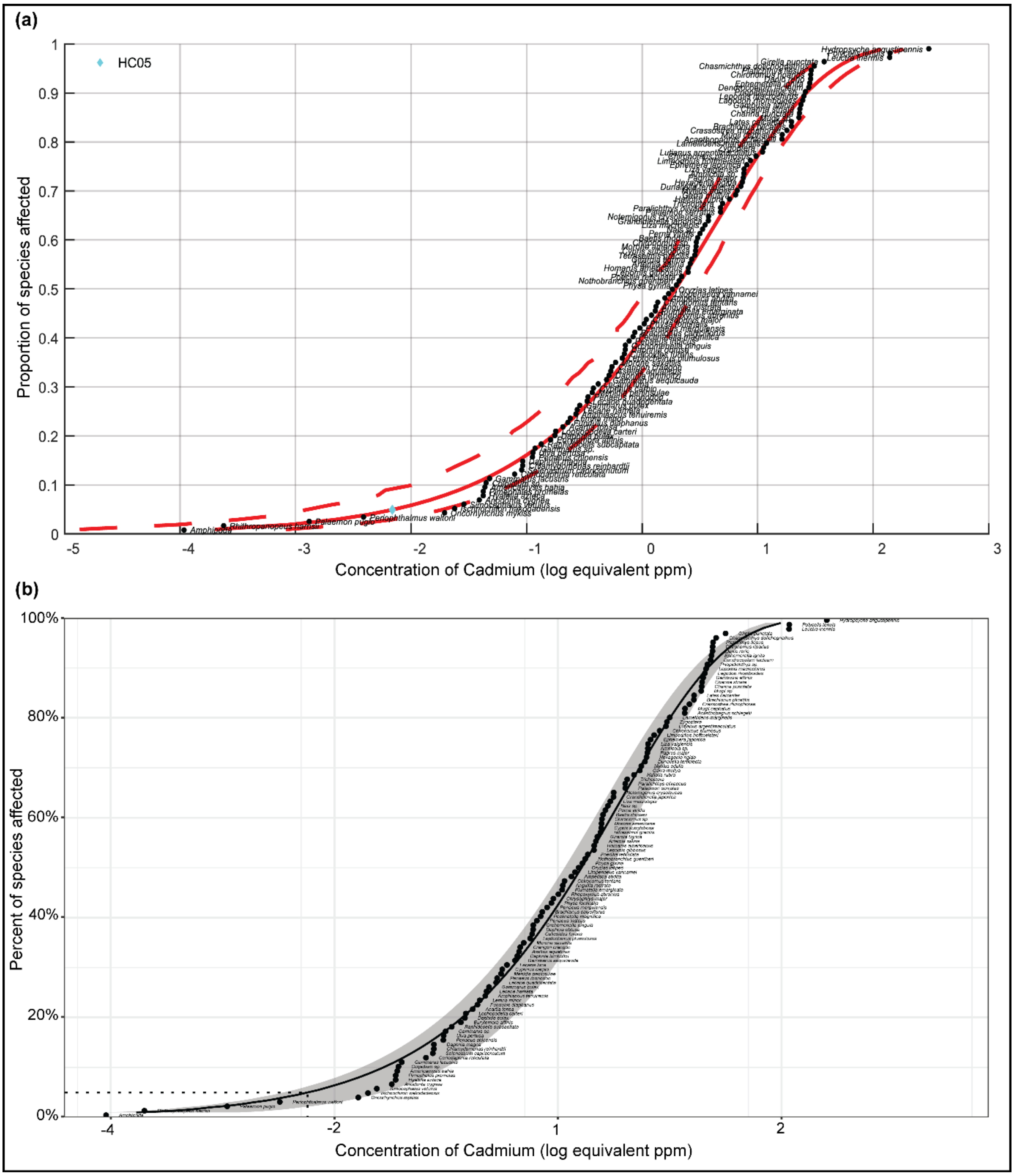


**Figure S12**: The plots of SSD for Cadmium (CAS:7440-43-9) based on acute toxicity data and computed using the best-fit Weibull model. **(a)** As determined by US EPA SSD Toolbox where the HC05 value is denoted by cyan colored diamond. **(b)** As determined by ssdtools where the HC05 value is denoted by a dotted line.

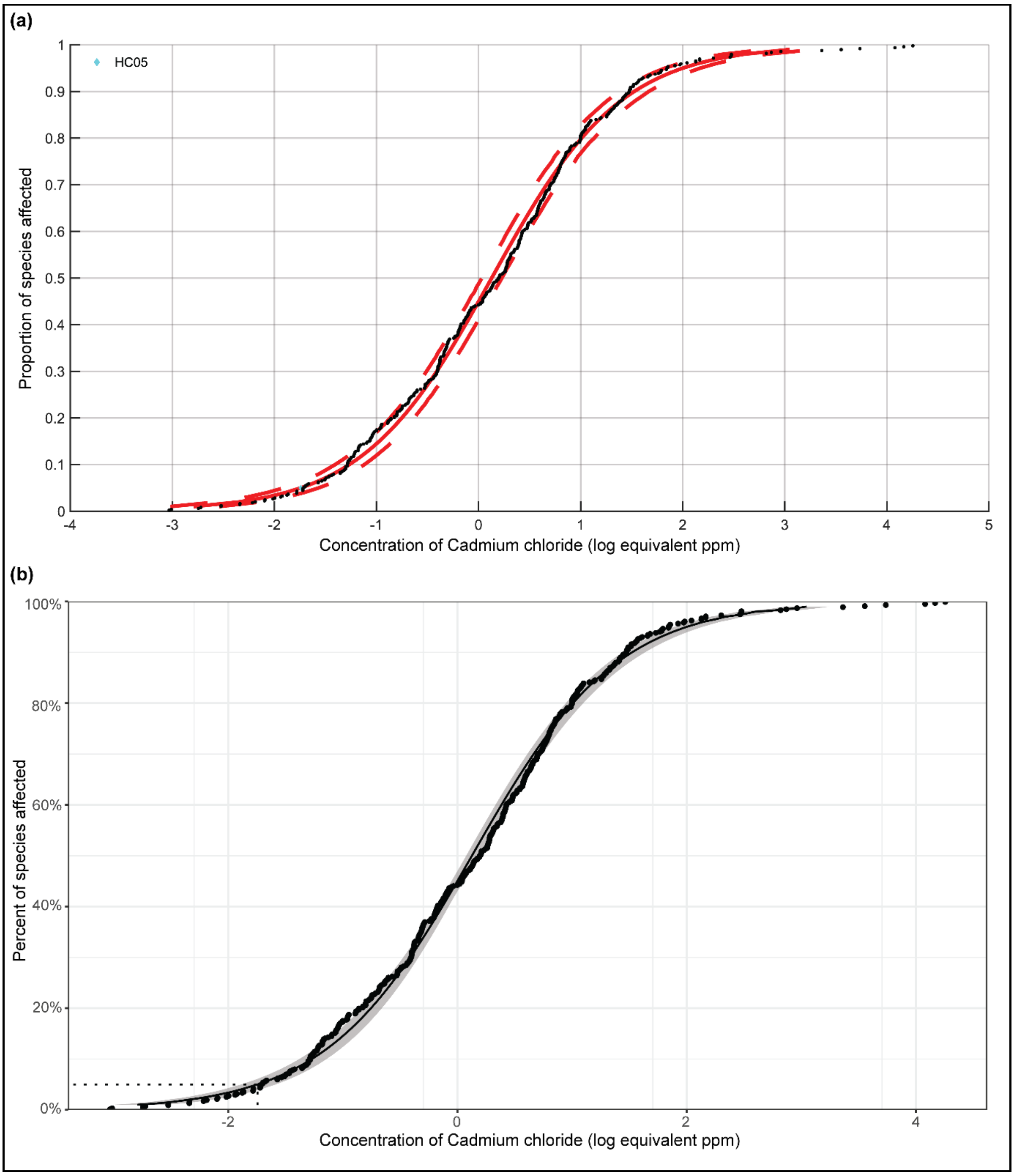


**Figure S13**: The plots of SSD for Cadmium chloride (CAS:10108-64-2) based on acute toxicity data and computed using the best-fit Log-Logistic model. **(a)** As determined by US EPA SSD Toolbox where the HC05 value is denoted by cyan colored diamond. **(b)** As determined by ssdtools where the HC05 value is denoted by a dotted line.

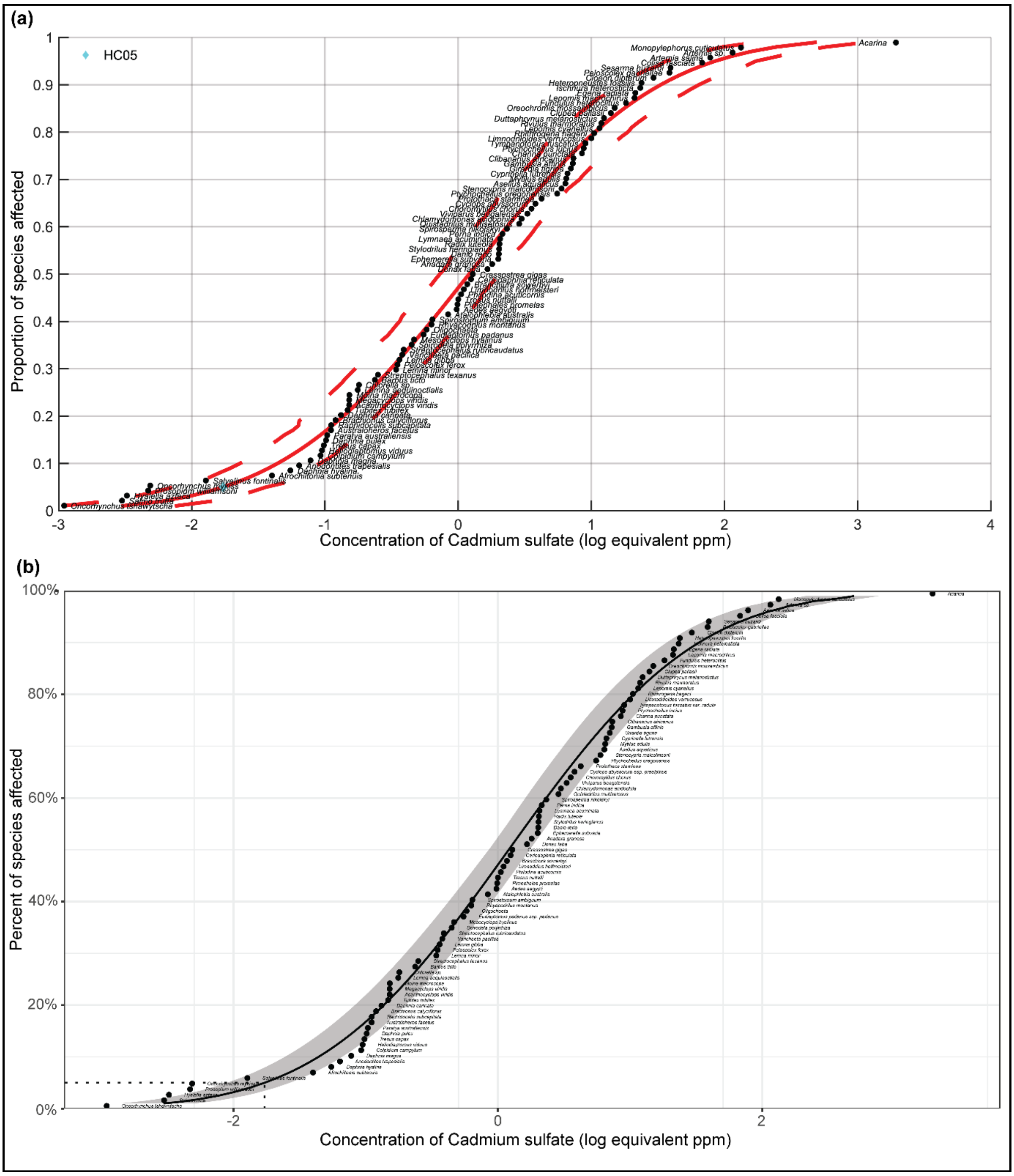


**Figure S14**: The plots of SSD for Cadmium sulfate (CAS:10124-36-4) based on acute toxicity data and computed using the best-fit Log-Normal model. **(a)** As determined by US EPA SSD Toolbox where the HC05 value is denoted by cyan colored diamond. **(b)** As determined by ssdtools where the HC05 value is denoted by a dotted line.

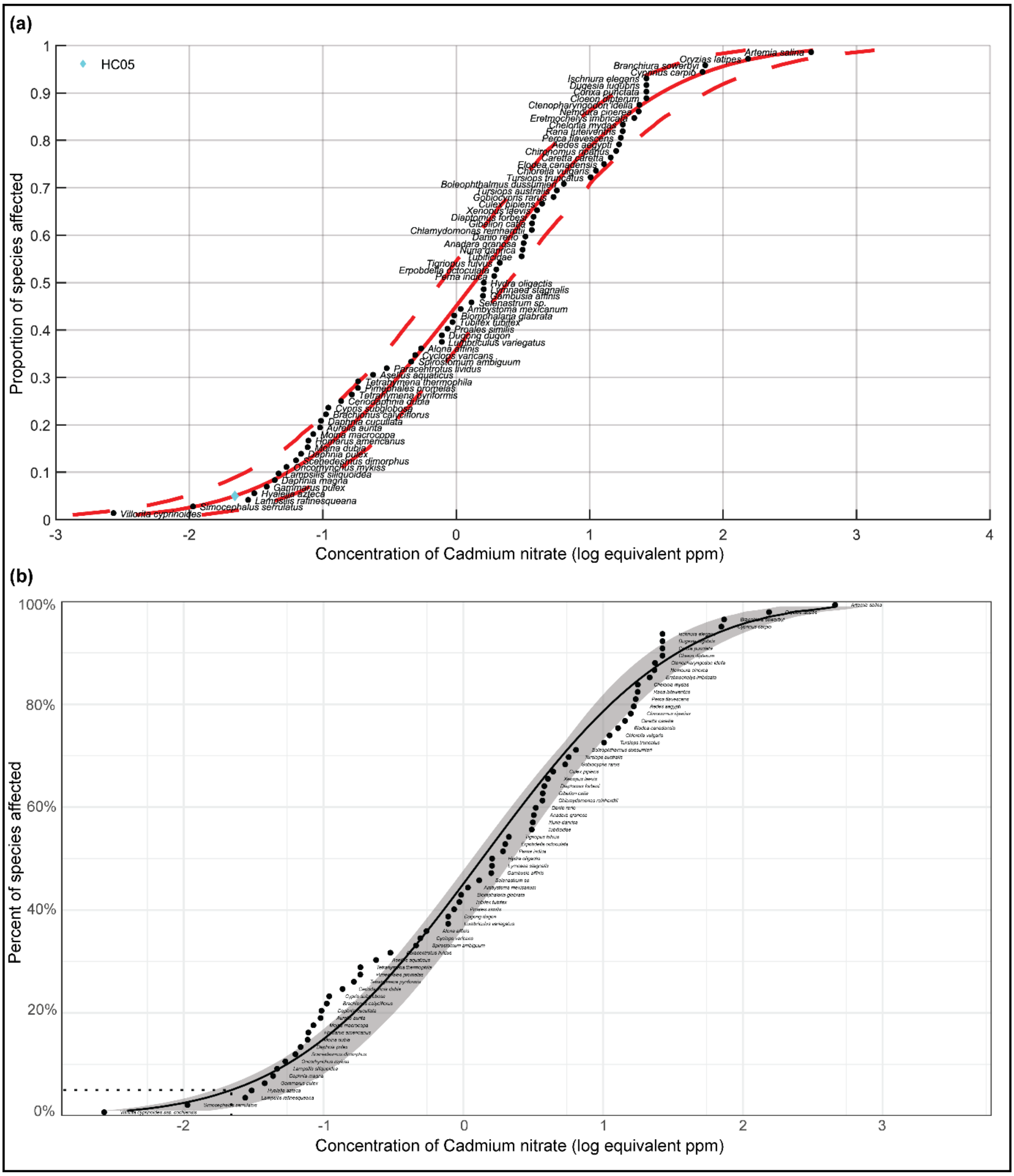


**Figure S15**: The plots of SSD for Cadmium nitrate (CAS:10325-94-7) based on acute toxicity data and computed using the best-fit Weibull model. **(a)** As determined by US EPA SSD Toolbox where the HC05 value is denoted by cyan colored diamond. **(b)** As determined by ssdtools where the HC05 value is denoted by a dotted line.

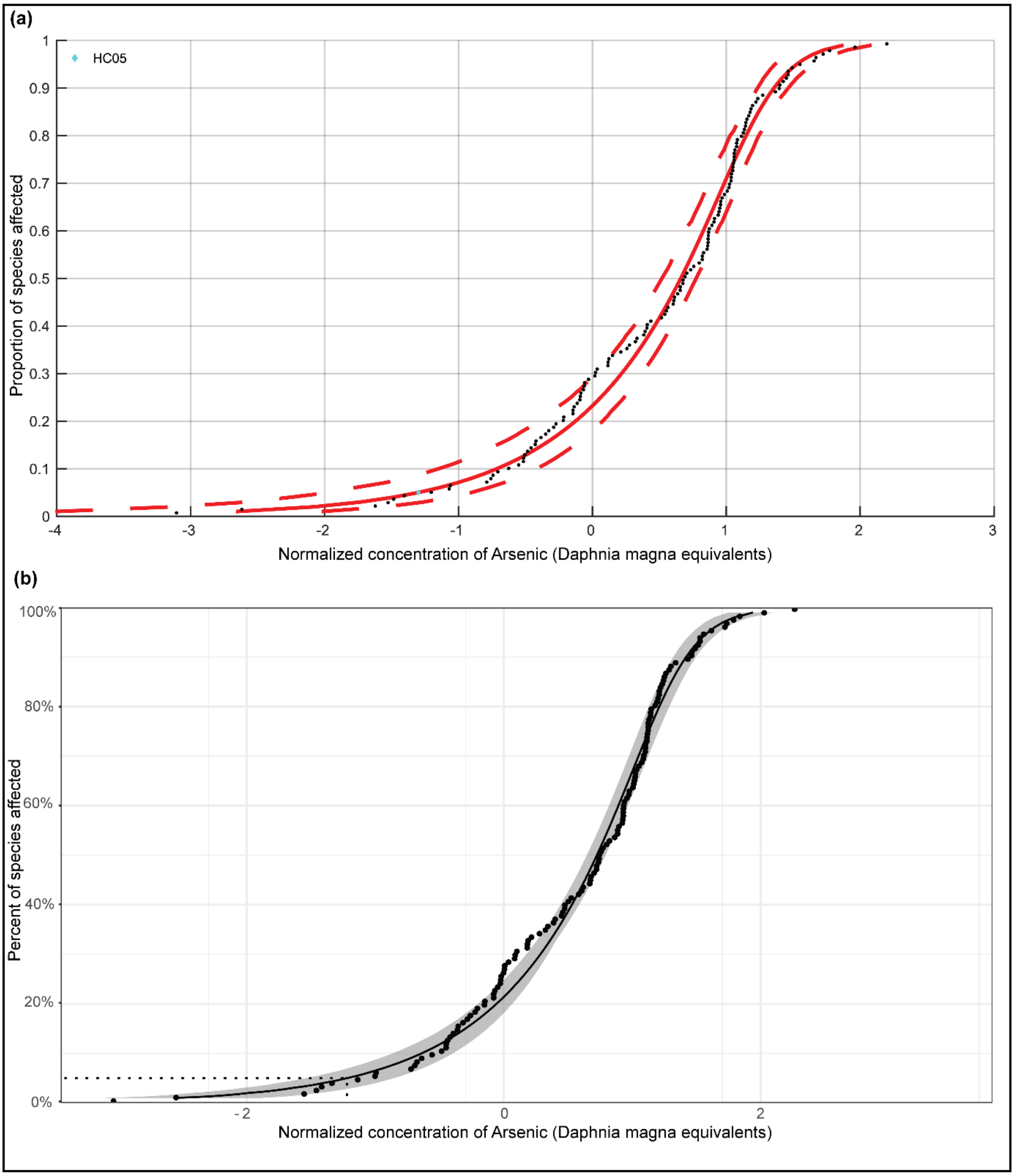


**Figure S16**: The plots of toxicity-normalized SSD (SSDn) for Arsenic based on the acute toxicity data and computed using the best-fit BurrIII model. **(a)** As determined by US EPA SSD Toolbox where the HC05 value is denoted by cyan colored diamond. **(b)** As determined by ssdtools where the HC05 value is denoted by a dotted line.

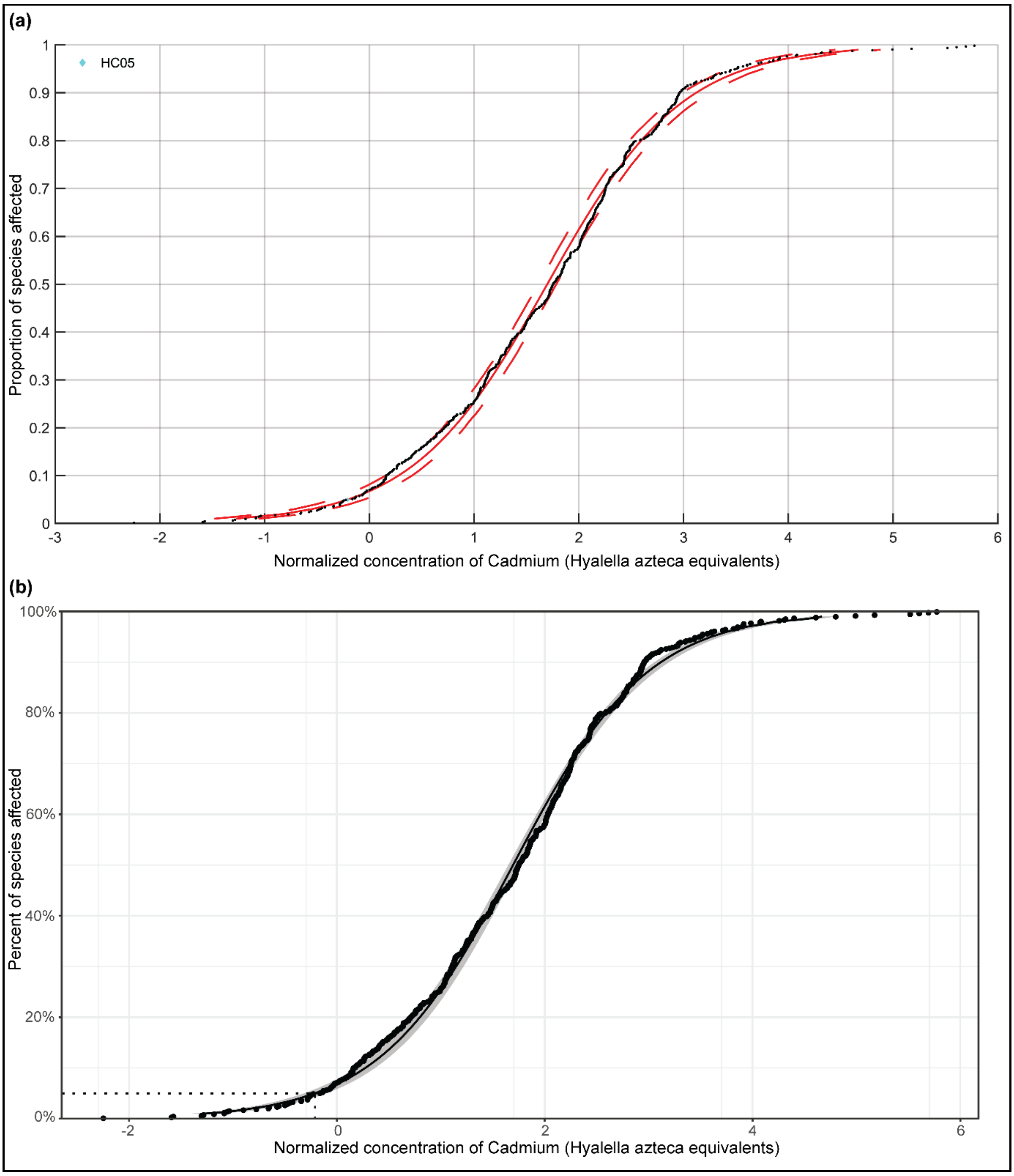


**Figure S17**: The plots of toxicity-normalized SSD (SSDn) for Cadmium based on the acute toxicity data and computed using the best-fit Log-Logistic model. **(a)** As determined by US EPA SSD Toolbox where the HC05 value is denoted by cyan colored diamond. **(b)** As determined by ssdtools where the HC05 value is denoted by a dotted line.

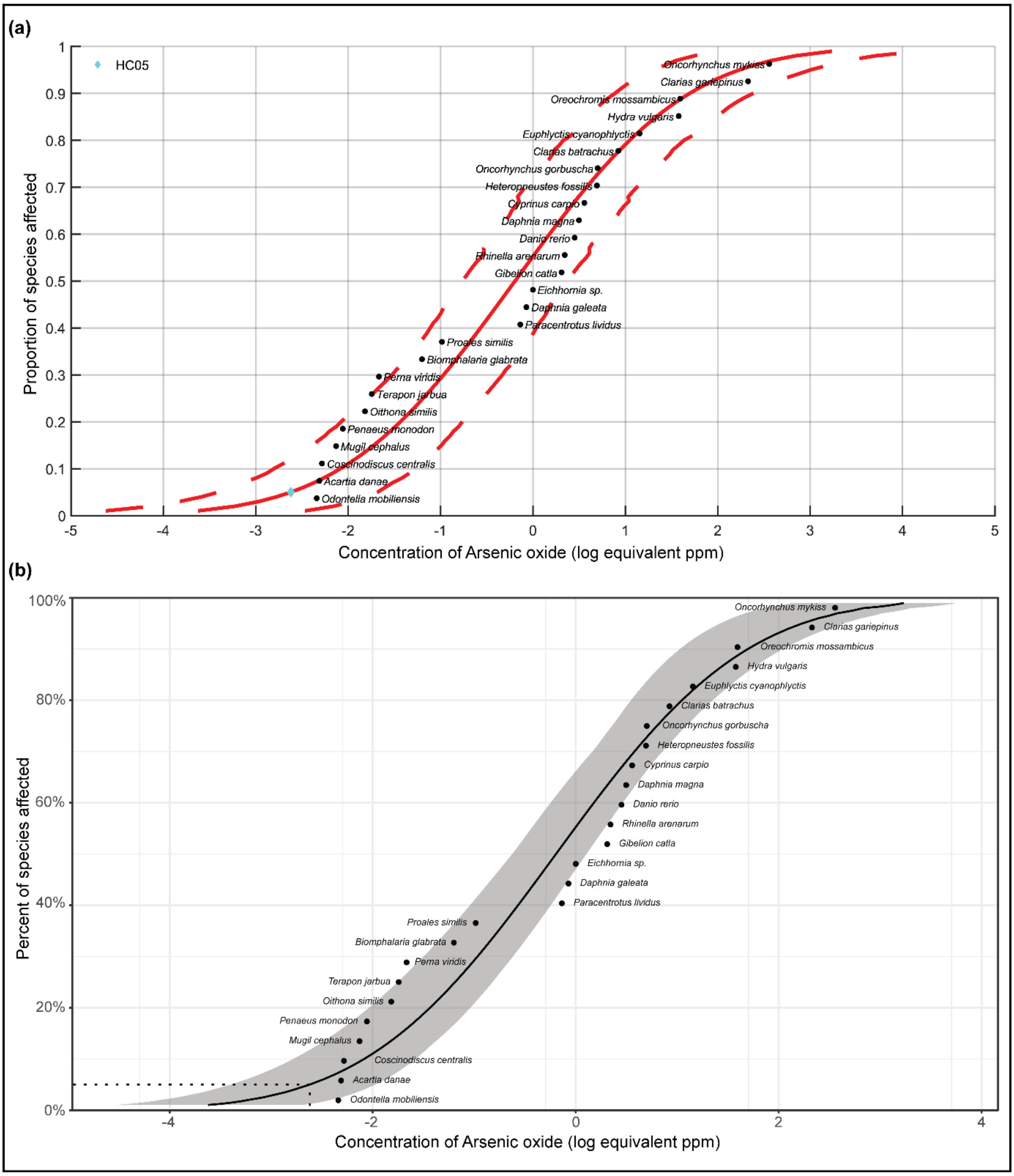


**Figure S18**: The plots of SSD for Arsenic oxide (CAS:1327-53-3) based on chronic toxicity data and computed using the best-fit Log-Normal model. **(a)** As determined by US EPA SSD Toolbox where the HC05 value is denoted by cyan colored diamond. **(b)** As determined by ssdtools where the HC05 value is denoted by a dotted line.

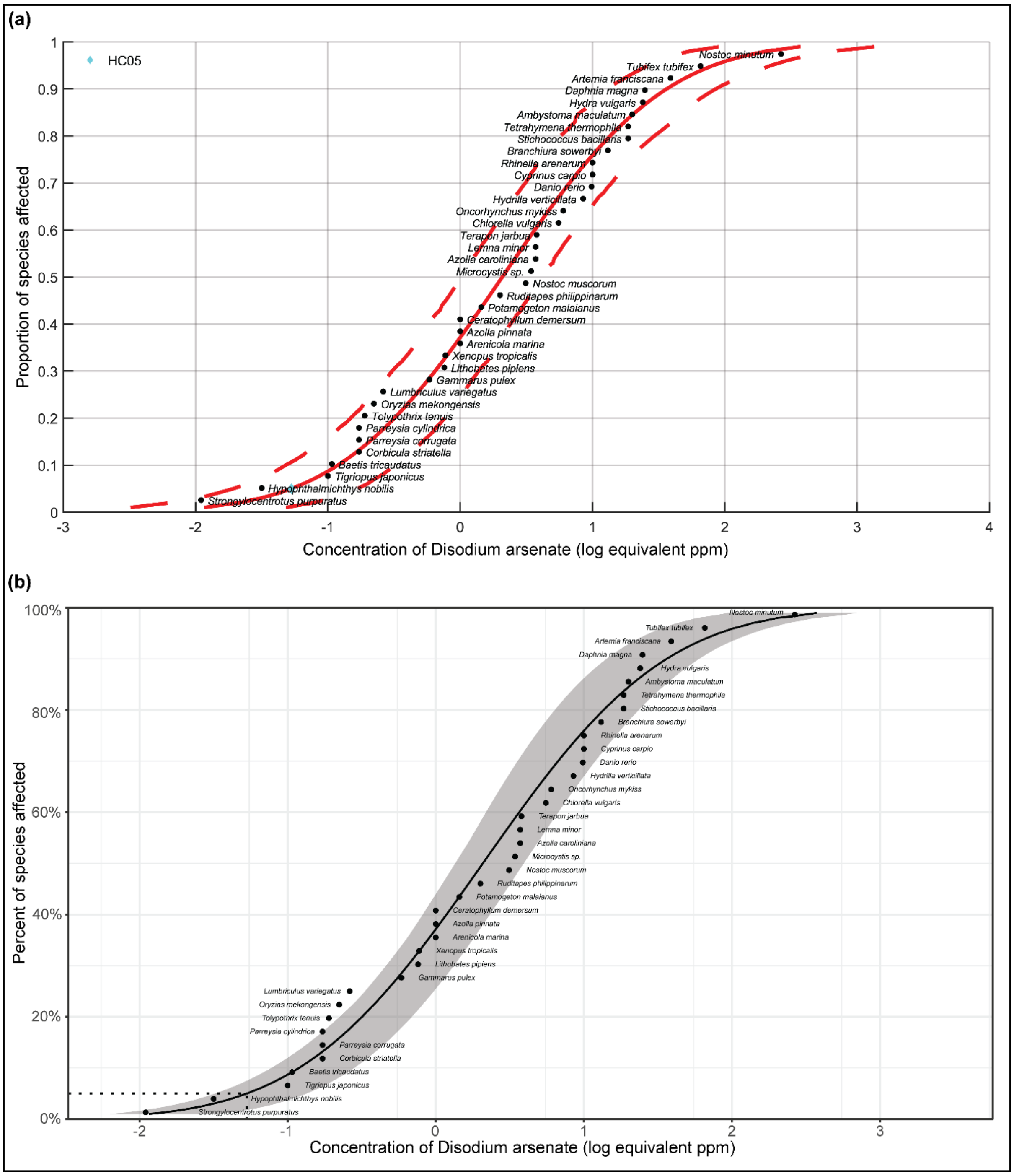


**Figure S19**: The plots of SSD for Disodium arsenate (CAS:7778-43-0) based on chronic toxicity data and computed using the best-fit Log-Normal model. **(a)** As determined by US EPA SSD Toolbox where the HC05 value is denoted by cyan colored diamond. **(b)** As determined by ssdtools where the HC05 value is denoted by a dotted line.

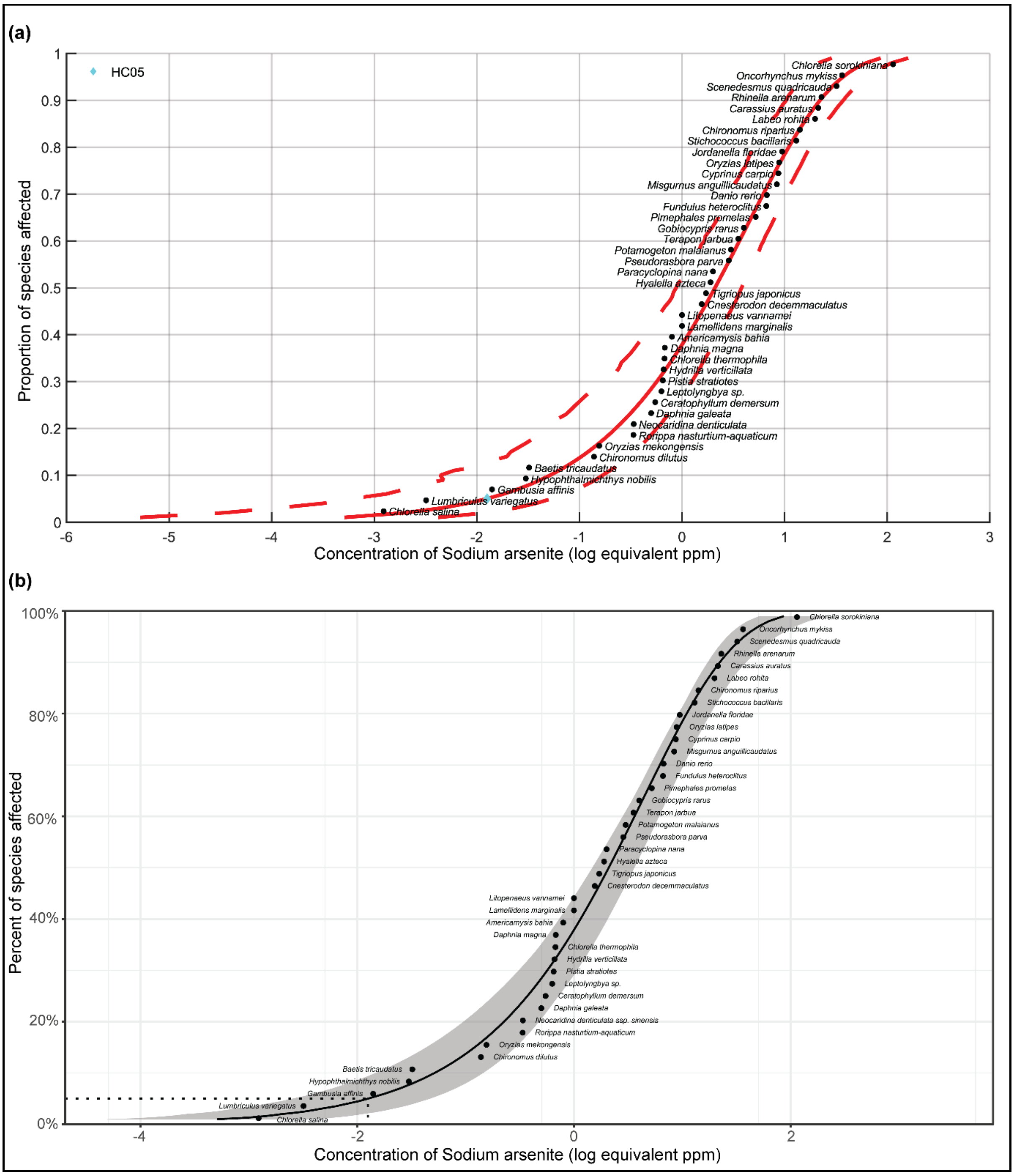


**Figure S20**: The plots of SSD for Sodium arsenite (CAS:7784-46-5) based on chronic toxicity data and computed using the best-fit BurrIII model. **(a)** As determined by US EPA SSD Toolbox where the HC05 value is denoted by cyan colored diamond. **(b)** As determined by ssdtools where the HC05 value is denoted by a dotted line.

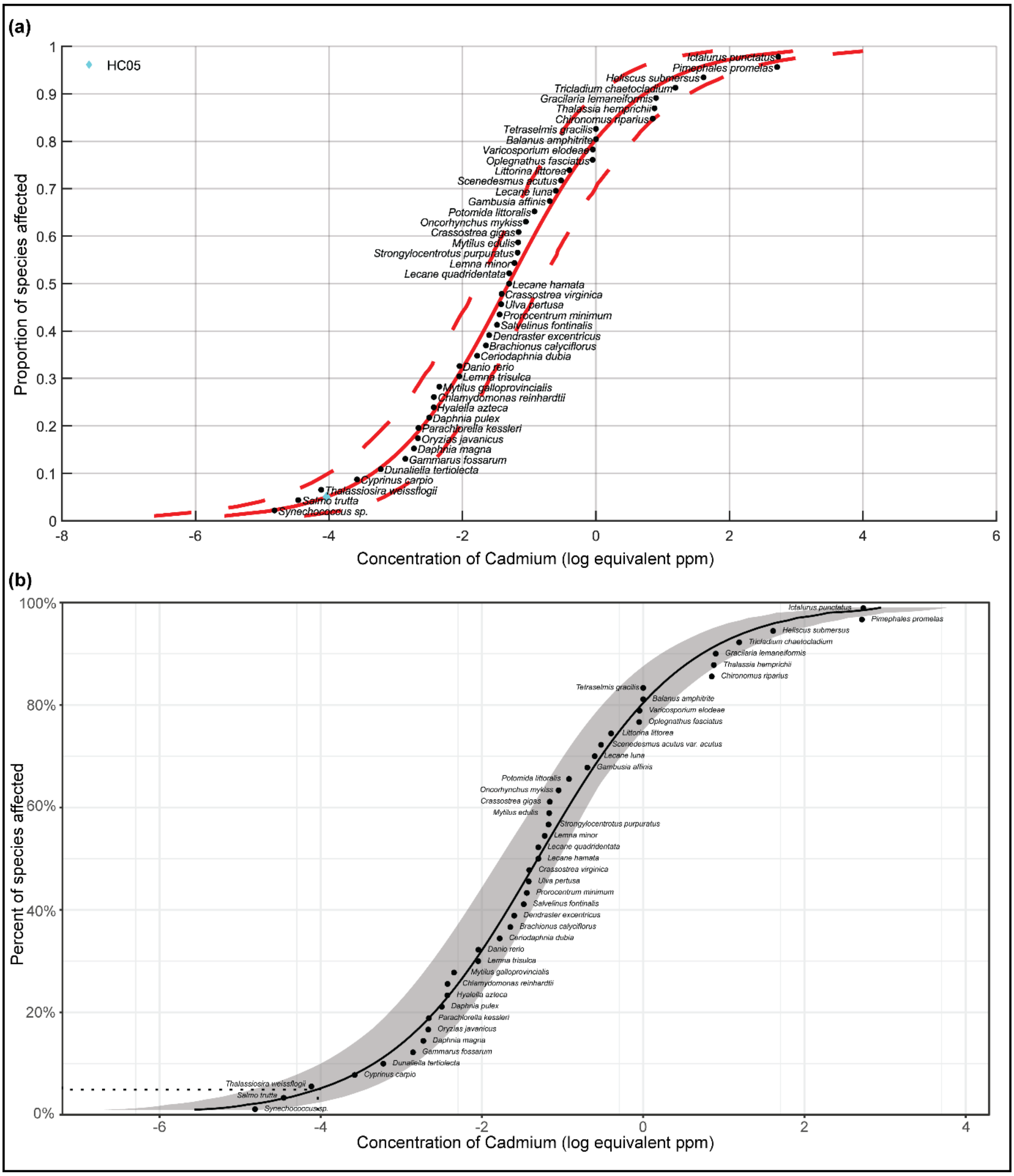


**Figure S21**: The plots of SSD for Cadmium (CAS:7440-43-9) based on chronic toxicity data and computed using the best-fit Log-Logistic model. **(a)** As determined by US EPA SSD Toolbox where the HC05 value is denoted by cyan colored diamond. **(b)** As determined by ssdtools where the HC05 value is denoted by a dotted line.

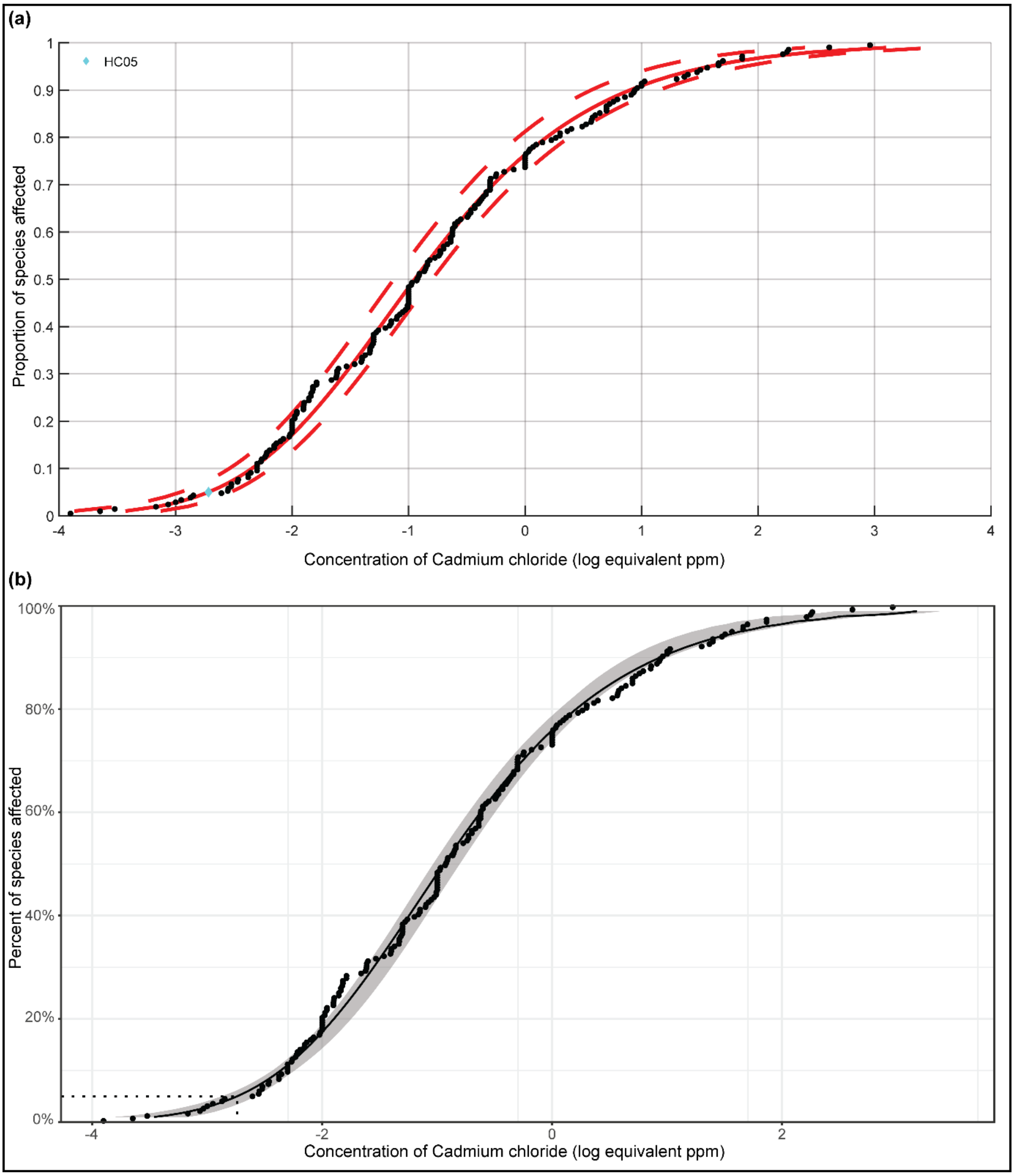


**Figure S22**: The plots of SSD for Cadmium chloride (CAS:10108-64-2) based on chronic toxicity data and computed using the best-fit BurrIII model. **(a)** As determined by US EPA SSD Toolbox where the HC05 value is denoted by cyan colored diamond. **(b)** As determined by ssdtools where the HC05 value is denoted by a dotted line.

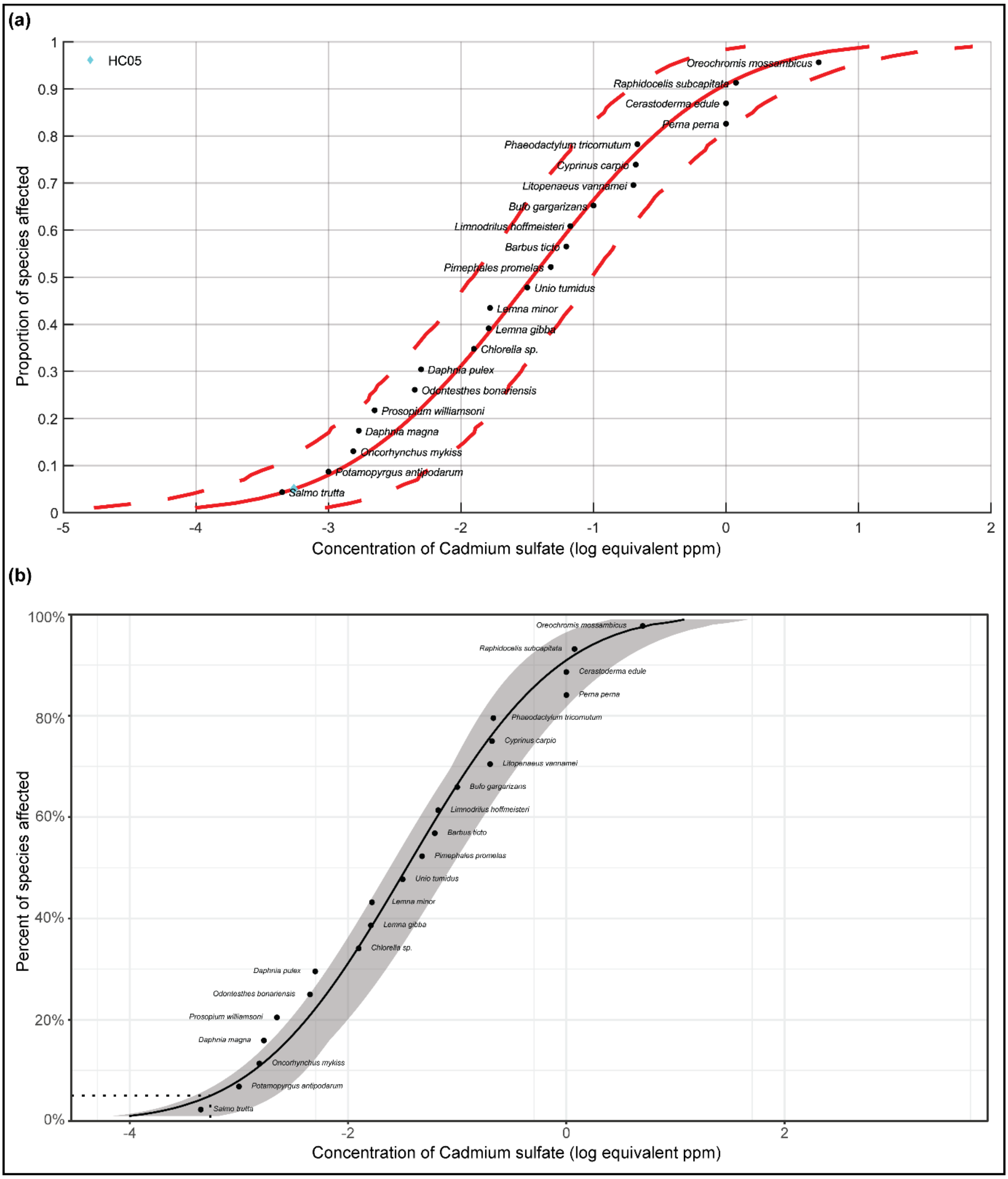


**Figure S23**: The plots of SSD for Cadmium sulfate (CAS:10124-36-4) based on chronic toxicity data and computed using the best-fit Log-Normal model. **(a)** As determined by US EPA SSD Toolbox where the HC05 value is denoted by cyan colored diamond. **(b)** As determined by ssdtools where the HC05 value is denoted by a dotted line.

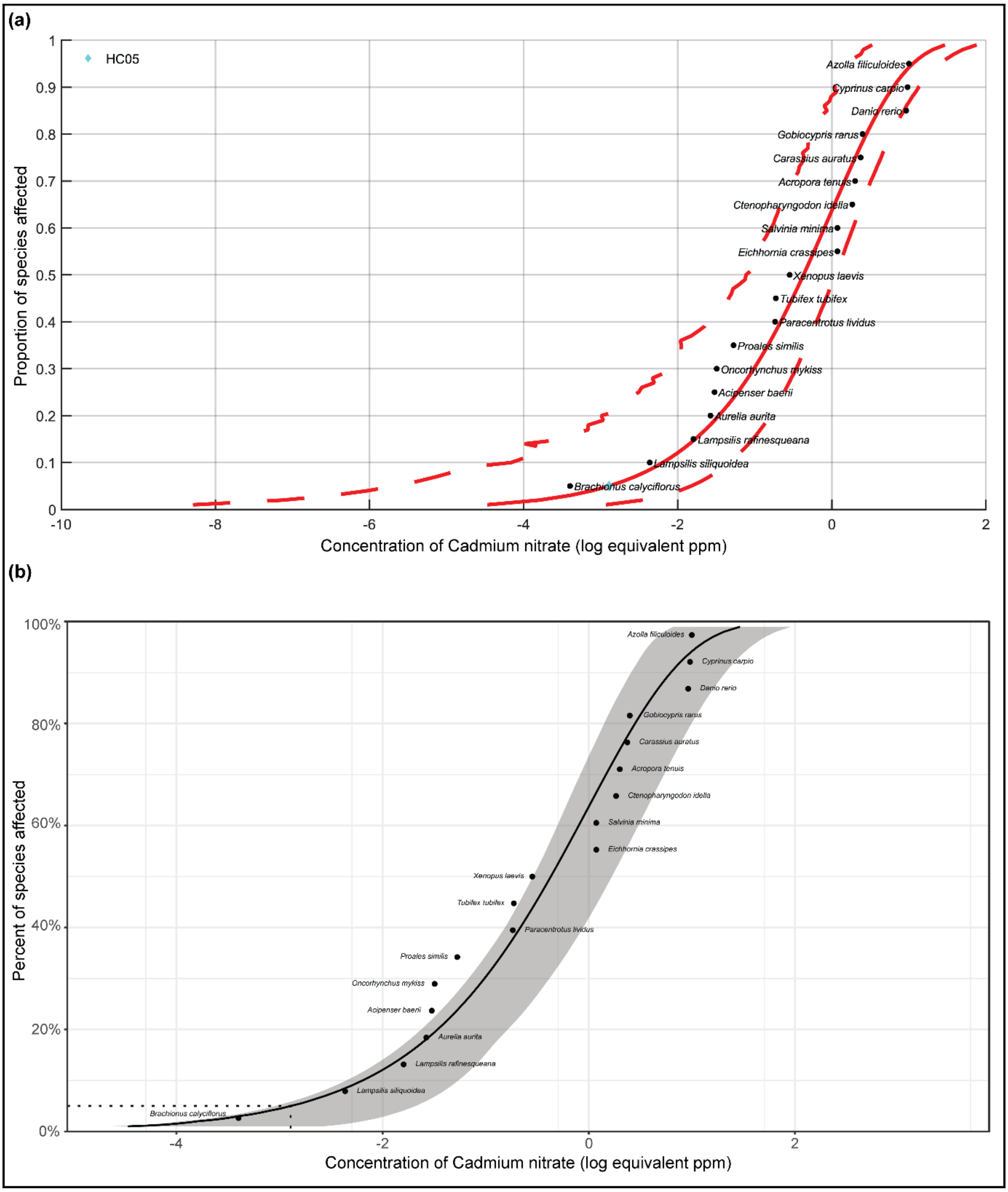


**Figure S24**: The plots of SSD for Cadmium nitrate (CAS:10325-94-7) based on chronic toxicity data and computed using the best-fit Weibull model. **(a)** As determined by US EPA SSD Toolbox where the HC05 value is denoted by cyan colored diamond. **(b)** As determined by ssdtools where the HC05 value is denoted by a dotted line.

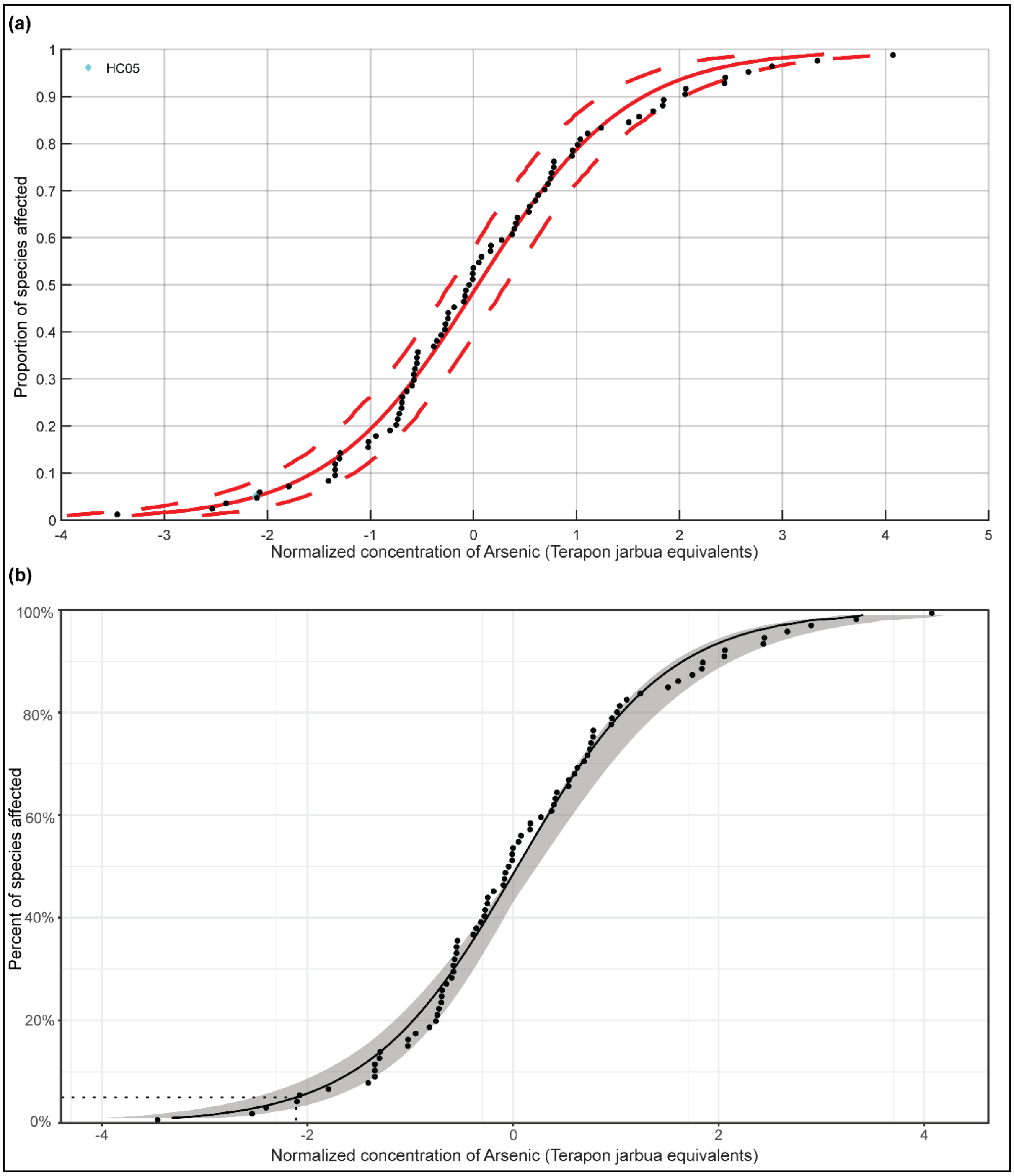


**Figure S25**: The plots of toxicity-normalized SSD (SSDn) for Arsenic based on the chronic toxicity data and computed using the best-fit Log-Logistic model. **(a)** As determined by US EPA SSD Toolbox where the HC05 value is denoted by cyan colored diamond. **(b)** As determined by ssdtools where the HC05 value is denoted by a dotted line.

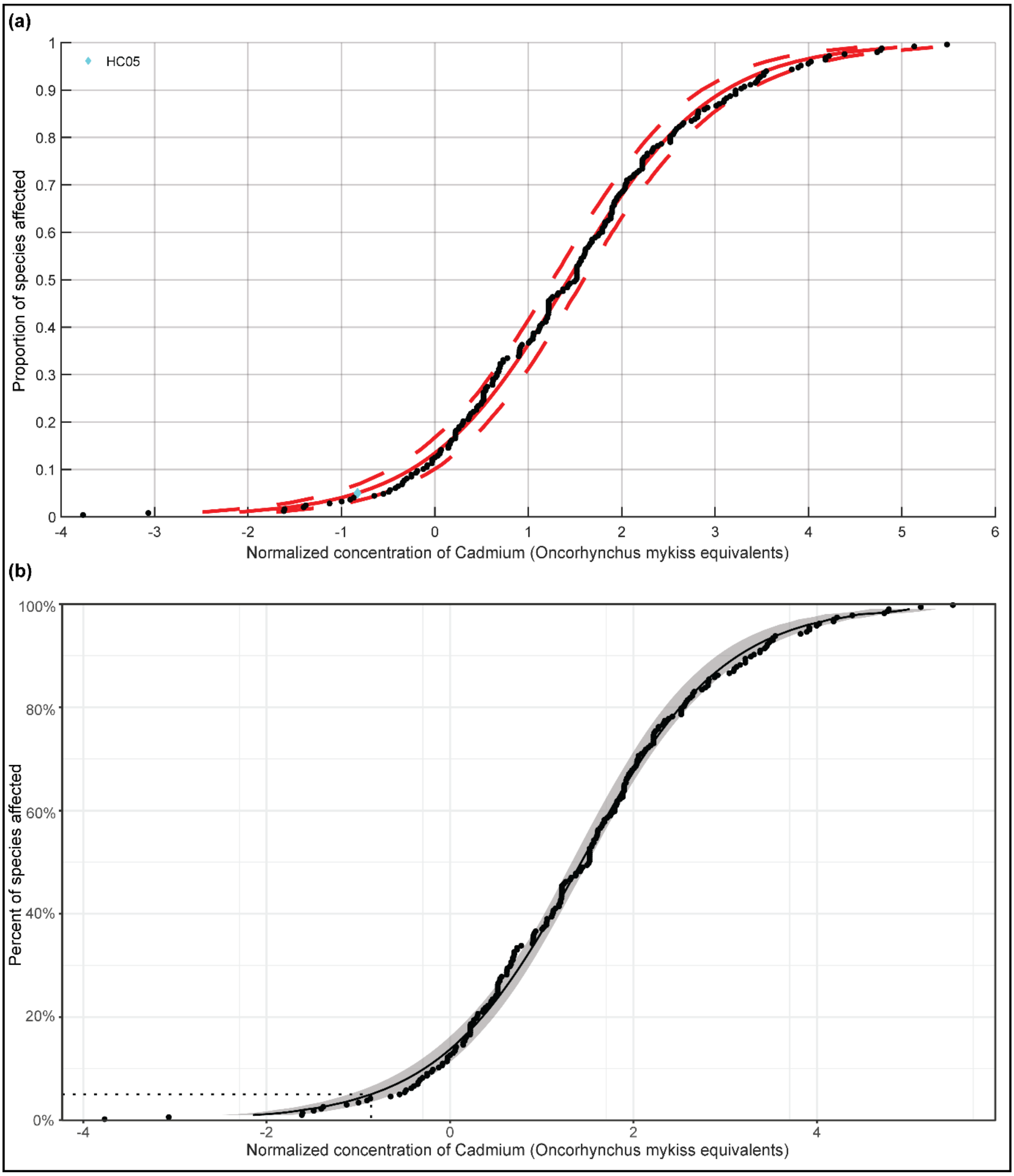


**Figure S26**: The plots of toxicity-normalized SSD (SSDn) for Cadmium based on the chronic toxicity data and computed using the best-fit Log-Logistic model. **(a)** As determined by US EPA SSD Toolbox where the HC05 value is denoted by cyan colored diamond. **(b)** As determined by ssdtools where the HC05 value is denoted by a dotted line.
